# Ophidian physique: the capacity of middle trunk vertebral shape for quantitative taxonomic delimitation in snakes

**DOI:** 10.1101/2023.08.01.551561

**Authors:** John J. Jacisin, A. Michelle Lawing

## Abstract

Skeletal specializations in snakes have resulted in incredible locomotive adaptability, including oft-overlooked vertebral complexity. Snake vertebrae are usually identified via qualitative descriptions of morphological traits; however, identifying and describing snake vertebrae in a scientifically replicable way has long hindered fossil snake research, where attempts to identify between or within snake groups can be onerous. Here, we build a framework of extant snake middle trunk vertebral shape using 2D geometric morphometrics (GMM) and quantitative methods to explore the viability of using these tools to assign and identify snake trunk vertebrae taxonomically and ecologically. We use 23 landmarks to evaluate anterior vertebral shape variation in 504 snake trunk vertebrae representing 189 species across 11 families for delimiting taxonomy and primary foraging habitat. We found that snake vertebral shape variation of overall proportions and articular surfaces contained statistically significant taxonomic and ecomorphological information useful for group assignment. Differences in primary foraging habitat also resulted in similar morphological trends within taxonomic groups in shape space. We then used linear discriminant functions to test the reliability of taxonomic assignments based on the shape captured by our landmark scheme. Analysis of the full dataset had high overall accuracy for family and subfamily, but only moderate success for genus, species, and primary foraging habitat. When applied to a single subfamily, overall accuracy greatly increased for genus and primary foraging habitat, implying that iterative application of this method may improve results. This study presents the framework for a new replicable method to supplement qualitative morphological descriptions of taxa. We recommend that GMM is best employed alongside qualitative descriptions for the optimal and reproducible delimitation of snake vertebrae. Finally, this method will allow non-expert diagnosticians to have more confidence in identifying fossil snake vertebrae, helping increase the number of snake fossils identified in museum collections.

## Introduction

Snakes are amongst the most specialized vertebrates in history, equipped with evolutionary innovations for sensing the environment, feeding, and diverse modes of locomotion. While snakes are not the only limbless vertebrates, the morphology of their vertebrae is more complex and efficient than their limbless vertebrate compatriots (Holman, 2000). The elongate body form, skull specialization, and lack of appendages in snakes combine to result in impressive locomotive adaptability and oft-overlooked morphological complexity in the vertebral column (Johnson, 1955; Savitzky, 1980; Holman, 2000; Lillywhite et al., 2000; Jayne, 2020; Jurestovsky et al., 2020). In fact, at least 11 locomotive gaits for traveling across various substrates have been identified in snakes thus far, with recent research suggesting that rectilinear and sidewinding locomotion are unique gaits, but that lateral undulation and concertina categories can be further divided into five and four distinct motor patterns for specific needs to traverse on, across, or through different mediums (Jayne, 2020). In cases where characters based on color pattern, scale counts, linear measurements, head shape, and molecular data as typically used are impossible, such as in the fossil record or in dry museum specimens, vertebral morphology has been used to implicate both taxonomy and ecology (Gilmore, 1938; Johnson, 1955; Meylan, 1982; Holman, 2000; Lillywhite et al., 2000; Shine et al., 2002; Fabien et al., 2004; Manier, 2004; Vincent et al., 2004; Klein et al., 2021).

Snake trunk vertebrae are the most common snake fossil elements in North America, but diagnosing and interpreting vertebral morphology in both living and fossil snakes based on apomorphies is limited to relatively few expert diagnosticians (Rage, 1984; Holman, 2000; Bell et al., 2010; Parmley and Hunter, 2010; Smith, 2013). The difficulty in these endeavors lies in discerning minute but complex differences in overall shape, proportion, and the effects of an exceptionally elaborate system of axial muscles and vertebral articulations, which substitute for the lack of appendicular appendages and allow for extreme locomotive behaviors (e.g., cantilevering, tying their bodies in knots, and spinning; Cundall, 1987; Holman, 2000; Jayne, 2020). As a result of this difficulty, fossil snakes (much like other fossil squamates) are not well studied in comparison with coeval mammalian counterparts despite samples being available in many localities. Detecting identifiable morphological features or patterns in vertebrae inherent to groups of snakes is useful for describing and assigning taxa and ecological relationships through time.

Snake vertebrae are usually identified via qualitative descriptions of morphological traits (e.g., Auffenberg, 1963; Szyndlar, 1984; LaDuke, 1991; Holman, 2000) that are occasionally supplemented by quantified linear measurements and ratios (e.g., Johnson, 1955; Meylan, 1982; Jasinski and Moscato, 2017; Klein et al., 2021). Unfortunately, the dearth of spectacular specimens and the difficulty of identifying and describing snake elements in a scientifically replicable way has long hindered progress in research of the snake skeleton and in snake fossils. Attempts to identify between (e.g., Coluber and Masticophis) or within (e.g., crotalines) some snake groups at various taxonomic levels are often onerous and hindered by a lack of comparative methods and a poor understanding of snake osteology (Holman, 2000). In particularly complex cases, new, easily applicable methods are necessary to improve the processes of description, differentiation, and identification of snake skeletal material for both expert and inexperienced diagnosticians. Geometric morphometrics is one such methodology with excellent potential to assist in the process of assigning snake vertebrae to taxon and ecology. The implementation of geometric morphometrics would therefore provide a wealth of previously unexplored data on snake fossils and the snake skeleton.

Isolated snake vertebral elements are historically understudied, overlooked, or ignored completely in samples and museum collections (Holman, 2000), inhibiting the accurate representation of snake richness and functional diversity in many taxonomic and paleoecological interpretations. Geometric morphometrics present an alternative, shape-based, and quantitative approach for comparing and depicting snake vertebrae. While these methods have not been widely applied to snake vertebrae, or even to snakes in general (e.g., Gentilli et al., 2009; Lawing and Polly, 2011; Lawing et al., 2012; Ruane, 2015; Huntley et al., 2021), some studies have shown that geometric morphometrics can detect morphological differences that support molecular taxonomic hypotheses at clade and even subspecies levels (Gentilli et al., 2009; Lawing and Polly, 2011; Meik et al., 2012; Jacisin, 2021). The ability of geometric morphometric techniques to work on isolated elements such as snake trunk vertebrae would additionally provide a path forward to shape-based studies relating morphology to functional trait-environment relationships at individual to community levels, as in ecometrics, as well as skeletal studies of sexual dimorphism, allometry, and inter- or intraspecies variation.

Here, we evaluate the separation of anterior trunk vertebral shape among taxonomic categories using geometric morphometrics and additional multivariate analyses as a method of specimen assignment and shape differentiation. This methodology is applied across a vast array of extant snake groups and a large number of individuals. In doing so, we seek to create a methodological framework using extant snake taxa to answer the following questions: 1) can shape detected via geometric morphometrics act as a supplement to visual descriptions for assigning, identifying and describing taxa and their ecologies; 2) what shape changes explain the most variation in the anterior aspect of snake trunk vertebrae; and 3) how well do the observed differences in shape associate with taxa at the family, subfamily, genus, and species levels? If there are differences in shape related to phylogeny,we can expect greater morphological differences, and therefore increased predictive capacity, at higher levels of the taxonomic hierarchy. Furthermore, if these methods are capable of taxonomic delimitation, then two-dimensional geometric morphometrics stands as a relatively simple – but powerful – tool for identifying both extant and fossil material based solely on vertebral shape, while requiring less anatomical expertise than more traditional methods.

## Methods

We used geometric morphometrics to explore the viability of quantitative shape analysis as a method for assigning recent and fossil snake trunk vertebrae to already described and delimited species. We photographed the anterior view of 504 extant snake middle trunk vertebrae, ranging from one to three vertebrae per individual, one to ten individuals per species, and one to fifteen species per genus. Photographs were taken of material from the Texas A&M University Biodiversity Research and Teaching Collections (TCWC), the Texas Vertebrate Paleontology Collections (TMM), the University of Texas at Arlington (UTA), the William R. Adams Zooarcheology Lab, Indiana University (WRA), the University of Nebraska State Museum (UNSM), and the Smithsonian National Museum of Natural History (USNM). We used a DinoLite Edge digital microscope or a Canon EOS Rebel SX camera with a Canon Zoom EF-S lens (18-55 mm) for all photographs. The vertebrae comprised 11 families, 16 subfamilies, 89 genera, and 189 species from around the world, but primarily from North America (62%), based on availability of museum skeletal material with isolated trunk vertebrae. Each isolated vertebra was secured such that the anterior surface of the vertebra was as planar as possible, perpendicular and centered below the camera lens in order to minimize distortion from the curvature of the camera lens as recommended by Head et al. (2005). We used TPSUTIL, version 1.74 to build a TPS file from the photographs. We also ordered a Microsoft Excel file with the classification information for each specimen for use in RStudio.

We described the shape for the anterior view of snake middle trunk vertebrae by digitizing 23 homologous landmarks in the TPSDIG2, version 2.30 application. This landmark scheme is based on the landmark scheme of Lawing et al. (2012), as We landmarked both lateral sides of the anterior vertebral aspect. All landmarks selected are homologous, clear on all specimens used, and best represent the anterior aspect of the vertebra (Fig. 1; Table 1). We selected the anterior aspect because it contains the most information on overall shape and articular surfaces on a single plane, and is therefore most ideal for capturing variation related to function and locomotive behaviors in two dimensions (Lawing et al., 2012).

**Figure 1.**
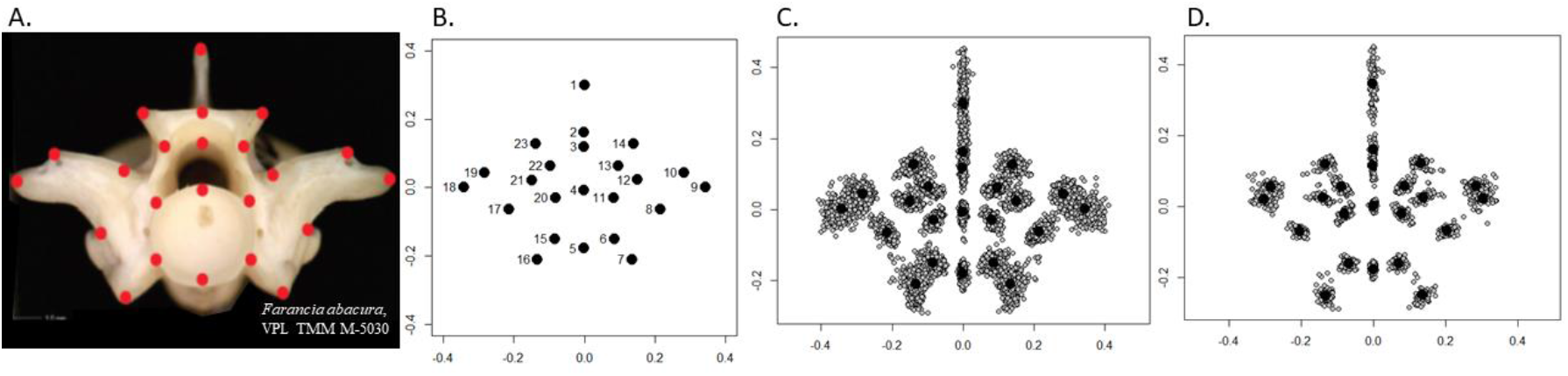
Overall landmark distributions of the data after implementing Procrustes superimposition. A: a photograph of the anterior aspect of a middle trunk vertebra with landmarks placed; B: the landmarks numbered in order of placement for each vertebra; C: Procrustes superimposed landmarks for vertebrae among all snake families in this study; and D: Procrustes superimposed landmarks for vertebrae for the Crotalinae-only dataset. Centroid values of each landmark are represented by large black circles.

**Table 1.** List of 23 vertebral landmarks used for anterior shape in this study.

| Landmark | Description |
| --- | --- |
| 1 | Dorsal edge of neural spine |
| 2 | Midline dorsal edge of neural arch |
| 3 | Midline ventral edge of neural arch |
| 4 | Midline dorsal edge of centrum |
| 5 | Midline ventral edge of centrum |
| 6 | Right ventrolateral contact between apophysis & centrum |
| 7 | Right ventromedial edge of parapophysis |
| 8 | Right dorsomedial edge of diapophysis |
| 9 | Right lateral edge of prezygapophyseal body |
| 10 | Right dorsolateral edge of prezygapophyseal articular facet |
| 11 | Right medial contact between centrum & neural arch |
| 12 | Right ventromedial edge of prezygapophyseal articular facet |
| 13 | Right ventrolateral contact between zygosphenes & neural canal |
| 14 | Right dorsolateral edge of zygosphenes |
| 15 | Left ventrolateral contact between apophysis & centrum |
| 16 | Left ventromedial edge of parapophysis |
| 17 | Left dorsomedial edge of diapophysis |
| 18 | Left lateral edge of prezygapophyseal body |
| 19 | Left dorsolateral edge of prezygapophyseal articular facet |
| 20 | Left medial contact between centrum & neural arch |
| 21 | Left ventromedial edge of prezygapophyseal articular facet |
| 22 | Left ventrolateral contact between zygosphenes & neural canal |
| 23 | Left dorsolateral edge of zygosphenes |

To remove shape differences related to absolute size, orientation, and location, we used Procrustes superimposed landmarks (Zelditch et al., 2004; Lawing and Polly, 2010). Elimination of non-shape variation through superimposition differentiates geometric morphometrics from traditional morphometric methods, which are linear and capture size information better than shape information (Rohlf and Marcas, 1993). The overall arrangement of landmarks across all specimens and the centroids for each landmark can be viewed in Fig. 1. We then produced outlier plots for each of the 23 landmarks to ensure that the landmarks were properly ordered across all specimens in the data, as well as to find any outliers potentially indicative of a misdiagnosis a vertebra as belonging to the middle trunk region (Appendix Fig. 1). The proper order of landmarks across all specimens was ensured and outliers were examined and addressed as necessary.

We performed a principal component analysis (PCA) on the middle trunk vertebral shape data using the *geomorph* package (Adams and Otarola-Castillo, 2013) in Rstudio (www.rstudio.com/products/rstudio/download2/) to examine the morphological space of the specimens and quantify independent differences in vertebral shape represented by the PC axes. The maximum variance within the dataset as determined by the PCA is effectively the linear combination of variables after the landmark arrangements are fitted around a mean shape constellation through Procrustes superimposition (Rohlf and Slice, 1990). We also took a subset of the data within a single subfamily, the Crotalinae, and ran a second PCA to investigate differences in the morphological changes representing the most variation within different taxonomic levels. We selected the Crotalinae because it had the best sample size at the subfamily level in the dataset.

In order to graphically illustrate and visually examine the variation in shape for each PC axis, we produced vector plots of the shape variation along each PC axis for both the all-group analysis and the Crotalinae-only analysis. These vector plots show the direction and degree of change at each individual landmark from three negative standard deviations to three positive standard deviations along a given axis. We chose the first six PCs for further analysis based on the results of the PCA, as the first six PCs combined to explain >80% of variance in both datasets.

To assess morphological differences between taxonomic groups, we used multivariate analyses of variance (MANOVAs) on the PCs of shape change for vertebrae at each taxonomic level, first for all groups, then for the Crotalinae-only group. The results of these analyses revealed which axes of shape change represented systematic (non-random) factors useful for assigning taxonomic identities. We also administered a post hoc Tukey’s test in order to determine which specific groups comprised most of the variation at the family and subfamily levels. Finally, to determine whether already delimited species have shape differences that can be used to assign extant and fossil taxa, we used discriminant function analyses (DFAs) at the family, subfamily, genus, and species levels for the all-group dataset and at the genus and species levels for the Crotalinae-only dataset. The DFA classified all individuals included in the study into different groups a priori based on the combined morphological characters represented by the PC axes of shape variation. Application of DFA to both the all-group and the Crotalinae-only dataset allowed us to investigate if the discriminatory ability of lower taxonomic levels improves when using the shape variation present within a single, smaller group rather than across all groups. DFA has been shown to be more robust in the accuracy of its classifications and is relatively unaffected in its values with larger sample sizes when compared to methods such as cluster analysis (Jaiswara et al., 2013). We excluded all groups with a sample size of one (n = 1) from these analyses.

## Results

### Shape variation among snake families

Most of the shape variation was contained in the first few dimensions of the 46 principal components (PC). Specifically, the first six orthogonal PCs accounted for 84.5% of all observed variation in anterior vertebral shape captured by the landmark scheme (Figs. 2, 3; Appendix Fig. 2). Vector plots depicting shape change along each of the first six PC axes in the all-group analysis revealed that most shape variation was explained by differences in seven different locations: the neural spine, the zygosphene, the neural canal, the cotyl, the parapophyses, the diapophyses, and the prezygapophyses (with associated articular facets) (Fig. 4). These seven locations are all associated with articulations in the vertebrae and the overall proportions of height and width, and provide important information on taxonomic and ecological differences. PC1 accounted for 39.8% of the overall variation, and describes shape differences involving the neural spine and parapophyses, and therefore, the relative dorso-ventral height of the vertebra (Fig. 4A). Shapes ranged from proportionally tall vertebrae with long neural spines and parapophyses, but relatively narrower prezygapophyses in the terrestrial viperid *Sistrurus tergeminus* to the dorso-ventrally compressed, relatively wider vertebrae of the fossorial leptotyphlopid *Rena humilis*.

**Figure 2.**
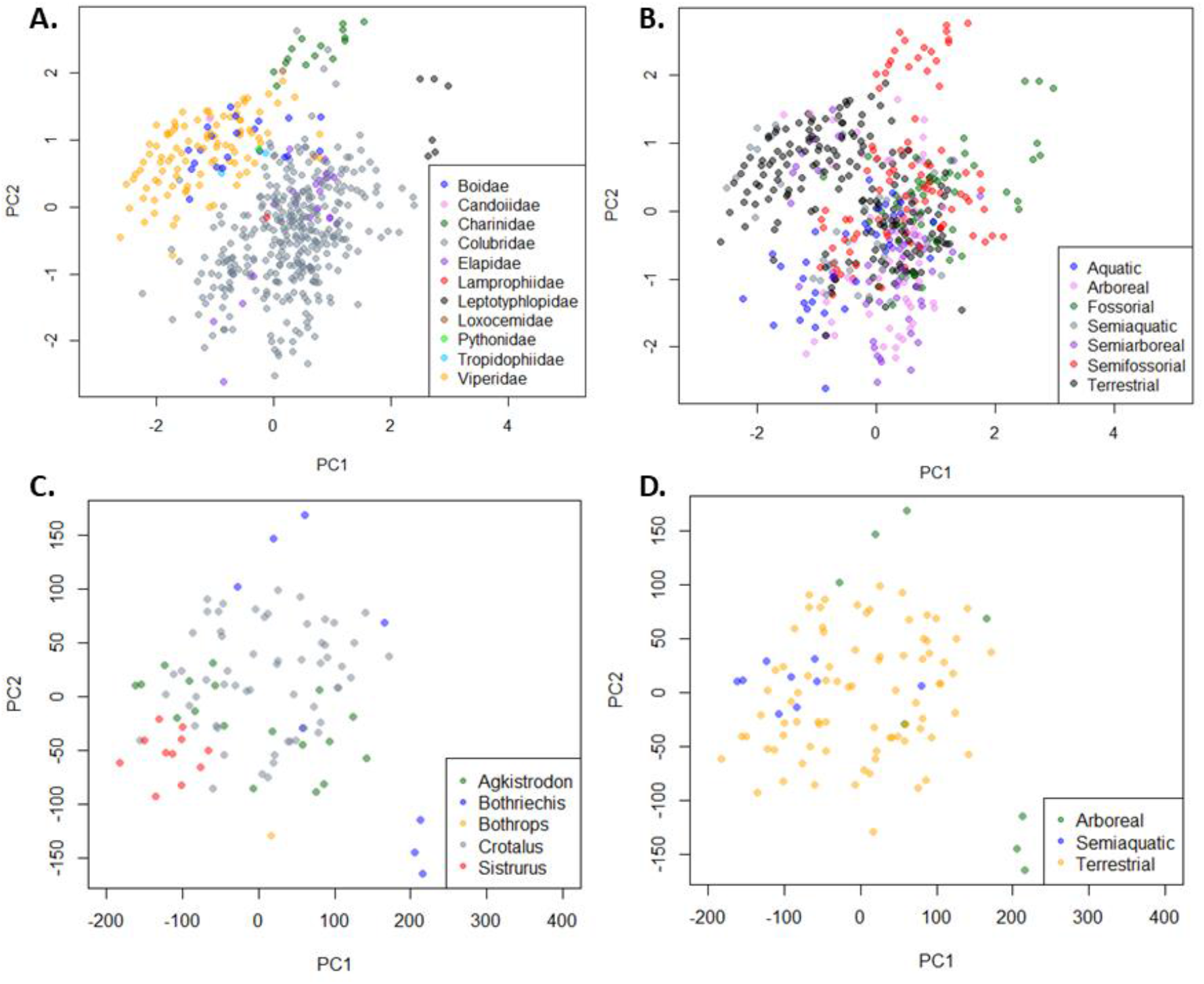
PCA plots of PC1 (x-axis) and PC2 (y-axis) for the all-group (A, B) and Crotalinae (C, D) analyses. Plots A and C are colored by family and genus, respectively, while plots B and D are colored by primary foraging habitat.

**Figure 3.**
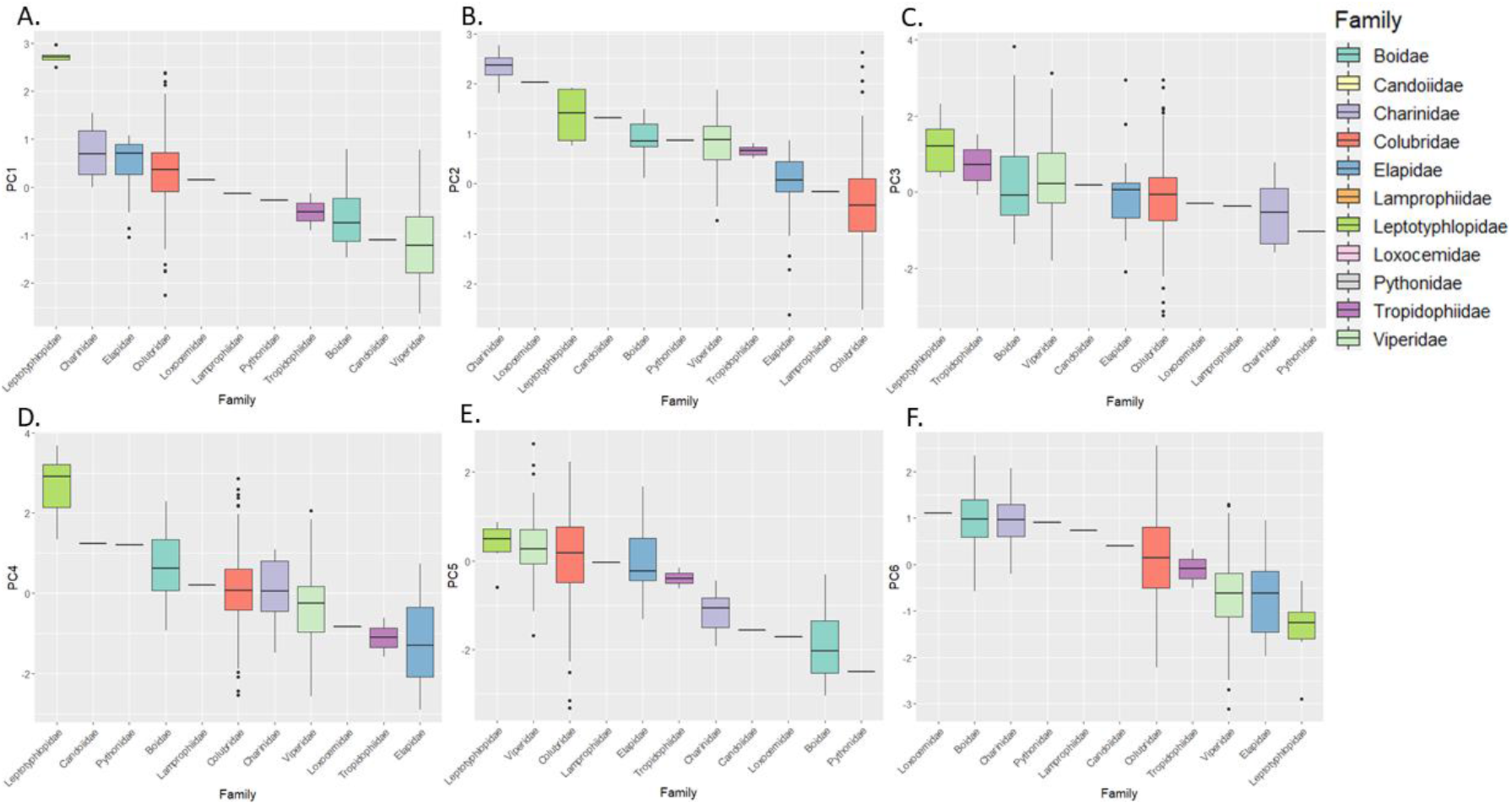
Box and whisker plots of PCs 1-6 in the all-group analysis by family. Colors represent family association. Lines within boxes represent median values, black circles are outliers. Single black lines without boxes are sample sizes of n=1.

**Figure 4.**
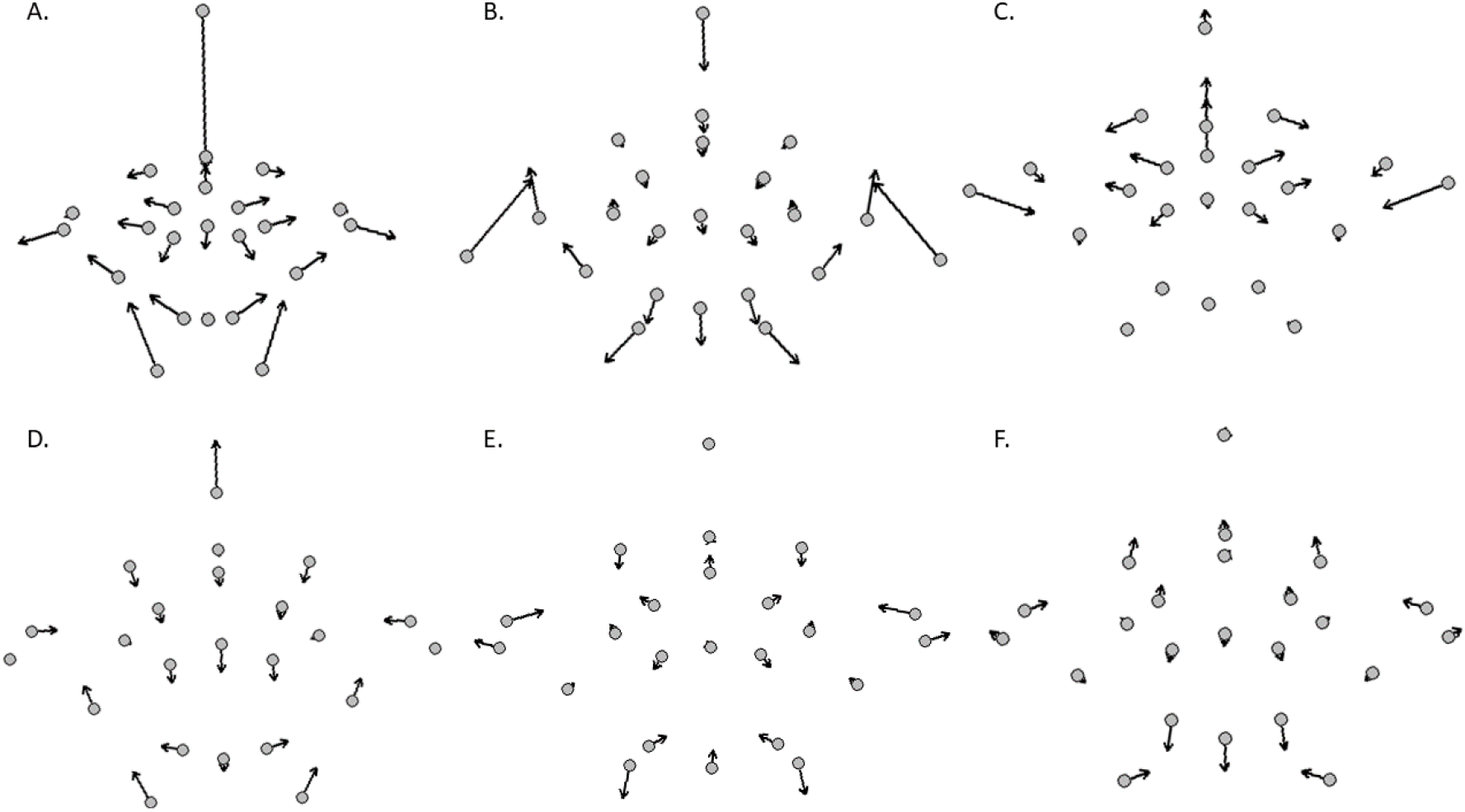
Vector plots for PCs 1-6 of the all-group analysis. The gray circles represent the negative standard deviations of the mean shape, while the arrows indicate shape change along each axis, ending at three positive standard deviations of mean shape.

PC2 represented 20.4% of explained variation, and indicated that this variation was mainly in the orientation of the laterally located structures of the vertebrae (Fig. 4B). Shapes range from more ventrally located diapophyses and more ventrally oriented prezygapophyses with shorter prezygapophyseal articular facets in the aquatic elapid sea snake, *Pelamis platura*, to more dorsally located diapophyses and dorsally oriented prezygapophyses with relatively wider prezygapophyseal articular facets in the semifossorial charinid rubber boa, *Charina bottae*. These differences were also associated with taller and shorter neural spines, respectively, and a ventral shift in the cotyl and parapophyses. Notably, PC1 and PC2 exhibited clearly visible separation of several snake clades, including the viperids, boids, charinids, and leptotyphlopids, while colubrids and elapids filled similar spaces in the two axes of variation (Fig 2A, 2B; Fig. 3).

PC3, which represented 11.6% of the overall variation, was mostly explained by variation in the dorsal region of the vertebrae, including the overall width of the vertebrae and the width of both the lateral and medial articular surfaces (Fig. 4C). Shapes for PC3 vary from relatively wider prezygapophyses and prezygapophyseal articular facets but narrower, thinner neural arches and narrower zygosphenes in the semifossorial dipsadinine colubrid, *Heterodon platirhinos,* to shorter prezygapophyses and prezygapophyseal facets but wide, convex neural arches and wide zygosphenes in the arboreal boid, *Corallus annulatus*.

PC4 represented 5.4% of the total variation and is attributable to changes in the ventral portion of the vertebrae, particularly cotylar shape and the orientation of the synapophyses (Fig. 4D). This ranges from the taller, circular cotyl and ventrally projecting synapophyses of the aquatic natricine *Liodytes pygaea*, to the wider, ellipsoidal cotyl and more laterally orientated synapophyses of the fossorial leptotyphlopid *Rena humilis*.

PC5 accounted for only 3.9% of the total variation, but exhibited important shape differences in the dorsal and ventral regions of the vertebrae (Fig. 4E). Shapes in PC5 ranged from a wider, lower, and thinner zygosphene associated with shorter parapophyses, narrower prezygapophyses, and wider prezygapophyseal articular facets in the terrestrial viperid, *Crotalus cerastes,* to a taller and thicker zygosphene associated with longer parapophyses, wider prezygapophyses, and narrower prezygapophyseal articular facets in the semiarboreal boid, *Boa constrictor*.

Finally, PC6 is mainly explained by the distance between the bottom of the cotyl and the synapophyses, as well as the relative height of the prezygapophyseal articular facets and the top of the zygosphene (Fig. 4F). This ranges from longer parapophyses projecting ventrally away from the cotyl, with a more ventrally located zygosphene and prezygapophyseal articular facets in the arboreal viperid *Bothriechis nigroviridis*, to a relatively lower cotyl and shorter, medially located parapophyses with a more dorsally located zygosphene and prezygapophyseal articular facets in the semifossorial natricine *Tropidoclonion lineatum*.

### Shape variation within Crotalinae

In the Crotalinae-only analysis, results differed somewhat overall, both in the amount of variance explained by PC and in the morphological changes themselves. The first six PCs explained 81.9% of all variation in Crotalinae (Figs. 2C, 2D; Fig. 5). A plot of all PC percentages for the Crotalinae-only analysis is available in Appendix Fig. 2.

**Figure 5.**
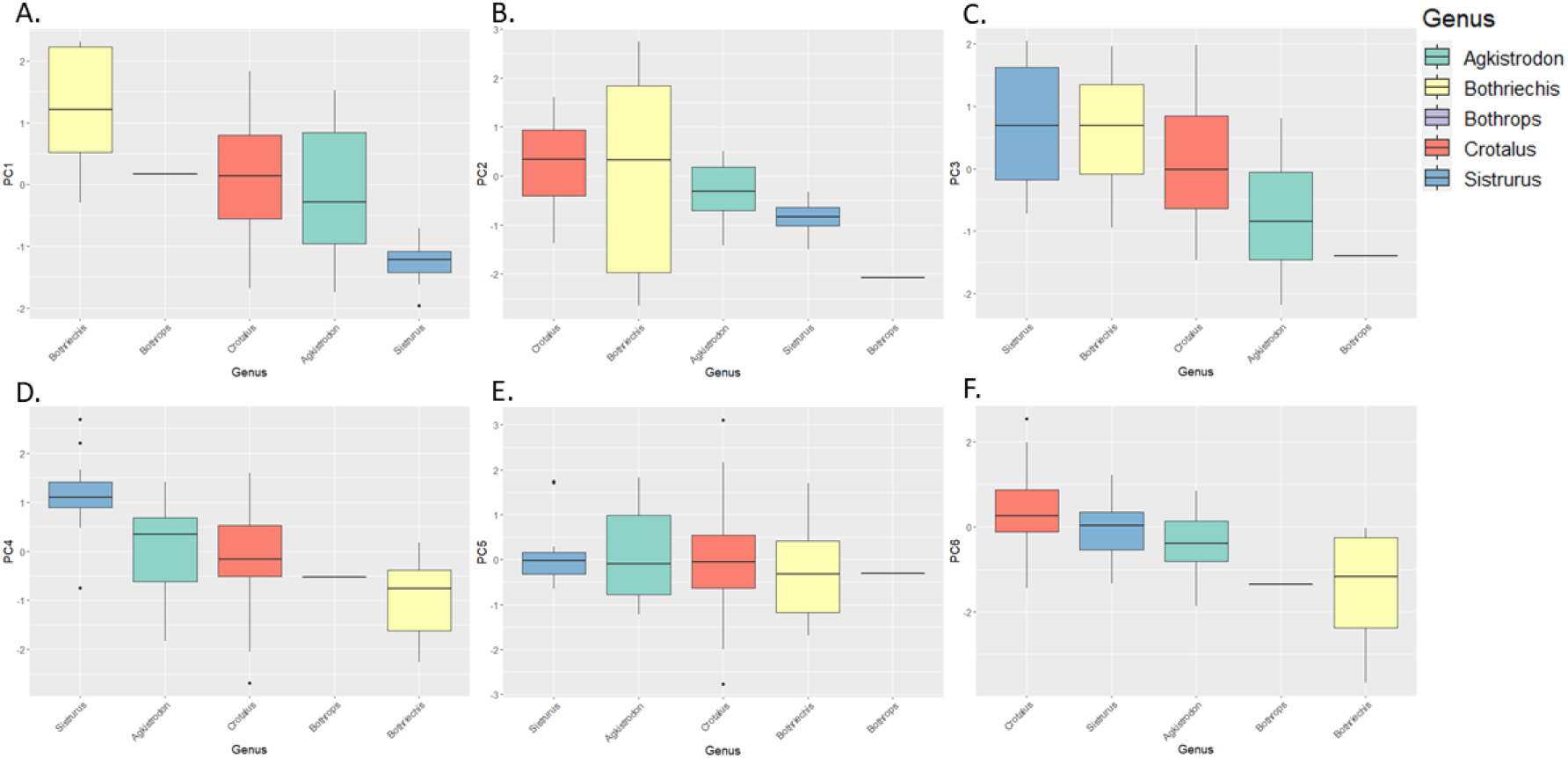
Box and whisker plots of PCs 1-6 (in numerical order) for the Crotalinae-only analysis by genus. Colors represent genus association. Lines within boxes represent median values, black circles are outliers. Single black lines without boxes represent sample sizes of n = 1.

PC1 explained 44.1 % of total variance and was similar to that of the previous analysis in that most of the variation is in the relative height of the vertebrae, composed primarily of neural spine and parapophyseal height (Fig. 6A). Unlike the previous analysis, the relative width was determined less by the prezygapophyses specifically, and was distributed more evenly throughout the vertebrae at each landmark. End members include terrestrial *Sistrurus miliarius* (negative) and arboreal *Bothriechis schlegelii* (positive). Beyond PC1, however, most of the PCs showed marked differences in the crotaline analysis compared to the all-group analysis.

**Figure 6.**
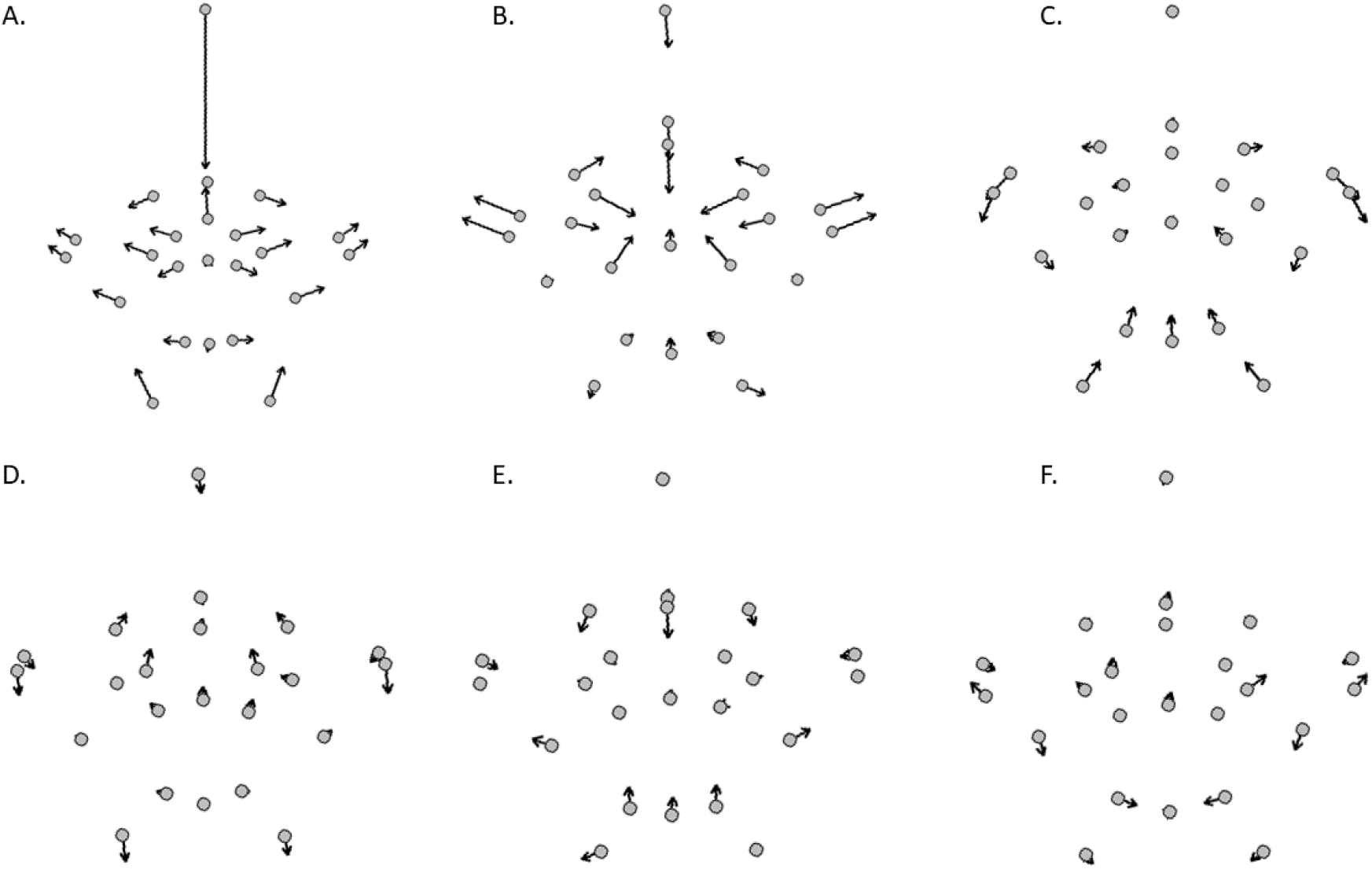
Vector plots for PCs 1-6 of the Crotalinae-only analysis (in numerical order). The gray circles represent the negative standard deviations of the mean shape, while the arrows indicate shape change along each axis, ending at three positive standard deviations of mean shape.

PC2 represented 19.5% of the total variation and was almost entirely explained by dorso-ventral compression and widening in the dorsal region of the vertebrae, where the prezygapophyses and associated articular facets would widen in parallel to the compression of the neural spine, zygosphene, and neural canal (Fig. 6B). The end members of PC2 include arboreal *Bothriechis schlegelii* (negative) and arboreal *Bothriechis nigroviridis* (positive).

PC3 explained 7.3% of the total variation, and intriguingly showed changes of the landmarks opposite to those of PC2 (Fig. 6C). Whereas PC2 explained relative width at the prezygapophyses and associated articular facets, PC3 explained the relative heights of those same structures. Similarly, while PC2 explained variation in the dorsal region of the vertebrae, PC3 explained variation in the ventral part of the vertebrae. Lower prezygapophyses and prezygapophyseal articular facets were associated with a smaller, rounder cotyl and shorter parapophyses in arboreal *Bothriechis schlegelii* (positive), and vice versa in terrestrial *Agkistrodon contortrix* (negative).

Beyond PC3, shape variation was much lower overall. PC4 represented 4.2% of variance and was attributable to the shape of the zygosphene and neural canal (Fig. 6D). Narrower, thinner zygosphenes were associated with taller neural canals in terrestrial *Sisturus miliarius* (positive), while terrestrial *Crotalus ruber* (negative) exhibited the opposing shape along this axis.

PC5 was 3.7% of total variation and showed a difference between the synapophyses compared to all other vertebral articulations (Fig. 6E). As the zygosphene, cotyl, and parapophyseal articular facets moved towards the center of the vertebrae, the parapophyses and diapophyses migrated laterally away from the center of the vertebrae. End members include terrestrial *Crotalus molossus* (negative) and terrestrial *Crotalus cerastes* (positive).

Finally, PC6 represented 3.1% of shape variation and is explained by minor differences in the orientation of the prezygapophyses and associated articular facets, as well as the orientation of the diapophyses and shape of the cotyl (Fig. 6F). Dorsally oriented prezygapophyses were associated with slightly narrower prezygapophyseal articular facets, more ventrally located diapophyses, and a narrower but slightly taller cotyl in terrestrial *Crotalus cerastes* (positive) and vice versa in arboreal *Bothriechis schlegelii* (negative).

### Multivariate analysis of snake taxonomy

The Analysis of Variance (ANOVA) was different from zero variance explained (p < 0.05) for each of the first six selected PCs at the family, subfamily, genus, and species taxonomic levels in the all-group data and for the genus and species levels in the Crotalinae-only data (Appendix Tables B2-B9). Additionally, the F-values suggest that the variation among the group means at each taxonomic level were greater than would be expected by chance. For the all-group data, Tukey’s tests at the family level suggested that much of the variation is found in the boids, charinids, leptotyphlopids, and viperids, although low sample sizes in some groups, such as in the loxocemids, may be obscuring some comparisons (Appendix Table B10). At the subfamily level, a second Tukey’s test indicated that a large number of subfamilies, including boas, sea snakes, other elapids, leptotyphlopids, vipers, and several different colubrid subfamilies, varied notably from at least one other group (Appendix Table B11). In the Crotalinae-only group, a genus-level Tukey’s test most commonly indicated differences in *Sistrurus* and *Bothriechis* from other crotalines (Appendix Table B13). *Agkistrodon* commonly differed from both those two genera and occasionally from *Crotalus*, while *Bothrops* was poorly represented in the sample, and did not allow for thorough exploration.

We applied a discriminant function analysis to all PCs to classify individuals into certain groups, and performed best at higher taxonomic levels. The overall accuracy of the discriminant function analysis was 96.0% at the family level, 85.8% at the subfamily level, 68.2% at the genus level, 58.3% at the species level, and 57.1% for primary foraging habitat in the all-group data (Appendix Tables B10-B12). In the Crotalinae-only data, percentages increased to 82.3% at the genus level, 63.3% at the species level, and 89.7% for primary foraging habitat (Appendix Tables B13, B14).

## Discussion

Geometric morphometrics have been used with increasing prevalence to examine isolated elements in both modern and fossil data (Lawing and Polly, 2010). The addition of geometric morphometrics for snake vertebral shape across a wide variety of snake species is a much-needed addition to the geometric morphometric body of literature. It opens new, replicable avenues for examining the evolution, ecology, and taxonomy of snakes in the fossil record as well as in the present. This study is the first to assess the ability of geometric morphometrics to delimit snake species using trunk vertebrae, and uses a large number of individuals and taxa from different clades. Overall, the analyses suggest that there is a strong relationship between middle trunk vertebral shape and taxonomy, with differences between groups at family to species levels. The taxonomic groups can be separated in shape space and through discriminant functions; however, groups with smaller sample sizes and at lower taxonomic levels were more difficult to delimit, as predicted.

The PCA revealed a few important patterns of vertebral morphology in snakes. Throughout all the axes of variation considered in this study, specific structures of the vertebrae were consistently notable, some of which also contributed to the relative height and width of the vertebrae: the neural spine, the prezygapophyses, and the articular surfaces – including the prezygapophyseal articular facets on the distal regions of the vertebrae, the zygosphene and cotyl on the proximal region of the vertebrae, and the rib articulations at the synapophyses and diapophyses (Figs. 4, 6). In each of these cases, the relative shape, size, and orientation of the structures produced notable variation in the total shape of the vertebra.

The plot of PC1 and PC2 for the all-group data shows distinct groupings of proportionally heavy-bodied snakes, consisting of mainly booids and viperids on the upper left (tall neural spines, more dorsally oriented prezygapophyses and diapophyses, and wider prezygapophyseal articular facets), with more slender, agile snakes, primarily consisting of colubrines towards the bottom right of the plot (Fig. 2A). Tall neural spines tend to be found on large, active snakes, while low neural spines are found on snakes that exhibit more secretive behaviors and dorso-ventrally flatten the body for tight spaces, such as in the charinids, leptotyphlopids, and some dipsadinines (Holman, 2000). The position of the diapophyses and parapophyses is indicative of the general orientation of the ribs, which is associated with the need, or lack thereof, to prevent downward displacement of muscle when performing behaviors such as cantilevering (Lillywhite, 2014). Larger prezygapophyseal articular facets with more anteriorly oriented prezygapophyses indicate different proportions and uses for axial musculature when using their primary mode of locomotion and when procuring and feeding on prey (Johnson, 1955; Lillywhite et al., 2000; Lillywhite, 2014). Finally, the anterior to posterior articulations of the cotyl to the condyle and the zygosphene to the zygantrum are indicative of the amount of torsion and flexibility allowed by the vertebrae when using muscles for movement (Johnson, 1955; Moon, 1999; Lillywhite et al., 2000).

Along each PC axis, at least one of the maximum or minimum shapes were occupied by individuals possessing a specialized behavior for procuring and feeding on prey (e.g., viperid striking) or locomotion (e.g., leptotyphlopid burrowing) in the larger dataset. Furthermore, the individuals occupying the extremes of each axis of variation belonged to different families of snakes. Within the Crotalinae-only dataset, this pattern did not remain; however, as in the larger dataset, the terrestrial taxa occupy the majority of shape space, obscuring the clearer differences in shape between ecological specialties – something clearer in the crotalines alone than when examining all snakes at once. When primary foraging habitats were examined within the dataset and along the axes of shape variation, different spaces were occupied by different foraging habitat preferences both overall and within taxonomic groups at the family level, albeit with some degree of overlap.

The plot of PC1 and PC2 shows that fossorial and semifossorial taxa occupied spaces from the highest to medial values along both axes one and two; fossorial generally occupied values higher than semifossorial taxa on axis one, but lower values than semifossorial taxa on axis two (Fig. 2). Terrestrial taxa primarily occupied medial to low value spaces along axis one, and medial areas of PC2. Arboreal and semiarboreal taxa occupied medial values to low values on axes one and two, but extended to lower values than terrestrial, fossorial, and semifossorial taxa along axis two; specifically, semiarboreal taxa did not extend as far as arboreal taxa along axis one, but reached lower values than arboreal taxa along axis two. Finally, aquatic and semiaquatic taxa filled medial to low values on axes one and two, but reached lower values than all non-terrestrial taxa on axis one and lower values than all non-arboreal or semiarboreal taxa along axis two.

Remarkably, while there was generally more overlap between primary foraging habitats than between clades, similar convergent shifts in shape occurred within clades where these habitats were concerned (e.g., the shifts between arboreal and terrestrial colubrids and boids and the differences between semiaquatic and arboreal crotalines and semiaquatic and arboreal snakes overall). These results match qualitative (Lillywhite et al., 2000), linear (Johnson, 1955), and preliminary geometric morphometric (Lawing et al., 2012) results in much smaller datasets, and suggest that vertebral shape is influenced by both phylogeny and functional demands for different ecologies. Despite what is often considered a “generalized” body form across all snakes, snake trunk vertebral morphology plays a locomotion-related functional role similar to that of limb bone proportions in mammals (Polly, 2010; Lawing et al., 2012; Short and Lawing, 2021).

Although the primary signal is taxonomic, primary foraging habitats still showed differences from the mean for all six PCs; likewise, the DFA for primary foraging habitat was similar to that of the DFA for species at 56.4% overall accuracy. Furthermore, both the PCA and the DFA showed stronger ecological differentiation when the resolution was changed to study within a specific subfamily, as in the crotaline example. This evidence suggests that snake vertebral morphology may have initially been influenced by adopting different types of predominant locomotor strategies when moving through different habitats or when using different prey capture methods. In this scenario, snakes would have then subsequently branched into other environments independently within their clades, eventually exhibiting a convergence of some shape alterations for specific primary foraging habitats. The separation of booids from most colubroids, the convergence of heavy-bodied booids and viperids in PC1 the all-group analyses, and the results of the crotaline DFA for PC1 and primary foraging habitat specifically support this interpretation.

A few considerations regarding the use of the geometric morphometric methods are necessary when interpreting results. The all-group dataset framework is best used for differentiating at higher taxonomic levels when including such a large sample and variety of groups. Similarly, the axes of shape change noted as responsible for the largest amounts of variation in the dataset are important at the scale of this study, but this may not be true for smaller analyses, such as within a single subfamily or genus. This is exemplified well in the Crotalinae-only analyses, which showed different shape variations across PC axes and better delimitation at the genus and species level as well as for primary foraging habitat in the DFA. It may therefore be necessary to run the analysis separately within smaller scales, dependent upon the goal of the study using these methods (e.g., including only the genera within a subfamily or species within a genus, as done in some other geometric morphometric papers for taxonomic delimitation).

There are several important future directions for work related to the morphometrics of snake vertebral shape. Increasing the sample size with both more taxa from around the world and with more total vertebrae per taxon would allow statistical analysis of all the groups that we were unable to include in this study. This would also allow for the identification of any changes in shape variation among different geographic regions of the world.

As one of the main purposes of this study was to create a geometric morphometric framework for use with the fossil record, implementing these analyses with the addition of fossil vertebrae is a logical step forward. Doing so will allow for objective, quantitative support for fossil identifications, will produce insights on the evolutionary relationships and ecologies of fossil snakes, and will allow for the detection of any specific changes in morphological regimes (e.g., the shift from booid- to colubroid-dominated snake assemblages in the Neogene of North America; Holman, 2000; Parmley and Hunter, 2010; Jacisin et al., 2015). Furthermore, these techniques should work when comparing fossil vertebrae whenever the homologous landmarks can be captured.

It is unlikely, based on both visual descriptions and linear measurements, that the anterior aspect of trunk vertebrae captures all diagnostic morphological variation in snakes. As such, the inclusion of other two-dimensional orientations for snake trunk vertebrae will likely produce additional information on shape variation in snakes that will improve these methods when combined with the anterior aspect. Finally, while the goal of this study was to create a simple, accessible method to improve taxonomic delimitation using snake vertebrae, we expect three-dimensional morphometrics to become more accessible and affordable in the future. As online databases of three-dimensional biological data continue to grow, techniques such as micro-computed tomography – and the programs with which to work on such data – will become more common and affordable. Three-dimensional morphometrics, then, will likely become a more commonly used, powerful tool on scales comparable to this study.

These results do not imply that geometric morphometrics should be used in lieu of gestalt and subjective visual identifications or traditional quantitative morphometrics and linear measurements, but rather as a more objective supplement to help build support for identifications in both extant and fossil skeletal material. Previous studies using geometric morphometrics for taxonomic delimitation on smaller scales have reached similar conclusions: the best results are likely to be achieved by combining morphometric shape data from homologous landmarks, traditional size morphometrics, and visual qualitative descriptions (Mutanen and Pretorius, 2007; Whitenack and Gottfried, 2010; Meik et al., 2012; Ruane, 2015). The vector plots produced practical, objective visualizations of diagnostic shape differences between taxa at specific landmarks. These methods are more effective at describing shape variation as opposed to size variation, where qualitative descriptions and numerical measurements may be more appropriate.

In conclusion, shape space of this study provides a new blueprint that can be referenced to assist in taxonomic identification and delimitation in both extant and fossil snakes using middle trunk vertebrae. Shape variation in snake trunk vertebrae show both phylogenetic and ecomorphological signals, which has implications for both the evolution of morphology between and within snake clades, and for the potential of methods such as ecometrics to be utilized in future studies. While geometric morphometrics is a powerful suite of tools with the capacity to assist in the delimitation of recent and fossil snake vertebrae, it is not yet precise enough alone with the dataset of this study to provide high confidence delimitation for some groups when analyzed for all snake groups simultaneously. Qualitative descriptions should continue to be used alongside shape analysis to identify and describe snake vertebrae in future studies.

