## Supplemental figures and tables for "Ophidian physique: the capacity of middle trunk vertebral shape for quantitative taxonomic delimitation in snakes"

**Jacisin and Lawing Supplement 1**

Contents:

Figure 1. Outlier plots for each of the 23 landmarks used in this study (in numerical order). (page 3)

Figure 2. Plot of the percent of morphological variation represented by each PC in from the whole group (left) and Crotalinae-only (right) PCA analyses. (page 4)

Table 1. List of taxa, including family, subfamily, genus, species, and primary foraging habitat. (page 5)

Table 2. ANOVA results for families within all groups. (page 19)

Table 3. ANOVA results for subfamilies within all groups. (page 20)

Table 4. ANOVA results for genera within all groups. (page 21)

Table 5. ANOVA results for species within all groups. (page 22)

Table 6. ANOVA results for primary foraging habitats within all groups. (page 23)

Table 7. ANOVA results for genera within Crotalinae. (page 24)

Table 8. ANOVA results for species within Crotalinae. (page 25)

Table 9. ANOVA results for primary foraging ecology within Crotalinae. (page 26)

Table 10. Tukey's test results of family-level taxonomy for PCs 1-6 of the all-groups data. (page 27)

Table 11. Tukey's test results of subfamily taxonomy for PCs 1-6 for all groups. (page 36)

Table 12. Tukey's test results of primary foraging habitat for PCs 1-6 of the all-groups data. (page 57)

Table 13. Tukey's test results of genus-level taxonomy for PCs 1-6 of the Crotalinae-only data. (page 61)

Table 14. Tukey's test results of primary foraging habitat for PCs 1-6 of the Crotalinae-only data. (page 63)

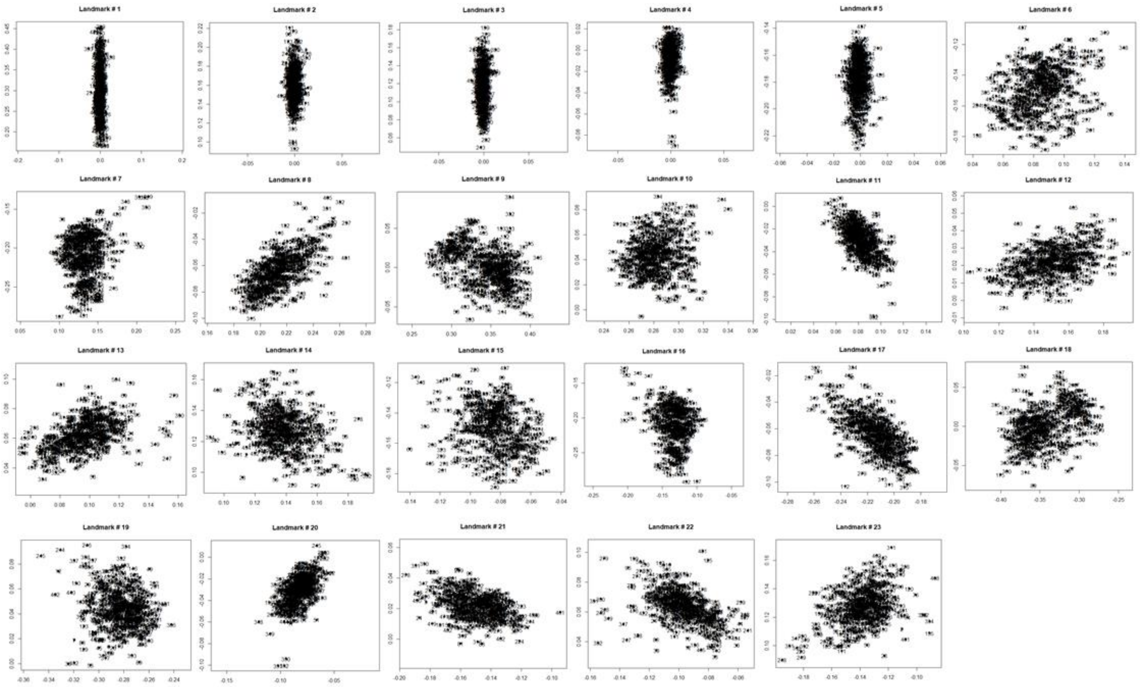


**Figure 1. Outlier plots for each of the 23 landmarks used in this study (in numerical order).**

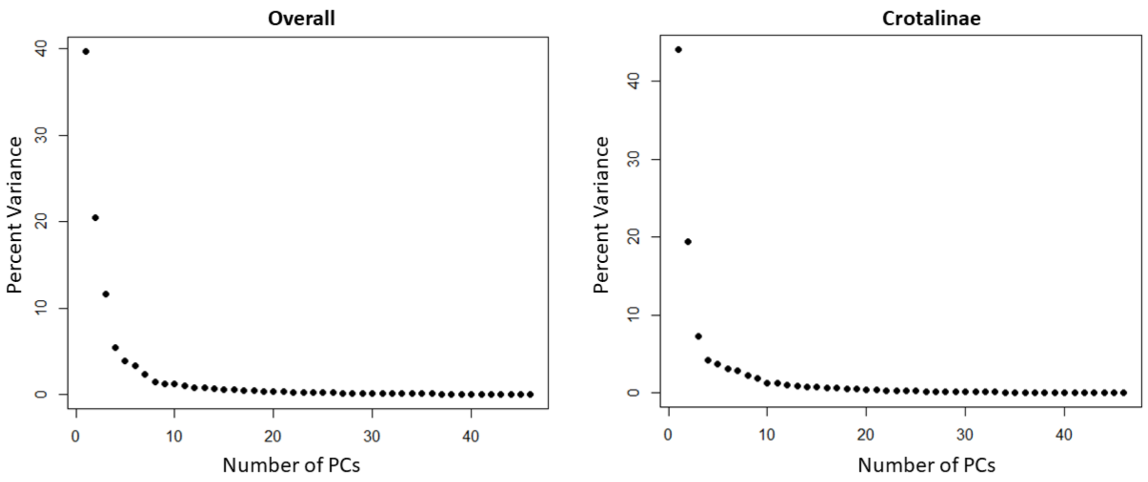


**Figure 2. Plot of the percent of morphological variation represented by each PC in from the whole group (left) and Crotalinae-only (right) PCA analyses.**

**Table 1. List of taxa, including family, subfamily, genus, species, and primary foraging habitat.**

| ID | Genus | Species | Family | Subfamily | Substrate |
|---|---|---|---|---|---|
| 1 | *Agkistrodon* | *contortrix* | Viperidae | Crotalinae | Terrestrial |
| 2 | *Agkistrodon* | *piscivorus* | Viperidae | Crotalinae | Semiaquatic |
| 3 | *Alsophis* | *antillensis* | Colubridae | Dipsadinae | Terrestrial |
| 4 | *Alsophis* | *cantherigerus* | Colubridae | Dipsadinae | Terrestrial |
| 5 | *Alsophis* | *portoricensis* | Colubridae | Dipsadinae | Terrestrial |
| 6 | *Alsophis* | *vudii* | Colubridae | Dipsadinae | Terrestrial |
| 7 | *Antillophis* | *parvifrons* | Colubridae | Dipsadinae | Terrestrial |
| 8 | *Atractus* | *trilineatus* | Colubridae | Dipsadinae | Semifossorial |
| 9 | *Boa* | *constrictor* | Boidae | Boinae | Semiarboreal |
| 10 | *Bogertophis* | *subocularis* | Colubridae | Colubrinae | Terrestrial |
| 11 | *Bothriechis* | *nigroviridis* | Viperidae | Crotalinae | Arboreal |
| 12 | *Bothriechis* | *schlegelii* | Viperidae | Crotalinae | Arboreal |
| 13 | *Bothrops* | *asper* | Viperidae | Crotalinae | Terrestrial |
| 14 | *Candoia* | *carinata* | Candoiidae | Candoiinae | Terrestrial |
| 15 | *Carphophis* | *amoenus* | Colubridae | Dipsadinae | Fossorial |
| 16 | *Cemophora* | *coccinea* | Colubridae | Colubrinae | Fossorial |
| 17 | *Charina* | *bottae* | Charinidae | Charininae | Semifossorial |
| 18 | *Chilomeniscus* | *stramineus* | Colubridae | Colubrinae | Fossorial |
| 19 | *Chironius* | *carinatus* | Colubridae | Colubrinae | Arboreal |
| 20 | *Chironius* | *scurrulus* | Colubridae | Colubrinae | Arboreal |
| 21 | *Clelia* | *clelia* | Colubridae | Dipsadinae | Terrestrial |
| 22 | *Coluber* | *constrictor* | Colubridae | Colubrinae | Semiarboreal |
| 23 | *Conophis* | *pulcher* | Colubridae | Dipsadinae | Terrestrial |
| 24 | *Contia* | *tenuis* | Colubridae | Dipsadinae | Semifossorial |
| 25 | *Crotalus* | *adamanteus* | Viperidae | Crotalinae | Terrestrial |
| 26 | *Crotalus* | *atrox* | Viperidae | Crotalinae | Terrestrial |
| 27 | *Crotalus* | *cerastes* | Viperidae | Crotalinae | Terrestrial |
| 28 | *Crotalus* | *enyo* | Viperidae | Crotalinae | Terrestrial |
| 29 | *Crotalus* | *horridus* | Viperidae | Crotalinae | Terrestrial |
| 30 | *Crotalus* | *mitchellii* | Viperidae | Crotalinae | Terrestrial |
| 31 | *Crotalus* | *molossus* | Viperidae | Crotalinae | Terrestrial |
| 32 | *Crotalus* | *ravus* | Viperidae | Crotalinae | Terrestrial |
| 33 | *Crotalus* | *ruber* | Viperidae | Crotalinae | Terrestrial |
| 34 | *Crotalus* | *scutulatus* | Viperidae | Crotalinae | Terrestrial |

**Table 1. (continued)**

| | | | | | |
|---|---|---|---|---|---|
| 35 | *Crotalus* | *viridis* | Viperidae | Crotalinae | Terrestrial |
| 36 | *Dendrophidion* | *vinitor* | Colubridae | Colubrinae | Semiarboreal |
| 37 | *Diadophis* | *punctatus* | Colubridae | Dipsadinae | Semifossorial |
| 38 | *Dipsas* | *articulata* | Colubridae | Dipsadinae | Arboreal |
| 39 | *Dipsas* | *variegata* | Colubridae | Dipsadinae | Arboreal |
| 40 | *Drymarchon* | *corais* | Colubridae | Colubrinae | Terrestrial |
| 41 | *Chilabothrus* | *angulifer* | Boidae | Boinae | Arboreal |
| 42 | *Epicrates* | *cenchria* | Boidae | Boinae | Terrestrial |
| 43 | *Chilabothrus* | *striatus* | Boidae | Boinae | Terrestrial |
| 44 | *Chilabothrus* | *subflavus* | Boidae | Boinae | Arboreal |
| 45 | *Erythrolamprus* | *aesculapii* | Colubridae | Dipsadinae | Terrestrial |
| 46 | *Erythrolamprus* | *ocellatus* | Colubridae | Dipsadinae | Terrestrial |
| 47 | *Farancia* | *abacura* | Colubridae | Dipsadinae | Semiaquatic |
| 48 | *Farancia* | *erytrogramma* | Colubridae | Dipsadinae | Aquatic |
| 49 | *Heterodon* | *nasicus* | Colubridae | Dipsadinae | Semifossorial |
| 50 | *Heterodon* | *platirhinos* | Colubridae | Dipsadinae | Semifossorial |
| 51 | *Heterodon* | *simus* | Colubridae | Dipsadinae | Semifossorial |
| 52 | *Hypsirhynchus* | *ferox* | Colubridae | Dipsadinae | Terrestrial |
| 53 | *Imantodes* | *cenchoa* | Colubridae | Dipsadinae | Arboreal |
| 54 | *Lampropeltis* | *calligaster* | Colubridae | Colubrinae | Semifossorial |
| 55 | *Lampropeltis* | *extenuata* | Colubridae | Colubrinae | Fossorial |
| 56 | *Lampropeltis* | *getula* | Colubridae | Colubrinae | Terrestrial |
| 57 | *Lampropeltis* | *triangulum* | Colubridae | Colubrinae | Semifossorial |
| 58 | *Leptodeira* | *annulata* | Colubridae | Dipsadinae | Semiaquatic |
| 59 | *Leptophis* | *ahaetulla* | Colubridae | Colubrinae | Arboreal |
| 60 | *Lichanura* | *trivirgata* | Charinidae | Charininae | Semifossorial |
| 61 | *Erythrolamprus* | *cobella* | Colubridae | Dipsadinae | Semiaquatic |
| 62 | *Erythrolamprus* | *melanotus* | Colubridae | Dipsadinae | Semiaquatic |
| 63 | *Erythrolamprus* | *reginae* | Colubridae | Dipsadinae | Aquatic |
| 64 | *Loxocemus* | *bicolor* | Loxocemidae | Loxoceminae | Semifossorial |
| 65 | *Masticophis* | *flagellum* | Colubridae | Colubrinae | Semiarboreal |
| 66 | *Masticophis* | *taeniatus* | Colubridae | Colubrinae | Arboreal |
| 67 | *Mastigodryas* | *pleei* | Colubridae | Colubrinae | Arboreal |
| 68 | *Micruroides* | *euryxanthus* | Elapidae | Elapinae | Fossorial |
| 69 | *Micrurus* | *diastema* | Elapidae | Elapinae | Terrestrial |
| 70 | *Micrurus* | *fulvius* | Elapidae | Elapinae | Fossorial |
| 71 | *Micrurus* | *psyches* | Elapidae | Elapinae | Fossorial |

**Table 1. (continued)**

| | | | | | |
|---|---|---|---|---|---|
| 72 | *Nerodia* | *cyclopion* | Colubridae | Natricinae | Aquatic |
| 73 | *Nerodia* | *erythrogaster* | Colubridae | Natricinae | Semiaquatic |
| 74 | *Nerodia* | *fasciata* | Colubridae | Natricinae | Aquatic |
| 75 | *Nerodia* | *floridana* | Colubridae | Natricinae | Aquatic |
| 76 | *Nerodia* | *rhombifer* | Colubridae | Natricinae | Semiaquatic |
| 77 | *Nerodia* | *sipedon* | Colubridae | Natricinae | Aquatic |
| 78 | *Nerodia* | *taxispilota* | Colubridae | Natricinae | Aquatic |
| 79 | *Ninia* | *atrata* | Colubridae | Dipsadinae | Semifossorial |
| 80 | *Opheodrys* | *aestivus* | Colubridae | Colubrinae | Arboreal |
| 81 | *Opheodrys* | *vernalis* | Colubridae | Colubrinae | Semiarboreal |
| 82 | *Oxybelis* | *aeneus* | Colubridae | Colubrinae | Arboreal |
| 83 | *Oxybelis* | *brevirostris* | Colubridae | Colubrinae | Arboreal |
| 84 | *Oxybelis* | *fulgidus* | Colubridae | Colubrinae | Arboreal |
| 85 | *Oxyrhopus* | *petolarius* | Colubridae | Dipsadinae | Terrestrial |
| 86 | *Pantherophis* | *guttatus* | Colubridae | Colubrinae | Terrestrial |
| 87 | *Pantherophis* | *obsoletus* | Colubridae | Colubrinae | Arboreal |
| 88 | *Pantherophis* | *vulpinus* | Colubridae | Colubrinae | Terrestrial |
| 89 | *Pelamis* | *platura* | Elapidae | Hydrophiinae | Aquatic |
| 90 | *Phyllorhynchus* | *decurtatus* | Colubridae | Colubrinae | Fossorial |
| 91 | *Pituophis* | *catenifer* | Colubridae | Colubrinae | Semifossorial |
| 92 | *Pituophis* | *deppei* | Colubridae | Colubrinae | Semifossorial |
| 93 | *Pituophis* | *melanoleucus* | Colubridae | Colubrinae | Semifossorial |
| 94 | *Phrynonax* | *poecilonotus* | Colubridae | Colubrinae | Semiarboreal |
| 95 | *Spilotes* | *sulphureus* | Colubridae | Colubrinae | Semiarboreal |
| 96 | *Regina* | *grahami* | Colubridae | Natricinae | Aquatic |
| 97 | *Regina* | *septemvittata* | Colubridae | Natricinae | Semiaquatic |
| 98 | *Rhadinaea* | *flavilata* | Colubridae | Colubrinae | Terrestrial |
| 99 | *Rhinocheilus* | *lecontei* | Colubridae | Colubrinae | Semifossorial |
| 100 | *Salvadora* | *grahamiae* | Colubridae | Colubrinae | Semiarboreal |
| 101 | *Salvadora* | *hexalepis* | Colubridae | Colubrinae | Semiarboreal |
| 102 | *Salvadora* | *intermedia* | Colubridae | Colubrinae | Semiarboreal |
| 103 | *Seminatrix* | *pygaea* | Colubridae | Natricinae | Aquatic |
| 104 | *Sibon* | *anthracops* | Colubridae | Dipsadinae | Arboreal |
| 105 | *Sibon* | *nebulatus* | Colubridae | Dipsadinae | Arboreal |
| 106 | *Sistrurus* | *catenatus* | Viperidae | Crotalinae | Terrestrial |
| 107 | *Sistrurus* | *miliarius* | Viperidae | Crotalinae | Terrestrial |
| 108 | *Spilotes* | *pullatus* | Colubridae | Colubrinae | Semiarboreal |

**Table 1. (continued)**

| | | | | | |
|---|---|---|---|---|---|
| 109 | *Stenorrhina* | *degenhardtii* | Colubridae | Colubrinae | Terrestrial |
| 110 | *Storeria* | *dekayi* | Colubridae | Natricinae | Terrestrial |
| 111 | *Storeria* | *occipitomaculata* | Colubridae | Natricinae | Terrestrial |
| 112 | *Tantilla* | *relicta* | Colubridae | Colubrinae | Fossorial |
| 113 | *Thamnophis* | *brachystoma* | Colubridae | Natricinae | Terrestrial |
| 114 | *Thamnophis* | *butleri* | Colubridae | Natricinae | Terrestrial |
| 115 | *Thamnophis* | *cyrtopsis* | Colubridae | Natricinae | Terrestrial |
| 116 | *Thamnophis* | *elegans* | Colubridae | Natricinae | Terrestrial |
| 117 | *Thamnophis* | *eques* | Colubridae | Natricinae | Terrestrial |
| 118 | *Thamnophis* | *marcianus* | Colubridae | Natricinae | Semiaquatic |
| 119 | *Thamnophis* | *proximus* | Colubridae | Natricinae | Semiaquatic |
| 120 | *Thamnophis* | *radix* | Colubridae | Natricinae | Terrestrial |
| 121 | *Thamnophis* | *sauritus* | Colubridae | Natricinae | Semiaquatic |
| 122 | *Thamnophis* | *sirtalis* | Colubridae | Natricinae | Semiaquatic |
| 123 | *Trimorphodon* | *biscutatus* | Colubridae | Colubrinae | Terrestrial |
| 124 | *Siphlophis* | *compressus* | Colubridae | Dipsadinae | Terrestrial |
| 125 | *Tropidoclonion* | *lineatum* | Colubridae | Natricinae | Semifossorial |
| 126 | *Tropidophis* | *canus* | Tropidophiidae | Tropidophiinae | Fossorial |
| 127 | *Tropidophis* | *haetianus* | Tropidophiidae | Tropidophiinae | Fossorial |
| 128 | *Uromacer* | *catesbyi* | Colubridae | Dipsadinae | Arboreal |
| 129 | *Uromacer* | *oxyrhynchus* | Colubridae | Dipsadinae | Arboreal |
| 130 | *Virginia* | *valeriae* | Colubridae | Natricinae | Semifossorial |
| 131 | *Nerodia* | *harteri* | Colubridae | Natricinae | Aquatic |
| 132 | *Heterodon* | *nasicus* | Colubridae | Dipsadinae | Semifossorial |
| 133 | *Heterodon* | *nasicus* | Colubridae | Dipsadinae | Semifossorial |
| 134 | *Agkistrodon* | *piscivorus* | Viperidae | Crotalinae | Semiaquatic |
| 135 | *Crotalus* | *atrox* | Viperidae | Crotalinae | Terrestrial |
| 136 | *Crotalus* | *atrox* | Viperidae | Crotalinae | Terrestrial |
| 137 | *Crotalus* | *atrox* | Viperidae | Crotalinae | Terrestrial |
| 138 | *Agkistrodon* | *piscivorus* | Viperidae | Crotalinae | Semiaquatic |
| 139 | *Agkistrodon* | *piscivorus* | Viperidae | Crotalinae | Semiaquatic |
| 140 | *Agkistrodon* | *piscivorus* | Viperidae | Crotalinae | Semiaquatic |
| 141 | *Nerodia* | *rhombifer* | Colubridae | Natricinae | Semiaquatic |
| 142 | *Nerodia* | *rhombifer* | Colubridae | Natricinae | Semiaquatic |
| 143 | *Nerodia* | *rhombifer* | Colubridae | Natricinae | Semiaquatic |
| 144 | *Heterodon* | *nasicus* | Colubridae | Dipsadinae | Semifossorial |
| 145 | *Heterodon* | *nasicus* | Colubridae | Dipsadinae | Semifossorial |

**Table1. (continued)**

| | | | | | |
|---|---|---|---|---|---|
| 146 | *Heterodon* | *nasicus* | Colubridae | Dipsadinae | Semifossorial |
| 147 | *Crotalus* | *atrox* | Viperidae | Crotalinae | Terrestrial |
| 148 | *Crotalus* | *atrox* | Viperidae | Crotalinae | Terrestrial |
| 149 | *Crotalus* | *atrox* | Viperidae | Crotalinae | Terrestrial |
| 150 | *Crotalus* | *molossus* | Viperidae | Crotalinae | Terrestrial |
| 151 | *Pantherophis* | *obsoletus* | Colubridae | Colubrinae | Arboreal |
| 152 | *Pantherophis* | *obsoletus* | Colubridae | Colubrinae | Arboreal |
| 153 | *Pantherophis* | *obsoletus* | Colubridae | Colubrinae | Arboreal |
| 154 | *Pantherophis* | *obsoletus* | Colubridae | Colubrinae | Arboreal |
| 155 | *Nerodia* | *rhombifer* | Colubridae | Natricinae | Semiaquatic |
| 156 | *Crotalus* | *lepidus* | Viperidae | Crotalinae | Terrestrial |
| 157 | *Crotalus* | *scutulatus* | Viperidae | Crotalinae | Terrestrial |
| 158 | *Crotalus* | *tigris* | Viperidae | Crotalinae | Terrestrial |
| 159 | *Micruroides* | *euryxanthus* | Elapidae | Elapinae | Fossorial |
| 160 | *Arizona* | *elegans* | Colubridae | Colubrinae | Semifossorial |
| 161 | *Chionactis* | *occipitalis* | Colubridae | Colubrinae | Fossorial |
| 162 | *Chilomeniscus* | *stramineus* | Colubridae | Colubrinae | Fossorial |
| 163 | *Diadophis* | *punctatus* | Colubridae | Dipsadinae | Semifossorial |
| 164 | *Senticolis* | *triaspis* | Colubridae | Colubrinae | Semiarboreal |
| 165 | *Senticolis* | *triaspis* | Colubridae | Colubrinae | Semiarboreal |
| 166 | *Bogertophis* | *subocularis* | Colubridae | Colubrinae | Terrestrial |
| 167 | *Hypsiglena* | *torquata* | Colubridae | Dipsadinae | Terrestrial |
| 168 | *Lampropeltis* | *pyromelana* | Colubridae | Colubrinae | Terrestrial |
| 169 | *Lampropeltis* | *pyromelana* | Colubridae | Colubrinae | Terrestrial |
| 170 | *Lampropeltis* | *pyromelana* | Colubridae | Colubrinae | Terrestrial |
| 171 | *Lampropeltis* | *triangulum* | Colubridae | Colubrinae | Semifossorial |
| 172 | *Lampropeltis* | *triangulum* | Colubridae | Colubrinae | Semifossorial |
| 173 | *Lampropeltis* | *triangulum* | Colubridae | Colubrinae | Semifossorial |
| 174 | *Lampropeltis* | *triangulum* | Colubridae | Colubrinae | Semifossorial |
| 175 | *Masticophis* | *bilineatus* | Colubridae | Colubrinae | Semiarboreal |
| 176 | *Masticophis* | *flagellum* | Colubridae | Colubrinae | Semiarboreal |
| 177 | *Nerodia* | *erythrogaster* | Colubridae | Natricinae | Semiaquatic |
| 178 | *Nerodia* | *fasciata* | Colubridae | Natricinae | Aquatic |
| 179 | *Nerodia* | *rhombifer* | Colubridae | Natricinae | Semiaquatic |
| 180 | *Nerodia* | *sipedon* | Colubridae | Natricinae | Aquatic |
| 181 | *Opheodrys* | *aestivus* | Colubridae | Colubrinae | Arboreal |
| 182 | *Phyllorhynchus* | *decurtatus* | Colubridae | Colubrinae | Fossorial |

**Table 1. (continued)**

| | | | | | |
|---|---|---|---|---|---|
| 183 | *Rhinocheilus* | *lecontei* | Colubridae | Colubrinae | Semifossorial |
| 184 | *Salvadora* | *hexalepis* | Colubridae | Colubrinae | Semiarboreal |
| 185 | *Trimorphodon* | *lambda* | Colubridae | Colubrinae | Terrestrial |
| 186 | *Trimorphodon* | *biscutatus* | Colubridae | Colubrinae | Terrestrial |
| 187 | *Thamnophis* | *cyrtopsis* | Colubridae | Natricinae | Terrestrial |
| 188 | *Thamnophis* | *elegans* | Colubridae | Natricinae | Terrestrial |
| 189 | *Thamnophis* | *eques* | Colubridae | Natricinae | Terrestrial |
| 190 | *Thamnophis* | *marcianus* | Colubridae | Natricinae | Semiaquatic |
| 191 | *Thamnophis* | *marcianus* | Colubridae | Natricinae | Semiaquatic |
| 192 | *Thamnophis* | *proximus* | Colubridae | Natricinae | Semiaquatic |
| 193 | *Thamnophis* | *rufipunctatus* | Colubridae | Natricinae | Aquatic |
| 194 | *Thamnophis* | *sirtalis* | Colubridae | Natricinae | Semiaquatic |
| 195 | *Agkistrodon* | *contortrix* | Viperidae | Crotalinae | Terrestrial |
| 196 | *Agkistrodon* | *contortrix* | Viperidae | Crotalinae | Terrestrial |
| 197 | *Agkistrodon* | *contortrix* | Viperidae | Crotalinae | Terrestrial |
| 198 | *Agkistrodon* | *contortrix* | Viperidae | Crotalinae | Terrestrial |
| 199 | *Agkistrodon* | *contortrix* | Viperidae | Crotalinae | Terrestrial |
| 200 | *Agkistrodon* | *contortrix* | Viperidae | Crotalinae | Terrestrial |
| 201 | *Agkistrodon* | *contortrix* | Viperidae | Crotalinae | Terrestrial |
| 202 | *Agkistrodon* | *contortrix* | Viperidae | Crotalinae | Terrestrial |
| 203 | *Pantherophis* | *guttatus* | Colubridae | Colubrinae | Terrestrial |
| 204 | *Pantherophis* | *emoryi* | Colubridae | Colubrinae | Terrestrial |
| 205 | *Pantherophis* | *emoryi* | Colubridae | Colubrinae | Terrestrial |
| 206 | *Pantherophis* | *emoryi* | Colubridae | Colubrinae | Terrestrial |
| 207 | *Pantherophis* | *emoryi* | Colubridae | Colubrinae | Terrestrial |
| 208 | *Pantherophis* | *emoryi* | Colubridae | Colubrinae | Terrestrial |
| 209 | *Pantherophis* | *emoryi* | Colubridae | Colubrinae | Terrestrial |
| 210 | *Pantherophis* | *emoryi* | Colubridae | Colubrinae | Terrestrial |
| 211 | *Pantherophis* | *guttatus* | Colubridae | Colubrinae | Terrestrial |
| 212 | *Pantherophis* | *guttatus* | Colubridae | Colubrinae | Terrestrial |
| 213 | *Pantherophis* | *guttatus* | Colubridae | Colubrinae | Terrestrial |
| 214 | *Pantherophis* | *guttatus* | Colubridae | Colubrinae | Terrestrial |
| 215 | *Micrurus* | *tener* | Elapidae | Elapinae | Fossorial |
| 216 | *Micrurus* | *tener* | Elapidae | Elapinae | Fossorial |
| 217 | *Micrurus* | *tener* | Elapidae | Elapinae | Fossorial |
| 218 | *Micrurus* | *tener* | Elapidae | Elapinae | Fossorial |
| 219 | *Phyllorhynchus* | *sp* | Colubridae | Colubrinae | Fossorial |

**Table 1. (continued)**

| | | | | | |
|---|---|---|---|---|---|
| 220 | *Phyllorhynchus* | *sp* | Colubridae | Colubrinae | Fossorial |
| 221 | *Lampropeltis* | *calligaster* | Colubridae | Colubrinae | Semifossorial |
| 222 | *Lampropeltis* | *calligaster* | Colubridae | Colubrinae | Semifossorial |
| 223 | *Lampropeltis* | *calligaster* | Colubridae | Colubrinae | Semifossorial |
| 224 | *Drymarchon* | *couperi* | Colubridae | Colubrinae | Terrestrial |
| 225 | *Sonora* | *semiannulata* | Colubridae | Colubrinae | Semifossorial |
| 226 | *Sonora* | *semiannulata* | Colubridae | Colubrinae | Semifossorial |
| 227 | *Crotalus* | *molossus* | Viperidae | Crotalinae | Terrestrial |
| 228 | *Crotalus* | *molossus* | Viperidae | Crotalinae | Terrestrial |
| 229 | *Crotalus* | *molossus* | Viperidae | Crotalinae | Terrestrial |
| 230 | *Crotalus* | *molossus* | Viperidae | Crotalinae | Terrestrial |
| 231 | *Crotalus* | *molossus* | Viperidae | Crotalinae | Terrestrial |
| 232 | *Arizona* | *elegans* | Colubridae | Colubrinae | Semifossorial |
| 233 | *Arizona* | *elegans* | Colubridae | Colubrinae | Semifossorial |
| 234 | *Arizona* | *elegans* | Colubridae | Colubrinae | Semifossorial |
| 235 | *Arizona* | *elegans* | Colubridae | Colubrinae | Semifossorial |
| 236 | *Arizona* | *elegans* | Colubridae | Colubrinae | Semifossorial |
| 237 | *Arizona* | *elegans* | Colubridae | Colubrinae | Semifossorial |
| 238 | *Boa* | *constrictor* | Boidae | Boinae | Semiarboreal |
| 239 | *Boa* | *constrictor* | Boidae | Boinae | Semiarboreal |
| 240 | *Boa* | *constrictor* | Boidae | Boinae | Semiarboreal |
| 241 | *Boa* | *constrictor* | Boidae | Boinae | Semiarboreal |
| 242 | *Boa* | *constrictor* | Boidae | Boinae | Semiarboreal |
| 243 | *Boa* | *constrictor* | Boidae | Boinae | Semiarboreal |
| 244 | *Bothriechis* | *nigroviridis* | Viperidae | Crotalinae | Arboreal |
| 245 | *Bothriechis* | *nigroviridis* | Viperidae | Crotalinae | Arboreal |
| 246 | *Bothriechis* | *nigroviridis* | Viperidae | Crotalinae | Arboreal |
| 247 | *Bothriechis* | *schlegelii* | Viperidae | Crotalinae | Arboreal |
| 248 | *Bothriechis* | *schlegelii* | Viperidae | Crotalinae | Arboreal |
| 249 | *Bothriechis* | *schlegelii* | Viperidae | Crotalinae | Arboreal |
| 250 | *Carphophis* | *amoenus* | Colubridae | Dipsadinae | Fossorial |
| 251 | *Carphophis* | *amoenus* | Colubridae | Dipsadinae | Fossorial |
| 252 | *Carphophis* | *amoenus* | Colubridae | Dipsadinae | Fossorial |
| 253 | *Cemophora* | *coccinea* | Colubridae | Colubrinae | Fossorial |
| 254 | *Cemophora* | *coccinea* | Colubridae | Colubrinae | Fossorial |
| 255 | *Cemophora* | *coccinea* | Colubridae | Colubrinae | Fossorial |
| 256 | *Charina* | *bottae* | Charinidae | Charininae | Semifossorial |

**Table 1. (continued)**

| | | | | | |
|---|---|---|---|---|---|
| 257 | *Charina* | *bottae* | Charinidae | Charininae | Semifossorial |
| 258 | *Charina* | *bottae* | Charinidae | Charininae | Semifossorial |
| 259 | *Lichanura* | *trivirgata* | Charinidae | Charininae | Semifossorial |
| 260 | *Lichanura* | *trivirgata* | Charinidae | Charininae | Semifossorial |
| 261 | *Lichanura* | *trivirgata* | Charinidae | Charininae | Semifossorial |
| 262 | *Chilomeniscus* | *stramineus* | Colubridae | Colubrinae | Fossorial |
| 263 | *Chilomeniscus* | *stramineus* | Colubridae | Colubrinae | Fossorial |
| 264 | *Chilomeniscus* | *stramineus* | Colubridae | Colubrinae | Fossorial |
| 265 | *Chilomeniscus* | *stramineus* | Colubridae | Colubrinae | Fossorial |
| 266 | *Chilomeniscus* | *stramineus* | Colubridae | Colubrinae | Fossorial |
| 267 | *Chilomeniscus* | *stramineus* | Colubridae | Colubrinae | Fossorial |
| 268 | *Chilomeniscus* | *stramineus* | Colubridae | Colubrinae | Fossorial |
| 269 | *Chilomeniscus* | *stramineus* | Colubridae | Colubrinae | Fossorial |
| 270 | *Chrysopelea* | *ornata* | Colubridae | Ahaetuliinae | Arboreal |
| 271 | *Coluber* | *constrictor* | Colubridae | Colubrinae | Semiarboreal |
| 272 | *Coluber* | *constrictor* | Colubridae | Colubrinae | Semiarboreal |
| 273 | *Coluber* | *constrictor* | Colubridae | Colubrinae | Semiarboreal |
| 274 | *Coluber* | *constrictor* | Colubridae | Colubrinae | Semiarboreal |
| 275 | *Coluber* | *constrictor* | Colubridae | Colubrinae | Semiarboreal |
| 276 | *Coluber* | *constrictor* | Colubridae | Colubrinae | Semiarboreal |
| 277 | *Corallus* | *annulatus* | Boidae | Boinae | Arboreal |
| 278 | *Corallus* | *annulatus* | Boidae | Boinae | Arboreal |
| 279 | *Corallus* | *annulatus* | Boidae | Boinae | Arboreal |
| 280 | *Crotalus* | *adamanteus* | Viperidae | Crotalinae | Terrestrial |
| 281 | *Crotalus* | *adamanteus* | Viperidae | Crotalinae | Terrestrial |
| 282 | *Crotalus* | *adamanteus* | Viperidae | Crotalinae | Terrestrial |
| 283 | *Crotalus* | *cerastes* | Viperidae | Crotalinae | Terrestrial |
| 284 | *Crotalus* | *cerastes* | Viperidae | Crotalinae | Terrestrial |
| 285 | *Crotalus* | *cerastes* | Viperidae | Crotalinae | Terrestrial |
| 286 | *Crotalus* | *cerastes* | Viperidae | Crotalinae | Terrestrial |
| 287 | *Crotalus* | *cerastes* | Viperidae | Crotalinae | Terrestrial |
| 288 | *Crotalus* | *cerastes* | Viperidae | Crotalinae | Terrestrial |
| 289 | *Crotalus* | *enyo* | Viperidae | Crotalinae | Terrestrial |
| 290 | *Crotalus* | *enyo* | Viperidae | Crotalinae | Terrestrial |
| 291 | *Crotalus* | *enyo* | Viperidae | Crotalinae | Terrestrial |
| 292 | *Crotalus* | *horridus* | Viperidae | Crotalinae | Terrestrial |
| 293 | *Crotalus* | *horridus* | Viperidae | Crotalinae | Terrestrial |

**Table 1. (continued)**

| | | | | | |
|---|---|---|---|---|---|
| 294 | *Crotalus* | *horridus* | Viperidae | Crotalinae | Terrestrial |
| 295 | *Crotalus* | *horridus* | Viperidae | Crotalinae | Terrestrial |
| 296 | *Crotalus* | *mitchellii* | Viperidae | Crotalinae | Terrestrial |
| 297 | *Crotalus* | *mitchellii* | Viperidae | Crotalinae | Terrestrial |
| 298 | *Crotalus* | *mitchellii* | Viperidae | Crotalinae | Terrestrial |
| 299 | *Crotalus* | *mitchellii* | Viperidae | Crotalinae | Terrestrial |
| 300 | *Crotalus* | *mitchellii* | Viperidae | Crotalinae | Terrestrial |
| 301 | *Crotalus* | *mitchellii* | Viperidae | Crotalinae | Terrestrial |
| 302 | *Crotalus* | *oreganus* | Viperidae | Crotalinae | Terrestrial |
| 303 | *Crotalus* | *oreganus* | Viperidae | Crotalinae | Terrestrial |
| 304 | *Crotalus* | *oreganus* | Viperidae | Crotalinae | Terrestrial |
| 305 | *Crotalus* | *oreganus* | Viperidae | Crotalinae | Terrestrial |
| 306 | *Crotalus* | *oreganus* | Viperidae | Crotalinae | Terrestrial |
| 307 | *Crotalus* | *oreganus* | Viperidae | Crotalinae | Terrestrial |
| 308 | *Crotalus* | *viridis* | Viperidae | Crotalinae | Terrestrial |
| 309 | *Crotalus* | *viridis* | Viperidae | Crotalinae | Terrestrial |
| 310 | *Crotalus* | *viridis* | Viperidae | Crotalinae | Terrestrial |
| 311 | *Drymarchon* | *corais* | Colubridae | Colubrinae | Terrestrial |
| 312 | *Drymarchon* | *corais* | Colubridae | Colubrinae | Terrestrial |
| 313 | *Drymarchon* | *corais* | Colubridae | Colubrinae | Terrestrial |
| 314 | *Drymobius* | *margaritiferus* | Colubridae | Colubrinae | Arboreal |
| 315 | *Ahaetulla* | *nasutus* | Colubridae | Ahaetuliinae | Arboreal |
| 316 | *Ahaetulla* | *nasutus* | Colubridae | Ahaetuliinae | Arboreal |
| 317 | *Ahaetulla* | *nasutus* | Colubridae | Ahaetuliinae | Arboreal |
| 318 | *Pantherophis* | *bairdi* | Colubridae | Colubrinae | Arboreal |
| 319 | *Pantherophis* | *bairdi* | Colubridae | Colubrinae | Arboreal |
| 320 | *Pantherophis* | *bairdi* | Colubridae | Colubrinae | Arboreal |
| 321 | *Epicrates* | *cenchria* | Boidae | Boinae | Terrestrial |
| 322 | *Epicrates* | *cenchria* | Boidae | Boinae | Terrestrial |
| 323 | *Epicrates* | *cenchria* | Boidae | Boinae | Terrestrial |
| 324 | *Epicrates* | *cenchria* | Boidae | Boinae | Terrestrial |
| 325 | *Eunectes* | *murinus* | Boidae | Boinae | Semiaquatic |
| 326 | *Farancia* | *abacura* | Colubridae | Dipsadinae | Semiaquatic |
| 327 | *Farancia* | *abacura* | Colubridae | Dipsadinae | Semiaquatic |
| 328 | *Farancia* | *abacura* | Colubridae | Dipsadinae | Semiaquatic |
| 329 | *Haldea* | *striatula* | Colubridae | Natricinae | Semifossorial |
| 330 | *Haldea* | *striatula* | Colubridae | Natricinae | Semifossorial |

**Table 1. (continued)**

| | | | | | |
|---|---|---|---|---|---|
| 331 | *Haldea* | *striatula* | Colubridae | Natricinae | Semifossorial |
| 332 | *Heterodon* | *platirhinos* | Colubridae | Dipsadinae | Semifossorial |
| 333 | *Heterodon* | *platirhinos* | Colubridae | Dipsadinae | Semifossorial |
| 334 | *Heterodon* | *platirhinos* | Colubridae | Dipsadinae | Semifossorial |
| 335 | *Lampropeltis* | *alterna* | Colubridae | Colubrinae | Terrestrial |
| 336 | *Lampropeltis* | *getula* | Colubridae | Colubrinae | Terrestrial |
| 337 | *Lampropeltis* | *getula* | Colubridae | Colubrinae | Terrestrial |
| 338 | *Lampropeltis* | *getula* | Colubridae | Colubrinae | Terrestrial |
| 339 | *Lampropeltis* | *getula* | Colubridae | Colubrinae | Terrestrial |
| 340 | *Lampropeltis* | *getula* | Colubridae | Colubrinae | Terrestrial |
| 341 | *Lampropeltis* | *getula* | Colubridae | Colubrinae | Terrestrial |
| 342 | *Lampropeltis* | *getula* | Colubridae | Colubrinae | Terrestrial |
| 343 | *Lampropeltis* | *zonata* | Colubridae | Colubrinae | Terrestrial |
| 344 | *Lampropeltis* | *zonata* | Colubridae | Colubrinae | Terrestrial |
| 345 | *Lampropeltis* | *zonata* | Colubridae | Colubrinae | Terrestrial |
| 346 | *Laticauda* | *colubrina* | Elapidae | Hydrophiinae | Aquatic |
| 347 | *Rena* | *dulcis* | Leptotyphlopidae | Leptotyphlopinae | Fossorial |
| 348 | *Rena* | *dulcis* | Leptotyphlopidae | Leptotyphlopinae | Fossorial |
| 349 | *Rena* | *dulcis* | Leptotyphlopidae | Leptotyphlopinae | Fossorial |
| 350 | *Rena* | *humilis* | Leptotyphlopidae | Leptotyphlopinae | Fossorial |
| 351 | *Rena* | *humilis* | Leptotyphlopidae | Leptotyphlopinae | Fossorial |
| 352 | *Rena* | *humilis* | Leptotyphlopidae | Leptotyphlopinae | Fossorial |
| 353 | *Lichanura* | *trivirgata* | Charinidae | Charininae | Semifossorial |
| 354 | *Lichanura* | *trivirgata* | Charinidae | Charininae | Semifossorial |
| 355 | *Lichanura* | *trivirgata* | Charinidae | Charininae | Semifossorial |
| 356 | *Lichanura* | *trivirgata* | Charinidae | Charininae | Semifossorial |
| 357 | *Lichanura* | *trivirgata* | Charinidae | Charininae | Semifossorial |
| 358 | *Lichanura* | *trivirgata* | Charinidae | Charininae | Semifossorial |
| 359 | *Masticophis* | *fuliginosus* | Colubridae | Colubrinae | Semiarboreal |
| 360 | *Masticophis* | *fuliginosus* | Colubridae | Colubrinae | Semiarboreal |
| 361 | *Masticophis* | *fuliginosus* | Colubridae | Colubrinae | Semiarboreal |
| 362 | *Masticophis* | *flagellum* | Colubridae | Colubrinae | Semiarboreal |
| 363 | *Masticophis* | *flagellum* | Colubridae | Colubrinae | Semiarboreal |
| 364 | *Masticophis* | *flagellum* | Colubridae | Colubrinae | Semiarboreal |
| 365 | *Masticophis* | *lateralis* | Colubridae | Colubrinae | Semiarboreal |
| 366 | *Masticophis* | *lateralis* | Colubridae | Colubrinae | Semiarboreal |
| 367 | *Masticophis* | *lateralis* | Colubridae | Colubrinae | Semiarboreal |

**Table 1. (continued)**

| | | | | | |
|---|---|---|---|---|---|
| 368 | *Masticophis* | *lateralis* | Colubridae | Colubrinae | Semiarboreal |
| 369 | *Masticophis* | *lateralis* | Colubridae | Colubrinae | Semiarboreal |
| 370 | *Masticophis* | *lateralis* | Colubridae | Colubrinae | Semiarboreal |
| 371 | *Masticophis* | *taeniatus* | Colubridae | Colubrinae | Arboreal |
| 372 | *Masticophis* | *taeniatus* | Colubridae | Colubrinae | Arboreal |
| 373 | *Masticophis* | *taeniatus* | Colubridae | Colubrinae | Arboreal |
| 374 | *Masticophis* | *taeniatus* | Colubridae | Colubrinae | Arboreal |
| 375 | *Micrurus* | *fulvius* | Elapidae | Elapinae | Fossorial |
| 376 | *Micrurus* | *fulvius* | Elapidae | Elapinae | Fossorial |
| 377 | *Micrurus* | *fulvius* | Elapidae | Elapinae | Fossorial |
| 378 | *Micrurus* | *spixii* | Elapidae | Elapinae | Fossorial |
| 379 | *Naja* | *naja* | Elapidae | Elapinae | Terrestrial |
| 380 | *Nerodia* | *cyclopion* | Colubridae | Natricinae | Aquatic |
| 381 | *Nerodia* | *cyclopion* | Colubridae | Natricinae | Aquatic |
| 382 | *Nerodia* | *cyclopion* | Colubridae | Natricinae | Aquatic |
| 383 | *Nerodia* | *taxispilota* | Colubridae | Natricinae | Aquatic |
| 384 | *Nerodia* | *taxispilota* | Colubridae | Natricinae | Aquatic |
| 385 | *Nerodia* | *taxispilota* | Colubridae | Natricinae | Aquatic |
| 386 | *Opheodrys* | *vernalis* | Colubridae | Colubrinae | Semiarboreal |
| 387 | *Opheodrys* | *vernalis* | Colubridae | Colubrinae | Semiarboreal |
| 388 | *Oxybelis* | *sp* | Colubridae | Colubrinae | Arboreal |
| 389 | *Oxybelis* | *sp* | Colubridae | Colubrinae | Arboreal |
| 390 | *Oxybelis* | *sp* | Colubridae | Colubrinae | Arboreal |
| 391 | *Pelamis* | *platura* | Elapidae | Hydrophiinae | Aquatic |
| 392 | *Pelamis* | *platura* | Elapidae | Hydrophiinae | Aquatic |
| 393 | *Pelamis* | *platura* | Elapidae | Hydrophiinae | Aquatic |
| 394 | *Phyllorhynchus* | *browni* | Colubridae | Colubrinae | Fossorial |
| 395 | *Phyllorhynchus* | *browni* | Colubridae | Colubrinae | Fossorial |
| 396 | *Phyllorhynchus* | *browni* | Colubridae | Colubrinae | Fossorial |
| 397 | *Pituophis* | *catenifer* | Colubridae | Colubrinae | Semifossorial |
| 398 | *Pituophis* | *catenifer* | Colubridae | Colubrinae | Semifossorial |
| 399 | *Pituophis* | *catenifer* | Colubridae | Colubrinae | Semifossorial |
| 400 | *Pituophis* | *catenifer* | Colubridae | Colubrinae | Semifossorial |
| 401 | *Pituophis* | *melanoleucas* | Colubridae | Colubrinae | Semifossorial |
| 402 | *Pituophis* | *melanoleucas* | Colubridae | Colubrinae | Semifossorial |
| 403 | *Pituophis* | *melanoleucas* | Colubridae | Colubrinae | Semifossorial |
| 404 | *Pituophis* | *sayi* | Colubridae | Colubrinae | Semifossorial |

**Table 1. (continued)**

| | | | | | |
|---|---|---|---|---|---|
| 405 | *Pituophis* | *sayi* | Colubridae | Colubrinae | Semifossorial |
| 406 | *Pituophis* | *sayi* | Colubridae | Colubrinae | Semifossorial |
| 407 | *Malayopython* | *reticulatus* | Pythonidae | Pythoninae | Terrestrial |
| 408 | *Salvadora* | *lineata* | Colubridae | Colubrinae | Semiarboreal |
| 409 | *Salvadora* | *lineata* | Colubridae | Colubrinae | Semiarboreal |
| 410 | *Salvadora* | *lineata* | Colubridae | Colubrinae | Semiarboreal |
| 411 | *Sistrurus* | *catenatus* | Viperidae | Crotalinae | Terrestrial |
| 412 | *Sistrurus* | *catenatus* | Viperidae | Crotalinae | Terrestrial |
| 413 | *Sistrurus* | *catenatus* | Viperidae | Crotalinae | Terrestrial |
| 414 | *Sistrurus* | *miliarus* | Viperidae | Crotalinae | Terrestrial |
| 415 | *Sistrurus* | *miliarus* | Viperidae | Crotalinae | Terrestrial |
| 416 | *Sistrurus* | *miliarus* | Viperidae | Crotalinae | Terrestrial |
| 417 | *Sistrurus* | *miliarus* | Viperidae | Crotalinae | Terrestrial |
| 418 | *Storeria* | *dekayi* | Colubridae | Natricinae | Terrestrial |
| 419 | *Storeria* | *dekayi* | Colubridae | Natricinae | Terrestrial |
| 420 | *Storeria* | *dekayi* | Colubridae | Natricinae | Terrestrial |
| 421 | *Tantilla* | *coronata* | Colubridae | Colubrinae | Semifossorial |
| 422 | *Tantilla* | *coronata* | Colubridae | Colubrinae | Semifossorial |
| 423 | *Tantilla* | *coronata* | Colubridae | Colubrinae | Semifossorial |
| 424 | *Tantilla* | *planiceps* | Colubridae | Colubrinae | Fossorial |
| 425 | *Tantilla* | *planiceps* | Colubridae | Colubrinae | Fossorial |
| 426 | *Tantilla* | *planiceps* | Colubridae | Colubrinae | Fossorial |
| 427 | *Thamnophis* | *hammondii* | Colubridae | Natricinae | Aquatic |
| 428 | *Thamnophis* | *hammondii* | Colubridae | Natricinae | Aquatic |
| 429 | *Thamnophis* | *hammondii* | Colubridae | Natricinae | Aquatic |
| 430 | *Thamnophis* | *ordinoides* | Colubridae | Natricinae | Terrestrial |
| 431 | *Thamnophis* | *ordinoides* | Colubridae | Natricinae | Terrestrial |
| 432 | *Thamnophis* | *ordinoides* | Colubridae | Natricinae | Terrestrial |
| 433 | *Thamnophis* | *radix* | Colubridae | Natricinae | Terrestrial |
| 434 | *Thamnophis* | *radix* | Colubridae | Natricinae | Terrestrial |
| 435 | *Thamnophis* | *radix* | Colubridae | Natricinae | Terrestrial |
| 436 | *Thamnophis* | *radix* | Colubridae | Natricinae | Terrestrial |
| 437 | *Thamnophis* | *radix* | Colubridae | Natricinae | Terrestrial |
| 438 | *Thamnophis* | *radix* | Colubridae | Natricinae | Terrestrial |
| 439 | *Tropidoclonion* | *lineatum* | Colubridae | Natricinae | Semifossorial |
| 440 | *Tropidoclonion* | *lineatum* | Colubridae | Natricinae | Semifossorial |
| 441 | *Tropidoclonion* | *lineatum* | Colubridae | Natricinae | Semifossorial |

**Table 1. (continued)**

| | | | | | |
|---|---|---|---|---|---|
| 442 | *Crotalus* | *willardi* | Viperidae | Crotalinae | Terrestrial |
| 443 | *Ficimia* | *streckeri* | Colubridae | Colubrinae | Fossorial |
| 444 | *Gyalopion* | *canum* | Colubridae | Colubrinae | Fossorial |
| 445 | *Hypsiglena* | *torquata* | Colubridae | Dipsadinae | Terrestrial |
| 446 | *Nerodia* | *paucimaculata* | Colubridae | Natricinae | Aquatic |
| 447 | *Tantilla* | *gracilis* | Colubridae | Colubrinae | Semifossorial |
| 448 | *Tantilla* | *hobartsmithi* | Colubridae | Colubrinae | Semifossorial |
| 449 | *Tantilla* | *nigriceps* | Colubridae | Colubrinae | Semifossorial |
| 450 | *Agkistrodon* | *bilineatus* | Viperidae | Crotalinae | Semiaquatic |
| 451 | *Agkistrodon* | *bilineatus* | Viperidae | Crotalinae | Semiaquatic |
| 452 | *Agkistrodon* | *piscivorus* | Viperidae | Crotalinae | Semiaquatic |
| 453 | *Agkistrodon* | *piscivorus* | Viperidae | Crotalinae | Semiaquatic |
| 454 | *Bitis* | *arietans* | Viperidae | Viperinae | Terrestrial |
| 455 | *Bitis* | *gabonica* | Viperidae | Viperinae | Terrestrial |
| 456 | *Boa* | *constrictor* | Boidae | Boinae | Semiarboreal |
| 457 | *Carphophis* | *amoenus* | Colubridae | Dipsadinae | Fossorial |
| 458 | *Charina* | *bottae* | Charinidae | Charininae | Semifossorial |
| 459 | *Chrysopelea* | *pelias* | Colubridae | Ahaetuliinae | Arboreal |
| 460 | *Coluber* | *constrictor* | Colubridae | Colubrinae | Semiarboreal |
| 461 | *Crotalus* | *adamanteus* | Viperidae | Crotalinae | Terrestrial |
| 462 | *Chilabothrus* | *angulifer* | Boidae | Boinae | Arboreal |
| 463 | *Farancia* | *abacura* | Colubridae | Dipsadinae | Semiaquatic |
| 464 | *Virginia* | *valeriae* | Colubridae | Natricinae | Semifossorial |
| 465 | *Heterodon* | *platirhinos* | Colubridae | Dipsadinae | Semifossorial |
| 466 | *Lampropeltis* | *calligaster* | Colubridae | Colubrinae | Semifossorial |
| 467 | *Lampropeltis* | *triangulum* | Colubridae | Colubrinae | Semifossorial |
| 468 | *Lampropeltis* | *getula* | Colubridae | Colubrinae | Terrestrial |
| 469 | *Lampropeltis* | *nigra* | Colubridae | Colubrinae | Terrestrial |
| 470 | *Lampropeltis* | *elapsoides* | Colubridae | Colubrinae | Semifossorial |
| 471 | *Leptodeira* | *annulata* | Colubridae | Dipsadinae | Semiaquatic |
| 472 | *Micrurus* | *fulvius* | Elapidae | Elapinae | Fossorial |
| 473 | *Naja* | *naja* | Elapidae | Elapinae | Terrestrial |
| 474 | *Nerodia* | *cyclopion* | Colubridae | Natricinae | Aquatic |
| 475 | *Nerodia* | *erythrogaster* | Colubridae | Natricinae | Semiaquatic |
| 476 | *Clonophis* | *kirtlandi* | Colubridae | Natricinae | Semiaquatic |
| 477 | *Natrix* | *natrix* | Colubridae | Natricinae | Aquatic |
| 478 | *Nerodia* | *rhombifera* | Colubridae | Natricinae | Semiaquatic |

**Table 1. (continued)**

| | | | | | |
|---|---|---|---|---|---|
| 479 | *Nerodia* | *taxispilota* | Colubridae | Natricinae | Aquatic |
| 480 | *Natrix* | *tessellata* | Colubridae | Natricinae | Aquatic |
| 481 | *Opheodrys* | *vernalis* | Colubridae | Colubrinae | Semiarboreal |
| 482 | *Oxybelis* | *aeneus* | Colubridae | Colubrinae | Arboreal |
| 483 | *Oxybelis* | *fulgidus* | Colubridae | Colubrinae | Arboreal |
| 484 | *Phyllorhynchus* | *decurtatus* | Colubridae | Colubrinae | Fossorial |
| 485 | *Pituophis* | *catenifer* | Colubridae | Colubrinae | Semifossorial |
| 486 | *Pituophis* | *melanoleucas* | Colubridae | Colubrinae | Semifossorial |
| 487 | *Pituophis* | *sayi* | Colubridae | Colubrinae | Semifossorial |
| 488 | *Pseudaspis* | *cana* | Lamprophiidae | Pseudaspidinae | Fossorial |
| 489 | *Rhinocheilus* | *lecontei* | Colubridae | Colubrinae | Semifossorial |
| 490 | *Rhadinaea* | *flavilata* | Colubridae | Dipsadinae | Terrestrial |
| 491 | *Liodytes* | *pygaea* | Colubridae | Natricinae | Aquatic |
| 492 | *Sistrurus* | *catenatus* | Viperidae | Crotalinae | Terrestrial |
| 493 | *Sistrurus* | *miliarus* | Viperidae | Crotalinae | Terrestrial |
| 494 | *Sonora* | *semiannulata* | Colubridae | Colubrinae | Semifossorial |
| 495 | *Spilotes* | *pullatus* | Colubridae | Colubrinae | Arboreal |
| 496 | *Storeria* | *dekayi* | Colubridae | Natricinae | Terrestrial |
| 497 | *Thamnophis* | *elegans* | Colubridae | Natricinae | Terrestrial |
| 498 | *Thamnophis* | *marcianus* | Colubridae | Natricinae | Semiaquatic |
| 499 | *Thamnophis* | *ordinoides* | Colubridae | Natricinae | Terrestrial |
| 500 | *Thamnophis* | *radix* | Colubridae | Natricinae | Terrestrial |
| 501 | *Thamnophis* | *sauritus* | Colubridae | Natricinae | Semiaquatic |
| 502 | *Thamnophis* | *sirtalis* | Colubridae | Natricinae | Semiaquatic |
| 503 | *Thamnophis* | *sirtalis* | Colubridae | Natricinae | Semiaquatic |
| 504 | *Virginia* | *valeriae* | Colubridae | Natricinae | Semifossorial |

**Table 2. ANOVA results for families within all groups.**

| | Df | Sum Sq | Mean Sq | F value | Pr(>F) |
|---|---|---|---|---|---|
| PC1 | | | | | |
| Family | 1<0.01 | 236.09 | 23.61 | 43.61 | <0.05 |
| Residuals | 493.00 | 266.90 | 0.54 | | |
| PC2 | | | | | |
| Family | 1<0.01 | 247.44 | 24.74 | 47.73 | <0.05 |
| Residuals | 493.00 | 255.56 | 0.52 | | |
| PC3 | | | | | |
| Family | 1<0.01 | 30.88 | 3.09 | 3.22 | <0.05 |
| Residuals | 493.00 | 472.12 | 0.96 | | |
| PC4 | | | | | |
| Family | 1<0.01 | 103.36 | 10.34 | 12.75 | <0.05 |
| Residuals | 493.00 | 399.64 | 0.81 | | |
| PC5 | | | | | |
| Family | 1<0.01 | 128.18 | 12.82 | 16.86 | <0.05 |
| Residuals | 493.00 | 374.82 | 0.76 | | |
| PC6 | | | | | |
| Family | 1<0.01 | 112.52 | 11.25 | 14.21 | <0.05 |
| Residuals | 493.00 | 390.48 | 0.79 | | |

**Table 3. ANOVA results for subfamilies within all groups.**

| | Df | Sum Sq | Mean Sq | F | Pr(>F) |
|---|---|---|---|---|---|
| PC1 | | | | | |
| Subfamily | 15.00 | 268.88 | 17.93 | 37.36 | <0.05 |
| Residuals | 488.00 | 234.12 | 0.48 | | |
| PC2 | | | | | |
| Subfamily | 15.00 | 279.02 | 18.60 | 40.53 | <0.05 |
| Residuals | 488.00 | 223.98 | 0.46 | | |
| PC3 | | | | | |
| Subfamily | 15.00 | 37.11 | 2.47 | 2.59 | <0.05 |
| Residuals | 488.00 | 465.89 | 0.95 | | |
| PC4 | | | | | |
| Subfamily | 15.00 | 140.33 | 9.36 | 12.59 | <0.05 |
| Residuals | 488.00 | 362.67 | 0.74 | | |
| PC5 | | | | | |
| Subfamily | 15.00 | 223.15 | 14.88 | 25.94 | <0.05 |
| Residuals | 488.00 | 279.85 | 0.57 | | |
| PC6 | | | | | |
| Subfamily | 15.00 | 140.79 | 9.39 | 12.65 | <0.05 |
| Residuals | 488.00 | 362.21 | 0.74 | | |

**Table 4. ANOVA results for genera within all groups.**

| | Df | Sum Sq | Mean Sq | F | Pr(>F) |
|---|---|---|---|---|---|
| PC1 | | | | | |
| Genus | 88.00 | 408.01 | 4.64 | 20.26 | <0.05 |
| Residuals | 415.00 | 94.99 | 0.23 | | |
| PC2 | | | | | |
| Genus | 88.00 | 413.03 | 4.69 | 21.65 | <0.05 |
| Residuals | 415.00 | 89.97 | 0.22 | | |
| PC3 | | | | | |
| Genus | 88.00 | 261.39 | 2.97 | 5.10 | <0.05 |
| Residuals | 415.00 | 241.61 | 0.58 | | |
| PC4 | | | | | |
| Genus | 88.00 | 296.07 | 3.36 | 6.75 | <0.05 |
| Residuals | 415.00 | 206.93 | 0.50 | | |
| PC5 | | | | | |
| Genus | 88.00 | 367.97 | 4.18 | 12.85 | <0.05 |
| Residuals | 415.00 | 135.03 | 0.33 | | |
| PC6 | | | | | |
| Genus | 88.00 | 268.88 | 3.06 | 5.42 | <0.05 |
| Residuals | 415.00 | 234.12 | 0.56 | | |

**Table 5. ANOVA results for species within all groups.**

| | Df | Sum Sq | Mean Sq | F | Pr(>F) |
|---|---|---|---|---|---|
| PC1 | | | | | |
| Species | 188.00 | 450.56 | 2.40 | 14.40 | <0.05 |
| Residuals | 315.00 | 52.44 | 0.17 | | |
| PC2 | | | | | |
| Species | 188.00 | 437.08 | 2.32 | 11.11 | <0.05 |
| Residuals | 315.00 | 65.92 | 0.21 | | |
| PC3 | | | | | |
| Species | 188.00 | 373.55 | 1.99 | 4.84 | <0.05 |
| Residuals | 315.00 | 129.45 | 0.41 | | |
| PC4 | | | | | |
| Species | 188.00 | 377.82 | 2.01 | 5.06 | <0.05 |
| Residuals | 315.00 | 125.18 | 0.40 | | |
| PC5 | | | | | |
| Species | 188.00 | 375.30 | 2.00 | 4.92 | <0.05 |
| Residuals | 315.00 | 127.70 | 0.41 | | |
| PC6 | | | | | |
| Species | 188.00 | 362.78 | 1.93 | 4.34 | <0.05 |
| Residuals | 315.00 | 140.22 | 0.45 | | |

**Table 6. ANOVA results for primary foraging habitats within all groups.**

| | Df | Sum Sq | Mean Sq | F | Pr(>F) |
|---|---|---|---|---|---|
| PC1 | | | | | |
| Substrate | 6.00 | 137.41 | 22.90 | 31.13 | <0.05 |
| Residuals | 497.00 | 365.59 | 0.74 | | |
| PC2 | | | | | |
| Substrate | 6.00 | 98.36 | 16.39 | 20.13 | <0.05 |
| Residuals | 497.00 | 404.64 | 0.81 | | |
| PC3 | | | | | |
| Substrate | 6.00 | 45.35 | 7.56 | 8.21 | <0.05 |
| Residuals | 497.00 | 457.65 | 0.92 | | |
| PC4 | | | | | |
| Substrate | 6.00 | 15.82 | 2.64 | 2.69 | <0.05 |
| Residuals | 497.00 | 487.18 | 0.98 | | |
| PC5 | | | | | |
| Substrate | 6.00 | 119.87 | 19.98 | 25.92 | <0.05 |
| Residuals | 497.00 | 383.13 | 0.77 | | |
| PC6 | | | | | |
| Substrate | 6.00 | 110.39 | 18.40 | 23.29 | <0.05 |
| Residuals | 497.00 | 392.61 | 0.79 | | |

**Table 7. ANOVA results for genera within Crotalinae.**

| | Df | Sum Sq | Mean Sq | F value | Pr(>F) |
|---|---|---|---|---|---|
| PC1 | | | | | |
| Crotalinae | 4 | 30.06 | 7.51 | 10.48 | <0.05 |
| Residuals | 92 | 65.94 | 0.72 | | |
| PC2 | | | | | |
| Crotalinae | 4 | 19.66 | 4.91 | 5.92 | <0.05 |
| Residuals | 92 | 76.34 | 0.83 | | |
| PC3 | | | | | |
| Crotalinae | 4 | 21.01 | 5.25 | 6.44 | <0.05 |
| Residuals | 92 | 74.99 | 0.82 | | |
| PC4 | | | | | |
| Crotalinae | 4 | 22.12 | 5.53 | 6.88 | <0.05 |
| Residuals | 92 | 73.89 | 0.80 | | |
| PC5 | | | | | |
| Crotalinae | 4 | 1.02 | 0.25 | 0.25 | 0.91 |
| Residuals | 92 | 94.98 | 1.03 | | |
| PC6 | | | | | |
| Crotalinae | 4 | 27.00 | 6.75 | 9.00 | <0.05 |
| Residuals | 92 | 69.00 | 0.75 | | |

**Table 8. ANOVA results for species within Crotalinae.**

| | Df | Sum Sq | Mean Sq | F value | Pr(>F) |
|---|---|---|---|---|---|
| PC1 | | | | | |
| Crotalinae | 23.00 | 71.13 | 3.09 | 9.08 | <0.05 |
| Residuals | 73.00 | 24.88 | 0.34 | | |
| PC2 | | | | | |
| Crotalinae | 23.00 | 56.22 | 2.44 | 4.49 | <0.05 |
| Residuals | 73.00 | 39.78 | 0.54 | | |
| PC3 | | | | | |
| Crotalinae | 23.00 | 53.66 | 2.33 | 4.02 | <0.05 |
| Residuals | 73.00 | 42.34 | 0.58 | | |
| PC4 | | | | | |
| Crotalinae | 23.00 | 62.19 | 2.70 | 5.84 | <0.05 |
| Residuals | 73.00 | 33.81 | 0.46 | | |
| PC5 | | | | | |
| Crotalinae | 23.00 | 59.82 | 2.60 | 5.25 | <0.05 |
| Residuals | 73.00 | 36.18 | 0.50 | | |
| PC6 | | | | | |
| Crotalinae | 23.00 | 45.61 | 1.98 | 2.87 | <0.05 |
| Residuals | 73.00 | 50.39 | 0.69 | | |

**Table 9. ANOVA results for primary foraging ecology within Crotalinae.**

| | Df | Sum Sq | Mean Sq | F value | Pr(>F) |
|---|---|---|---|---|---|
| PC1 | | | | | |
| Crotalinae | 2.00 | 19.35 | 9.68 | 11.87 | <0.05 |
| Residuals | 94.00 | 76.65 | 0.82 | | |
| PC2 | | | | | |
| Crotalinae | 2.00 | 0.30 | 0.15 | 0.15 | 0.86 |
| Residuals | 94.00 | 95.70 | 1.02 | | |
| PC3 | | | | | |
| Crotalinae | 2.00 | 6.14 | 3.07 | 3.21 | <0.05 |
| Residuals | 94.00 | 89.86 | 0.96 | | |
| PC4 | | | | | |
| Crotalinae | 2.00 | 16.10 | 8.05 | 9.47 | <0.05 |
| Residuals | 94.00 | 79.90 | 0.85 | | |
| PC5 | | | | | |
| Crotalinae | 2.00 | 6.36 | 3.18 | 3.33 | <0.05 |
| Residuals | 94.00 | 89.64 | 0.95 | | |
| PC6 | | | | | |
| Crotalinae | 2.00 | 19.70 | 9.85 | 12.13 | <0.05 |
| Residuals | 94.00 | 76.31 | 0.81 | | |

**Table 10. Tukey's test results of family-level taxonomy for PCs 1-6 of the all-groups data.**

| PC1 All | diff | lwr | upr | p adj |
|---|---|---|---|---|
| Candoiidae-Boidae | -0.50 | -2.94 | 1.93 | 1.00 |
| Charinidae-Boidae | 1.30 | 0.49 | 2.10 | <0.01 |
| Colubridae-Boidae | 0.88 | 0.35 | 1.42 | <0.01 |
| Elapidae-Boidae | 1.03 | 0.29 | 1.76 | <0.01 |
| Lamprophiidae-Boidae | 0.47 | -1.97 | 2.90 | 1.00 |
| Leptotyphlopidae-Boidae | 3.31 | 2.21 | 4.41 | <0.01 |
| Loxocemidae-Boidae | 0.74 | -1.70 | 3.17 | 1.00 |
| Pythonidae-Boidae | 0.32 | -2.11 | 2.76 | 1.00 |
| Tropidophiidae-Boidae | 0.07 | -1.69 | 1.84 | 1.00 |
| Viperidae-Boidae | -0.61 | -1.18 | -0.04 | 0.03 |
| Charinidae-Candoiidae | 1.80 | -0.66 | 4.26 | 0.39 |
| Colubridae-Candoiidae | 1.38 | -1.00 | 3.76 | 0.73 |
| Elapidae-Candoiidae | 1.53 | -0.91 | 3.96 | 0.63 |
| Lamprophiidae-Candoiidae | 0.97 | -2.40 | 4.33 | 1.00 |
| Leptotyphlopidae-Candoiidae | 3.81 | 1.24 | 6.38 | <0.01 |
| Loxocemidae-Candoiidae | 1.24 | -2.13 | 4.60 | 0.98 |
| Pythonidae-Candoiidae | 0.82 | -2.54 | 4.19 | 1.00 |
| Tropidophiidae-Candoiidae | 0.57 | -2.34 | 3.49 | 1.00 |
| Viperidae-Candoiidae | -0.11 | -2.50 | 2.28 | 1.00 |
| Colubridae-Charinidae | -0.42 | -1.05 | 0.21 | 0.54 |
| Elapidae-Charinidae | -0.27 | -1.08 | 0.53 | 0.99 |
| Lamprophiidae-Charinidae | -0.83 | -3.29 | 1.62 | 0.99 |
| Leptotyphlopidae-Charinidae | 2.01 | 0.86 | 3.16 | <0.01 |
| Loxocemidae-Charinidae | -0.56 | -3.02 | 1.90 | 1.00 |
| Pythonidae-Charinidae | -0.97 | -3.43 | 1.48 | 0.97 |
| Tropidophiidae-Charinidae | -1.22 | -3.02 | 0.57 | 0.50 |
| Viperidae-Charinidae | -1.91 | -2.57 | -1.25 | <0.01 |
| Elapidae-Colubridae | 0.14 | -0.39 | 0.68 | 1.00 |
| Lamprophiidae-Colubridae | -0.42 | -2.80 | 1.97 | 1.00 |
| Leptotyphlopidae-Colubridae | 2.43 | 1.45 | 3.41 | <0.01 |
| Loxocemidae-Colubridae | -0.14 | -2.53 | 2.24 | 1.00 |
| Pythonidae-Colubridae | -0.56 | -2.94 | 1.83 | 1.00 |
| Tropidophiidae-Colubridae | -0.81 | -2.49 | 0.88 | 0.90 |
| Viperidae-Colubridae | -1.49 | -1.76 | -1.22 | <0.01 |
| Lamprophiidae-Elapidae | -0.56 | -3.00 | 1.88 | 1.00 |
| Leptotyphlopidae-Elapidae | 2.28 | 1.18 | 3.38 | <0.01 |

**Table 10. (continued)**

| | | | | |
|---|---|---|---|---|
| Loxocemidae-Elapidae | -0.29 | -2.72 | 2.15 | 1.00 |
| Pythonidae-Elapidae | -0.70 | -3.14 | 1.73 | 1.00 |
| Tropidophiidae-Elapidae | -0.95 | -2.71 | 0.81 | 0.81 |
| Viperidae-Elapidae | -1.64 | -2.21 | -1.06 | <0.01 |
| Leptotyphlopidae-Lamprophiidae | 2.84 | 0.27 | 5.41 | 0.02 |
| Loxocemidae-Lamprophiidae | 0.27 | -3.09 | 3.64 | 1.00 |
| Pythonidae-Lamprophiidae | -0.14 | -3.51 | 3.22 | 1.00 |
| Tropidophiidae-Lamprophiidae | -0.39 | -3.31 | 2.52 | 1.00 |
| Viperidae-Lamprophiidae | -1.08 | -3.47 | 1.32 | 0.93 |
| Loxocemidae-Leptotyphlopidae | -2.57 | -5.14 | <0.01 | 0.05 |
| Pythonidae-Leptotyphlopidae | -2.98 | -5.55 | -0.41 | 0.01 |
| Tropidophiidae-Leptotyphlopidae | -3.23 | -5.18 | -1.29 | <0.01 |
| Viperidae-Leptotyphlopidae | -3.92 | -4.92 | -2.92 | <0.01 |
| Pythonidae-Loxocemidae | -0.41 | -3.78 | 2.95 | 1.00 |
| Tropidophiidae-Loxocemidae | -0.66 | -3.58 | 2.25 | 1.00 |
| Viperidae-Loxocemidae | -1.35 | -3.74 | 1.04 | 0.77 |
| Tropidophiidae-Pythonidae | -0.25 | -3.16 | 2.66 | 1.00 |
| Viperidae-Pythonidae | -0.93 | -3.33 | 1.46 | 0.97 |
| Viperidae-Tropidophiidae | -0.68 | -2.38 | 1.02 | 0.97 |
| PC2 All | | | | |
| Candoiidae-Boidae | 0.39 | -1.99 | 2.77 | 1.00 |
| Charinidae-Boidae | 1.42 | 0.64 | 2.21 | <0.01 |
| Colubridae-Boidae | -1.36 | -1.88 | -0.83 | <0.01 |
| Elapidae-Boidae | -1.04 | -1.76 | -0.33 | <0.01 |
| Lamprophiidae-Boidae | -1.08 | -3.46 | 1.30 | 0.93 |
| Leptotyphlopidae-Boidae | 0.44 | -0.63 | 1.52 | 0.96 |
| Loxocemidae-Boidae | 1.10 | -1.28 | 3.48 | 0.92 |
| Pythonidae-Boidae | -0.06 | -2.44 | 2.33 | 1.00 |
| Tropidophiidae-Boidae | -0.27 | -1.99 | 1.45 | 1.00 |
| Viperidae-Boidae | -0.12 | -0.67 | 0.44 | 1.00 |
| Charinidae-Candoiidae | 1.03 | -1.37 | 3.44 | 0.95 |
| Colubridae-Candoiidae | -1.75 | -4.08 | 0.58 | 0.35 |
| Elapidae-Candoiidae | -1.44 | -3.82 | 0.95 | 0.68 |
| Lamprophiidae-Candoiidae | -1.47 | -4.76 | 1.82 | 0.94 |
| Leptotyphlopidae-Candoiidae | 0.05 | -2.46 | 2.57 | 1.00 |
| Loxocemidae-Candoiidae | 0.71 | -2.59 | 4.00 | 1.00 |
| Pythonidae-Candoiidae | -0.45 | -3.74 | 2.85 | 1.00 |

**Table 10. (continued)**

| | | | | |
|---|---|---|---|---|
| Tropidophiidae-Candoiidae | -0.66 | -3.51 | 2.19 | 1.00 |
| Viperidae-Candoiidae | -0.51 | -2.85 | 1.83 | 1.00 |
| Colubridae-Charinidae | -2.78 | -3.39 | -2.17 | <0.01 |
| Elapidae-Charinidae | -2.47 | -3.25 | -1.68 | <0.01 |
| Lamprophiidae-Charinidae | -2.50 | -4.91 | -0.10 | 0.03 |
| Leptotyphlopidae-Charinidae | -0.98 | -2.10 | 0.15 | 0.16 |
| Loxocemidae-Charinidae | -0.32 | -2.73 | 2.08 | 1.00 |
| Pythonidae-Charinidae | -1.48 | -3.88 | 0.93 | 0.66 |
| Tropidophiidae-Charinidae | -1.69 | -3.45 | 0.06 | 0.07 |
| Viperidae-Charinidae | -1.54 | -2.18 | -0.89 | <0.01 |
| Elapidae-Colubridae | 0.31 | -0.21 | 0.84 | 0.69 |
| Lamprophiidae-Colubridae | 0.28 | -2.05 | 2.61 | 1.00 |
| Leptotyphlopidae-Colubridae | 1.80 | 0.84 | 2.76 | <0.01 |
| Loxocemidae-Colubridae | 2.46 | 0.12 | 4.79 | 0.03 |
| Pythonidae-Colubridae | 1.30 | -1.03 | 3.63 | 0.78 |
| Tropidophiidae-Colubridae | 1.09 | -0.56 | 2.74 | 0.56 |
| Viperidae-Colubridae | 1.24 | 0.98 | 1.51 | <0.01 |
| Lamprophiidae-Elapidae | -0.03 | -2.42 | 2.35 | 1.00 |
| Leptotyphlopidae-Elapidae | 1.49 | 0.41 | 2.57 | <0.01 |
| Loxocemidae-Elapidae | 2.14 | -0.24 | 4.53 | 0.12 |
| Pythonidae-Elapidae | 0.99 | -1.39 | 3.37 | 0.96 |
| Tropidophiidae-Elapidae | 0.77 | -0.95 | 2.50 | 0.93 |
| Viperidae-Elapidae | 0.93 | 0.37 | 1.49 | <0.01 |
| Leptotyphlopidae-Lamprophiidae | 1.52 | -0.99 | 4.04 | 0.68 |
| Loxocemidae-Lamprophiidae | 2.18 | -1.11 | 5.47 | 0.55 |
| Pythonidae-Lamprophiidae | 1.02 | -2.27 | 4.32 | 1.00 |
| Tropidophiidae-Lamprophiidae | 0.81 | -2.04 | 3.66 | 1.00 |
| Viperidae-Lamprophiidae | 0.96 | -1.38 | 3.30 | 0.96 |
| Loxocemidae-Leptotyphlopidae | 0.65 | -1.86 | 3.17 | 1.00 |
| Pythonidae-Leptotyphlopidae | -0.50 | -3.01 | 2.02 | 1.00 |
| Tropidophiidae-Leptotyphlopidae | -0.71 | -2.62 | 1.19 | 0.98 |
| Viperidae-Leptotyphlopidae | -0.56 | -1.54 | 0.42 | 0.75 |
| Pythonidae-Loxocemidae | -1.15 | -4.45 | 2.14 | 0.99 |
| Tropidophiidae-Loxocemidae | -1.37 | -4.22 | 1.48 | 0.90 |
| Viperidae-Loxocemidae | -1.21 | -3.55 | 1.13 | 0.85 |
| Tropidophiidae-Pythonidae | -0.21 | -3.07 | 2.64 | 1.00 |
| Viperidae-Pythonidae | -0.06 | -2.40 | 2.28 | 1.00 |

**Table 10. (continued)**

| | | | | |
|---|---|---|---|---|
| Viperidae-Tropidophiidae | 0.15 | -1.51 | 1.82 | 1.00 |
| PC3 All | | | | |
| Candoiidae-Boidae | -0.18 | -3.42 | 3.06 | 1.00 |
| Charinidae-Boidae | -0.93 | -2.00 | 0.14 | 0.15 |
| Colubridae-Boidae | -0.48 | -1.19 | 0.24 | 0.53 |
| Elapidae-Boidae | -0.40 | -1.38 | 0.57 | 0.96 |
| Lamprophiidae-Boidae | -0.75 | -3.99 | 2.49 | 1.00 |
| Leptotyphlopidae-Boidae | 0.83 | -0.64 | 2.29 | 0.76 |
| Loxocemidae-Boidae | -0.67 | -3.91 | 2.57 | 1.00 |
| Pythonidae-Boidae | -1.42 | -4.66 | 1.82 | 0.94 |
| Tropidophiidae-Boidae | 0.34 | -2.00 | 2.68 | 1.00 |
| Viperidae-Boidae | -0.08 | -0.84 | 0.68 | 1.00 |
| Charinidae-Candoiidae | -0.75 | -4.02 | 2.52 | 1.00 |
| Colubridae-Candoiidae | -0.29 | -3.46 | 2.88 | 1.00 |
| Elapidae-Candoiidae | -0.22 | -3.46 | 3.02 | 1.00 |
| Lamprophiidae-Candoiidae | -0.57 | -5.04 | 3.91 | 1.00 |
| Leptotyphlopidae-Candoiidae | 1.01 | -2.40 | 4.43 | 1.00 |
| Loxocemidae-Candoiidae | -0.48 | -4.96 | 3.99 | 1.00 |
| Pythonidae-Candoiidae | -1.23 | -5.71 | 3.24 | 1.00 |
| Tropidophiidae-Candoiidae | 0.52 | -3.35 | 4.40 | 1.00 |
| Viperidae-Candoiidae | 0.11 | -3.08 | 3.29 | 1.00 |
| Colubridae-Charinidae | 0.46 | -0.38 | 1.29 | 0.80 |
| Elapidae-Charinidae | 0.53 | -0.54 | 1.60 | 0.88 |
| Lamprophiidae-Charinidae | 0.18 | -3.09 | 3.45 | 1.00 |
| Leptotyphlopidae-Charinidae | 1.76 | 0.23 | 3.29 | 0.01 |
| Loxocemidae-Charinidae | 0.27 | -3.00 | 3.54 | 1.00 |
| Pythonidae-Charinidae | -0.48 | -3.75 | 2.78 | 1.00 |
| Tropidophiidae-Charinidae | 1.27 | -1.11 | 3.66 | 0.82 |
| Viperidae-Charinidae | 0.86 | -0.02 | 1.73 | 0.06 |
| Elapidae-Colubridae | 0.07 | -0.64 | 0.78 | 1.00 |
| Lamprophiidae-Colubridae | -0.28 | -3.45 | 2.89 | 1.00 |
| Leptotyphlopidae-Colubridae | 1.31 | <0.01 | 2.61 | 0.05 |
| Loxocemidae-Colubridae | -0.19 | -3.36 | 2.98 | 1.00 |
| Pythonidae-Colubridae | -0.94 | -4.11 | 2.23 | 1.00 |
| Tropidophiidae-Colubridae | 0.82 | -1.43 | 3.06 | 0.98 |
| Viperidae-Colubridae | 0.40 | 0.04 | 0.76 | 0.02 |
| Lamprophiidae-Elapidae | -0.35 | -3.59 | 2.89 | 1.00 |

**Table 10. (continued)**

| | | | | |
|---|---|---|---|---|
| Leptotyphlopidae-Elapidae | 1.23 | -0.23 | 2.70 | 0.19 |
| PC4 All | | | | |
| Candoiidae-Boidae | 0.59 | -2.39 | 3.57 | 1.00 |
| Charinidae-Boidae | -0.62 | -1.60 | 0.37 | 0.63 |
| Colubridae-Boidae | -0.56 | -1.21 | 0.10 | 0.18 |
| Elapidae-Boidae | -1.85 | -2.75 | -0.96 | <0.01 |
| Lamprophiidae-Boidae | -0.45 | -3.43 | 2.53 | 1.00 |
| Leptotyphlopidae-Boidae | 2.02 | 0.67 | 3.37 | <0.01 |
| Loxocemidae-Boidae | -1.49 | -4.47 | 1.49 | 0.88 |
| Pythonidae-Boidae | 0.55 | -2.43 | 3.53 | 1.00 |
| Tropidophiidae-Boidae | -1.75 | -3.91 | 0.40 | 0.24 |
| Viperidae-Boidae | -1.00 | -1.70 | -0.30 | <0.01 |
| Charinidae-Candoiidae | -1.21 | -4.22 | 1.80 | 0.97 |
| Colubridae-Candoiidae | -1.15 | -4.07 | 1.77 | 0.97 |
| Elapidae-Candoiidae | -2.45 | -5.43 | 0.54 | 0.22 |
| Lamprophiidae-Candoiidae | -1.04 | -5.16 | 3.08 | 1.00 |
| Leptotyphlopidae-Candoiidae | 1.43 | -1.72 | 4.57 | 0.93 |
| Loxocemidae-Candoiidae | -2.08 | -6.19 | 2.04 | 0.87 |
| Pythonidae-Candoiidae | -0.04 | -4.16 | 4.08 | 1.00 |
| Tropidophiidae-Candoiidae | -2.34 | -5.91 | 1.22 | 0.56 |
| Viperidae-Candoiidae | -1.59 | -4.52 | 1.33 | 0.80 |
| Colubridae-Charinidae | 0.06 | -0.71 | 0.83 | 1.00 |
| Elapidae-Charinidae | -1.23 | -2.22 | -0.25 | <0.01 |
| Lamprophiidae-Charinidae | 0.17 | -2.83 | 3.18 | 1.00 |
| Leptotyphlopidae-Charinidae | 2.64 | 1.23 | 4.05 | <0.01 |
| Loxocemidae-Charinidae | -0.87 | -3.87 | 2.14 | 1.00 |
| Pythonidae-Charinidae | 1.17 | -1.84 | 4.18 | 0.98 |
| Tropidophiidae-Charinidae | -1.13 | -3.33 | 1.06 | 0.85 |
| Viperidae-Charinidae | -0.38 | -1.19 | 0.42 | 0.91 |
| Elapidae-Colubridae | -1.29 | -1.95 | -0.64 | <0.01 |
| Lamprophiidae-Colubridae | 0.11 | -2.80 | 3.03 | 1.00 |
| Leptotyphlopidae-Colubridae | 2.58 | 1.38 | 3.78 | <0.01 |
| Loxocemidae-Colubridae | -0.93 | -3.84 | 1.99 | 0.99 |
| Pythonidae-Colubridae | 1.11 | -1.81 | 4.02 | 0.98 |
| Tropidophiidae-Colubridae | -1.19 | -3.26 | 0.87 | 0.74 |
| Viperidae-Colubridae | -0.44 | -0.78 | -0.11 | <0.01 |
| Lamprophiidae-Elapidae | 1.41 | -1.57 | 4.39 | 0.91 |

**Table 10. (continued)**

| | | | | |
|---|---|---|---|---|
| Leptotyphlopidae-Elapidae | 3.87 | 2.53 | 5.22 | <0.01 |
| Loxocemidae-Elapidae | 0.37 | -2.61 | 3.35 | 1.00 |
| Pythonidae-Elapidae | 2.40 | -0.58 | 5.38 | 0.25 |
| Tropidophiidae-Elapidae | 0.10 | -2.05 | 2.26 | 1.00 |
| Viperidae-Elapidae | 0.85 | 0.15 | 1.55 | <0.01 |
| Leptotyphlopidae-Lamprophiidae | 2.47 | -0.68 | 5.61 | 0.29 |
| Loxocemidae-Lamprophiidae | -1.04 | -5.16 | 3.08 | 1.00 |
| Pythonidae-Lamprophiidae | 1.00 | -3.12 | 5.11 | 1.00 |
| Tropidophiidae-Lamprophiidae | -1.31 | -4.87 | 2.26 | 0.98 |
| Viperidae-Lamprophiidae | -0.56 | -3.48 | 2.37 | 1.00 |
| Loxocemidae-Leptotyphlopidae | -3.50 | -6.65 | -0.36 | 0.02 |
| Pythonidae-Leptotyphlopidae | -1.47 | -4.61 | 1.68 | 0.92 |
| Tropidophiidae-Leptotyphlopidae | -3.77 | -6.15 | -1.40 | <0.01 |
| Viperidae-Leptotyphlopidae | -3.02 | -4.25 | -1.80 | <0.01 |
| Pythonidae-Loxocemidae | 2.04 | -2.08 | 6.15 | 0.88 |
| Tropidophiidae-Loxocemidae | -0.27 | -3.83 | 3.30 | 1.00 |
| Viperidae-Loxocemidae | 0.48 | -2.44 | 3.41 | 1.00 |
| Tropidophiidae-Pythonidae | -2.30 | -5.87 | 1.26 | 0.59 |
| Viperidae-Pythonidae | -1.55 | -4.48 | 1.37 | 0.83 |
| Viperidae-Tropidophiidae | 0.75 | -1.33 | 2.83 | 0.99 |
| PC5 All | | | | |
| Candoiidae-Boidae | 0.40 | -2.49 | 3.29 | 1.00 |
| Charinidae-Boidae | 0.79 | -0.16 | 1.74 | 0.21 |
| Colubridae-Boidae | 2.07 | 1.43 | 2.70 | <0.01 |
| Elapidae-Boidae | 1.92 | 1.05 | 2.79 | <0.01 |
| Lamprophiidae-Boidae | 1.94 | -0.95 | 4.82 | 0.53 |
| Leptotyphlopidae-Boidae | 2.33 | 1.02 | 3.63 | <0.01 |
| Loxocemidae-Boidae | 0.25 | -2.64 | 3.13 | 1.00 |
| Pythonidae-Boidae | -0.53 | -3.42 | 2.35 | 1.00 |
| Tropidophiidae-Boidae | 1.57 | -0.51 | 3.66 | 0.35 |
| Viperidae-Boidae | 2.28 | 1.61 | 2.96 | <0.01 |
| Charinidae-Candoiidae | 0.39 | -2.52 | 3.30 | 1.00 |
| Colubridae-Candoiidae | 1.67 | -1.16 | 4.49 | 0.71 |
| Elapidae-Candoiidae | 1.52 | -1.37 | 4.41 | 0.83 |
| Lamprophiidae-Candoiidae | 1.54 | -2.45 | 5.52 | 0.98 |
| Leptotyphlopidae-Candoiidae | 1.93 | -1.12 | 4.97 | 0.62 |
| Loxocemidae-Candoiidae | -0.15 | -4.14 | 3.83 | 1.00 |

**Table 10. (continued)**

| | | | | |
|---|---|---|---|---|
| Pythonidae-Candoiidae | -0.93 | -4.92 | 3.05 | 1.00 |
| Tropidophiidae-Candoiidae | 1.17 | -2.28 | 4.63 | 0.99 |
| Viperidae-Candoiidae | 1.88 | -0.95 | 4.72 | 0.54 |
| Colubridae-Charinidae | 1.28 | 0.53 | 2.02 | <0.01 |
| Elapidae-Charinidae | 1.13 | 0.18 | 2.08 | 0.01 |
| Lamprophiidae-Charinidae | 1.15 | -1.77 | 4.06 | 0.97 |
| Leptotyphlopidae-Charinidae | 1.54 | 0.17 | 2.90 | 0.01 |
| Loxocemidae-Charinidae | -0.54 | -3.46 | 2.37 | 1.00 |
| Pythonidae-Charinidae | -1.32 | -4.24 | 1.59 | 0.93 |
| Tropidophiidae-Charinidae | 0.78 | -1.34 | 2.90 | 0.98 |
| Viperidae-Charinidae | 1.49 | 0.71 | 2.28 | <0.01 |
| Elapidae-Colubridae | -0.15 | -0.78 | 0.49 | 1.00 |
| Lamprophiidae-Colubridae | -0.13 | -2.95 | 2.69 | 1.00 |
| Leptotyphlopidae-Colubridae | 0.26 | -0.90 | 1.42 | 1.00 |
| Loxocemidae-Colubridae | -1.82 | -4.64 | 1.00 | 0.59 |
| Pythonidae-Colubridae | -2.60 | -5.43 | 0.22 | 0.10 |
| Tropidophiidae-Colubridae | -0.49 | -2.49 | 1.50 | 1.00 |
| Viperidae-Colubridae | 0.22 | -0.10 | 0.54 | 0.51 |
| Lamprophiidae-Elapidae | 0.02 | -2.87 | 2.90 | 1.00 |
| Leptotyphlopidae-Elapidae | 0.41 | -0.90 | 1.71 | 1.00 |
| Loxocemidae-Elapidae | -1.67 | -4.56 | 1.21 | 0.73 |
| Pythonidae-Elapidae | -2.45 | -5.34 | 0.43 | 0.18 |
| Tropidophiidae-Elapidae | -0.35 | -2.43 | 1.74 | 1.00 |
| Viperidae-Elapidae | 0.37 | -0.31 | 1.04 | 0.81 |
| Leptotyphlopidae-Lamprophiidae | 0.39 | -2.66 | 3.44 | 1.00 |
| Loxocemidae-Lamprophiidae | -1.69 | -5.68 | 2.30 | 0.95 |
| Pythonidae-Lamprophiidae | -2.47 | -6.46 | 1.52 | 0.65 |
| Tropidophiidae-Lamprophiidae | -0.36 | -3.82 | 3.09 | 1.00 |
| Viperidae-Lamprophiidae | 0.35 | -2.49 | 3.18 | 1.00 |
| Loxocemidae-Leptotyphlopidae | -2.08 | -5.13 | 0.97 | 0.50 |
| Pythonidae-Leptotyphlopidae | -2.86 | -5.91 | 0.18 | 0.09 |
| Tropidophiidae-Leptotyphlopidae | -0.76 | -3.06 | 1.55 | 0.99 |
| Viperidae-Leptotyphlopidae | -0.04 | -1.23 | 1.14 | 1.00 |
| Pythonidae-Loxocemidae | -0.78 | -4.77 | 3.21 | 1.00 |
| Tropidophiidae-Loxocemidae | 1.33 | -2.13 | 4.78 | 0.98 |
| Viperidae-Loxocemidae | 2.04 | -0.80 | 4.87 | 0.42 |
| Tropidophiidae-Pythonidae | 2.11 | -1.35 | 5.56 | 0.67 |

**Table 10. (continued)**

| | | | | |
|---|---|---|---|---|
| Viperidae-Pythonidae | 2.82 | -0.01 | 5.65 | 0.05 |
| Viperidae-Tropidophiidae | 0.71 | -1.30 | 2.73 | 0.99 |
| PC6 All | | | | |
| Candoiidae-Boidae | -0.62 | -3.57 | 2.33 | 1.00 |
| Charinidae-Boidae | -0.03 | -1.01 | 0.94 | 1.00 |
| Colubridae-Boidae | -0.87 | -1.52 | -0.22 | <0.01 |
| Elapidae-Boidae | -1.77 | -2.66 | -0.88 | <0.01 |
| Lamprophiidae-Boidae | -0.29 | -3.23 | 2.66 | 1.00 |
| Leptotyphlopidae-Boidae | -2.42 | -3.76 | -1.09 | <0.01 |
| Loxocemidae-Boidae | 0.09 | -2.86 | 3.03 | 1.00 |
| Pythonidae-Boidae | -0.11 | -3.06 | 2.83 | 1.00 |
| Tropidophiidae-Boidae | -1.11 | -3.24 | 1.02 | 0.84 |
| Viperidae-Boidae | -1.67 | -2.36 | -0.98 | <0.01 |
| Charinidae-Candoiidae | 0.59 | -2.38 | 3.56 | 1.00 |
| Colubridae-Candoiidae | -0.25 | -3.13 | 2.63 | 1.00 |
| Elapidae-Candoiidae | -1.15 | -4.10 | 1.80 | 0.97 |
| Lamprophiidae-Candoiidae | 0.33 | -3.74 | 4.40 | 1.00 |
| Leptotyphlopidae-Candoiidae | -1.80 | -4.91 | 1.30 | 0.73 |
| Loxocemidae-Candoiidae | 0.71 | -3.36 | 4.78 | 1.00 |
| Pythonidae-Candoiidae | 0.51 | -3.56 | 4.58 | 1.00 |
| Tropidophiidae-Candoiidae | -0.49 | -4.01 | 3.04 | 1.00 |
| Viperidae-Candoiidae | -1.05 | -3.95 | 1.84 | 0.98 |
| Colubridae-Charinidae | -0.84 | -1.60 | -0.08 | 0.02 |
| Elapidae-Charinidae | -1.74 | -2.71 | -0.76 | <0.01 |
| Lamprophiidae-Charinidae | -0.25 | -3.23 | 2.72 | 1.00 |
| Leptotyphlopidae-Charinidae | -2.39 | -3.78 | -1.00 | <0.01 |
| Loxocemidae-Charinidae | 0.12 | -2.85 | 3.09 | 1.00 |
| Pythonidae-Charinidae | -0.08 | -3.05 | 2.89 | 1.00 |
| Tropidophiidae-Charinidae | -1.08 | -3.24 | 1.09 | 0.88 |
| Viperidae-Charinidae | -1.64 | -2.44 | -0.84 | <0.01 |
| Elapidae-Colubridae | -0.90 | -1.55 | -0.25 | <0.01 |
| Lamprophiidae-Colubridae | 0.58 | -2.30 | 3.47 | 1.00 |
| Leptotyphlopidae-Colubridae | -1.55 | -2.74 | -0.37 | <0.01 |
| Loxocemidae-Colubridae | 0.96 | -1.93 | 3.84 | 0.99 |
| Pythonidae-Colubridae | 0.76 | -2.12 | 3.64 | 1.00 |
| Tropidophiidae-Colubridae | -0.24 | -2.28 | 1.80 | 1.00 |
| Viperidae-Colubridae | -0.80 | -1.13 | -0.47 | <0.01 |

**Table 10. (continued)**

| | | | | |
|---|---|---|---|---|
| Lamprophiidae-Elapidae | 1.48 | -1.46 | 4.43 | 0.87 |
| Leptotyphlopidae-Elapidae | -0.65 | -1.99 | 0.68 | 0.89 |
| Loxocemidae-Elapidae | 1.86 | -1.09 | 4.80 | 0.62 |
| Pythonidae-Elapidae | 1.66 | -1.29 | 4.60 | 0.77 |
| Tropidophiidae-Elapidae | 0.66 | -1.47 | 2.79 | 1.00 |
| Viperidae-Elapidae | 0.10 | -0.59 | 0.79 | 1.00 |
| Leptotyphlopidae-Lamprophiidae | -2.14 | -5.25 | 0.97 | 0.49 |
| Loxocemidae-Lamprophiidae | 0.37 | -3.70 | 4.44 | 1.00 |
| Pythonidae-Lamprophiidae | 0.18 | -3.90 | 4.25 | 1.00 |
| Tropidophiidae-Lamprophiidae | -0.82 | -4.35 | 2.70 | 1.00 |
| Viperidae-Lamprophiidae | -1.39 | -4.28 | 1.51 | 0.90 |
| Loxocemidae-Leptotyphlopidae | 2.51 | -0.60 | 5.62 | 0.25 |
| Pythonidae-Leptotyphlopidae | 2.31 | -0.80 | 5.42 | 0.37 |
| Tropidophiidae-Leptotyphlopidae | 1.31 | -1.04 | 3.66 | 0.77 |
| Viperidae-Leptotyphlopidae | 0.75 | -0.46 | 1.96 | 0.64 |
| Pythonidae-Loxocemidae | -0.20 | -4.27 | 3.87 | 1.00 |
| Tropidophiidae-Loxocemidae | -1.20 | -4.72 | 2.33 | 0.99 |
| Viperidae-Loxocemidae | -1.76 | -4.65 | 1.13 | 0.67 |
| Tropidophiidae-Pythonidae | -1.00 | -4.52 | 2.53 | 1.00 |
| Viperidae-Pythonidae | -1.56 | -4.45 | 1.33 | 0.81 |
| Viperidae-Tropidophiidae | -0.56 | -2.62 | 1.49 | 1.00 |

**Table 11. Tukey's test results of subfamily taxonomy for PCs 1-6 for all groups.**

| PC1 All | diff | lwr | upr | p adj |
|---|---|---|---|---|
| Boinae-Ahaetuliinae | -1.19 | -2.37 | <0.01 | 0.05 |
| Candoiinae-Ahaetuliinae | -1.69 | -4.30 | 0.93 | 0.68 |
| Charininae-Ahaetuliinae | 0.11 | -1.12 | 1.34 | 1.00 |
| Colubrinae-Ahaetuliinae | -0.20 | -1.28 | 0.88 | 1.00 |
| Crotalinae-Ahaetuliinae | -1.79 | -2.88 | -0.69 | <0.01 |
| Dipsadinae-Ahaetuliinae | 0.05 | -1.07 | 1.16 | 1.00 |
| Elapinae-Ahaetuliinae | 0.09 | -1.13 | 1.31 | 1.00 |
| Hydrophiinae-Ahaetuliinae | -0.96 | -2.47 | 0.55 | 0.71 |
| Leptotyphlopinae-Ahaetuliinae | 2.12 | 0.68 | 3.57 | <0.01 |
| Loxoceminae-Ahaetuliinae | -0.45 | -3.06 | 2.16 | 1.00 |
| Natricinae-Ahaetuliinae | -0.77 | -1.87 | 0.33 | 0.53 |
| Pseudaspidinae-Ahaetuliinae | -0.72 | -3.33 | 1.89 | 1.00 |
| Pythoninae-Ahaetuliinae | -0.86 | -3.47 | 1.75 | 1.00 |
| Tropidophiinae-Ahaetuliinae | -1.11 | -3.11 | 0.88 | 0.87 |
| Viperinae-Ahaetuliinae | -2.27 | -4.26 | -0.27 | 0.01 |
| Candoiinae-Boinae | -0.50 | -2.94 | 1.94 | 1.00 |
| Charininae-Boinae | 1.30 | 0.49 | 2.11 | <0.01 |
| Colubrinae-Boinae | 0.99 | 0.44 | 1.53 | <0.01 |
| Crotalinae-Boinae | -0.60 | -1.17 | -0.03 | 0.03 |
| Dipsadinae-Boinae | 1.23 | 0.62 | 1.84 | <0.01 |
| Elapinae-Boinae | 1.28 | 0.48 | 2.07 | <0.01 |
| Hydrophiinae-Boinae | 0.23 | -0.96 | 1.42 | 1.00 |
| Leptotyphlopinae-Boinae | 3.31 | 2.20 | 4.41 | <0.01 |
| Loxoceminae-Boinae | 0.74 | -1.70 | 3.18 | 1.00 |
| Natricinae-Boinae | 0.42 | -0.16 | 1.00 | 0.50 |
| Pseudaspidinae-Boinae | 0.47 | -1.98 | 2.91 | 1.00 |
| Pythoninae-Boinae | 0.32 | -2.12 | 2.77 | 1.00 |
| Tropidophiinae-Boinae | 0.07 | -1.69 | 1.84 | 1.00 |
| Viperinae-Boinae | -1.08 | -2.84 | 0.69 | 0.76 |
| Charininae-Candoiinae | 1.80 | -0.66 | 4.26 | 0.46 |
| Colubrinae-Candoiinae | 1.49 | -0.91 | 3.88 | 0.74 |
| Crotalinae-Candoiinae | -0.10 | -2.50 | 2.30 | 1.00 |
| Dipsadinae-Candoiinae | 1.73 | -0.68 | 4.14 | 0.49 |
| Elapinae-Candoiinae | 1.78 | -0.68 | 4.24 | 0.48 |
| Hydrophiinae-Candoiinae | 0.73 | -1.88 | 3.34 | 1.00 |

**Table 11. (continued)**

| | | | | |
|---|---|---|---|---|
| Leptotyphlopinae-Candoiinae | 3.81 | 1.23 | 6.38 | <0.01 |
| Loxoceminae-Candoiinae | 1.24 | -2.14 | 4.61 | 1.00 |
| Natricinae-Candoiinae | 0.92 | -1.48 | 3.32 | 1.00 |
| Pseudaspidinae-Candoiinae | 0.97 | -2.41 | 4.34 | 1.00 |
| Pythoninae-Candoiinae | 0.82 | -2.55 | 4.20 | 1.00 |
| Tropidophiinae-Candoiinae | 0.57 | -2.35 | 3.50 | 1.00 |
| Viperinae-Candoiinae | -0.58 | -3.50 | 2.34 | 1.00 |
| Colubrinae-Charininae | -0.31 | -0.95 | 0.33 | 0.95 |
| Crotalinae-Charininae | -1.90 | -2.56 | -1.24 | <0.01 |
| Dipsadinae-Charininae | -0.07 | -0.76 | 0.63 | 1.00 |
| Elapinae-Charininae | -0.02 | -0.88 | 0.83 | 1.00 |
| Hydrophiinae-Charininae | -1.07 | -2.30 | 0.16 | 0.18 |
| Leptotyphlopinae-Charininae | 2.01 | 0.86 | 3.16 | <0.01 |
| Loxoceminae-Charininae | -0.56 | -3.03 | 1.90 | 1.00 |
| Natricinae-Charininae | -0.88 | -1.55 | -0.22 | <0.01 |
| Pseudaspidinae-Charininae | -0.83 | -3.30 | 1.63 | 1.00 |
| Pythoninae-Charininae | -0.97 | -3.44 | 1.49 | 0.99 |
| Tropidophiinae-Charininae | -1.22 | -3.02 | 0.57 | 0.59 |
| Viperinae-Charininae | -2.38 | -4.17 | -0.58 | <0.01 |
| Crotalinae-Colubrinae | -1.59 | -1.88 | -1.29 | <0.01 |
| Dipsadinae-Colubrinae | 0.24 | -0.12 | 0.61 | 0.62 |
| Elapinae-Colubrinae | 0.29 | -0.33 | 0.91 | 0.97 |
| Hydrophiinae-Colubrinae | -0.76 | -1.84 | 0.32 | 0.54 |
| Leptotyphlopinae-Colubrinae | 2.32 | 1.33 | 3.31 | <0.01 |
| Loxoceminae-Colubrinae | -0.25 | -2.64 | 2.14 | 1.00 |
| Natricinae-Colubrinae | -0.57 | -0.88 | -0.26 | <0.01 |
| Pseudaspidinae-Colubrinae | -0.52 | -2.91 | 1.87 | 1.00 |
| Pythoninae-Colubrinae | -0.66 | -3.05 | 1.73 | 1.00 |
| Tropidophiinae-Colubrinae | -0.91 | -2.61 | 0.78 | 0.90 |
| Viperinae-Colubrinae | -2.07 | -3.76 | -0.37 | <0.01 |
| Dipsadinae-Crotalinae | 1.83 | 1.43 | 2.23 | <0.01 |
| Elapinae-Crotalinae | 1.88 | 1.23 | 2.52 | <0.01 |
| Hydrophiinae-Crotalinae | 0.83 | -0.27 | 1.92 | 0.40 |
| Leptotyphlopinae-Crotalinae | 3.91 | 2.90 | 4.91 | <0.01 |
| Loxoceminae-Crotalinae | 1.34 | -1.06 | 3.74 | 0.87 |
| Natricinae-Crotalinae | 1.02 | 0.66 | 1.37 | <0.01 |
| Pseudaspidinae-Crotalinae | 1.07 | -1.33 | 3.46 | 0.98 |

**Table 11. (continued)**

| | | | | |
|---|---|---|---|---|
| Pythoninae-Crotalinae | 0.92 | -1.47 | 3.32 | 0.99 |
| Tropidophiinae-Crotalinae | 0.67 | -1.03 | 2.38 | 0.99 |
| Viperinae-Crotalinae | -0.48 | -2.18 | 1.23 | 1.00 |
| Elapinae-Dipsadinae | 0.04 | -0.63 | 0.72 | 1.00 |
| Hydrophiinae-Dipsadinae | -1.00 | -2.12 | 0.11 | 0.13 |
| Leptotyphlopinae-Dipsadinae | 2.08 | 1.05 | 3.10 | <0.01 |
| Loxoceminae-Dipsadinae | -0.49 | -2.90 | 1.91 | 1.00 |
| Natricinae-Dipsadinae | -0.82 | -1.23 | -0.40 | <0.01 |
| Pseudaspidinae-Dipsadinae | -0.76 | -3.17 | 1.64 | 1.00 |
| Pythoninae-Dipsadinae | -0.91 | -3.31 | 1.50 | 1.00 |
| Tropidophiinae-Dipsadinae | -1.16 | -2.87 | 0.56 | 0.61 |
| Viperinae-Dipsadinae | -2.31 | -4.03 | -0.59 | <0.01 |
| Hydrophiinae-Elapinae | -1.05 | -2.27 | 0.17 | 0.19 |
| Leptotyphlopinae-Elapinae | 2.03 | 0.89 | 3.17 | <0.01 |
| Loxoceminae-Elapinae | -0.54 | -3.00 | 1.92 | 1.00 |
| Natricinae-Elapinae | -0.86 | -1.51 | -0.21 | <0.01 |
| Pseudaspidinae-Elapinae | -0.81 | -3.27 | 1.65 | 1.00 |
| Pythoninae-Elapinae | -0.95 | -3.41 | 1.51 | 0.99 |
| Tropidophiinae-Elapinae | -1.20 | -2.99 | 0.59 | 0.61 |
| Viperinae-Elapinae | -2.36 | -4.14 | -0.57 | <0.01 |
| Leptotyphlopinae-Hydrophiinae | 3.08 | 1.63 | 4.52 | <0.01 |
| Loxoceminae-Hydrophiinae | 0.51 | -2.10 | 3.12 | 1.00 |
| Natricinae-Hydrophiinae | 0.19 | -0.91 | 1.28 | 1.00 |
| Pseudaspidinae-Hydrophiinae | 0.24 | -2.38 | 2.85 | 1.00 |
| Pythoninae-Hydrophiinae | 0.10 | -2.52 | 2.71 | 1.00 |
| Tropidophiinae-Hydrophiinae | -0.15 | -2.15 | 1.84 | 1.00 |
| Viperinae-Hydrophiinae | -1.31 | -3.30 | 0.69 | 0.66 |
| Loxoceminae-Leptotyphlopinae | -2.57 | -5.15 | 0.01 | 0.05 |
| Natricinae-Leptotyphlopinae | -2.89 | -3.90 | -1.88 | <0.01 |
| Pseudaspidinae-Leptotyphlopinae | -2.84 | -5.42 | -0.26 | 0.02 |
| Pythoninae-Leptotyphlopinae | -2.98 | -5.56 | -0.41 | 0.01 |
| Tropidophiinae-Leptotyphlopinae | -3.23 | -5.18 | -1.29 | <0.01 |
| Viperinae-Leptotyphlopinae | -4.39 | -6.33 | -2.44 | <0.01 |
| Natricinae-Loxoceminae | -0.32 | -2.72 | 2.08 | 1.00 |

**Table 11. (continued)**

| | | | | |
|---|---|---|---|---|
| Pseudaspidinae-Loxoceminae | -0.27 | -3.64 | 3.10 | 1.00 |
| Pythoninae-Loxoceminae | -0.41 | -3.79 | 2.96 | 1.00 |
| Tropidophiinae-Loxoceminae | -0.66 | -3.58 | 2.26 | 1.00 |
| Viperinae-Loxoceminae | -1.82 | -4.74 | 1.11 | 0.74 |
| Pseudaspidinae-Natricinae | 0.05 | -2.35 | 2.45 | 1.00 |
| Pythoninae-Natricinae | -0.09 | -2.49 | 2.31 | 1.00 |
| Tropidophiinae-Natricinae | -0.34 | -2.05 | 1.36 | 1.00 |
| Viperinae-Natricinae | -1.49 | -3.20 | 0.21 | 0.17 |
| Pythoninae-Pseudaspidinae | -0.14 | -3.52 | 3.23 | 1.00 |
| Tropidophiinae-Pseudaspidinae | -0.39 | -3.31 | 2.53 | 1.00 |
| Viperinae-Pseudaspidinae | -1.55 | -4.47 | 1.38 | 0.91 |
| Tropidophiinae-Pythoninae | -0.25 | -3.17 | 2.67 | 1.00 |
| Viperinae-Pythoninae | -1.40 | -4.33 | 1.52 | 0.96 |
| Viperinae-Tropidophiinae | -1.15 | -3.54 | 1.23 | 0.95 |
| PC2 All | | | | |
| Boinae-Ahaetuliinae | 2.38 | 1.21 | 3.54 | <0.01 |
| Candoiinae-Ahaetuliinae | 2.77 | 0.21 | 5.32 | 0.02 |
| Charininae-Ahaetuliinae | 3.80 | 2.59 | 5.00 | <0.01 |
| Colubrinae-Ahaetuliinae | 0.86 | -0.20 | 1.92 | 0.27 |
| Crotalinae-Ahaetuliinae | 2.25 | 1.18 | 3.32 | <0.01 |
| Dipsadinae-Ahaetuliinae | 1.43 | 0.34 | 2.52 | <0.01 |
| Elapinae-Ahaetuliinae | 1.73 | 0.53 | 2.92 | <0.01 |
| Hydrophiinae-Ahaetuliinae | 0.06 | -1.42 | 1.53 | 1.00 |
| Leptotyphlopinae-Ahaetuliinae | 2.82 | 1.41 | 4.23 | <0.01 |
| Loxoceminae-Ahaetuliinae | 3.47 | 0.92 | 6.03 | <0.01 |
| Natricinae-Ahaetuliinae | 1.16 | 0.08 | 2.23 | 0.02 |
| Pseudaspidinae-Ahaetuliinae | 1.30 | -1.26 | 3.85 | 0.93 |
| Pythoninae-Ahaetuliinae | 2.32 | -0.24 | 4.88 | 0.13 |
| Tropidophiinae-Ahaetuliinae | 2.11 | 0.15 | 4.06 | 0.02 |
| Viperinae-Ahaetuliinae | 2.59 | 0.64 | 4.54 | <0.01 |
| Candoiinae-Boinae | 0.39 | -2.00 | 2.78 | 1.00 |
| Charininae-Boinae | 1.42 | 0.63 | 2.21 | <0.01 |
| Colubrinae-Boinae | -1.51 | -2.05 | -0.98 | <0.01 |
| Crotalinae-Boinae | -0.12 | -0.68 | 0.44 | 1.00 |
| Dipsadinae-Boinae | -0.95 | -1.55 | -0.35 | <0.01 |
| Elapinae-Boinae | -0.65 | -1.42 | 0.13 | 0.23 |

**Table 11. (continued)**

| | | | | |
|---|---|---|---|---|
| Hydrophiinae-Boinae | -2.32 | -3.48 | -1.16 | <0.01 |
| Leptotyphlopinae-Boinae | 0.44 | -0.64 | 1.52 | 0.99 |
| Loxoceminae-Boinae | 1.10 | -1.29 | 3.49 | 0.97 |
| Natricinae-Boinae | -1.22 | -1.79 | -0.65 | <0.01 |
| Pseudaspidinae-Boinae | -1.08 | -3.47 | 1.31 | 0.97 |
| Pythoninae-Boinae | -0.06 | -2.44 | 2.33 | 1.00 |
| Tropidophiinae-Boinae | -0.27 | -2.00 | 1.46 | 1.00 |
| Viperinae-Boinae | 0.21 | -1.51 | 1.94 | 1.00 |
| Charininae-Candoiinae | 1.03 | -1.38 | 3.44 | 0.99 |
| Colubrinae-Candoiinae | -1.91 | -4.25 | 0.43 | 0.27 |
| Crotalinae-Candoiinae | -0.51 | -2.86 | 1.83 | 1.00 |
| Dipsadinae-Candoiinae | -1.34 | -3.69 | 1.02 | 0.85 |
| Elapinae-Candoiinae | -1.04 | -3.44 | 1.37 | 0.98 |
| Hydrophiinae-Candoiinae | -2.71 | -5.26 | -0.15 | 0.03 |
| Leptotyphlopinae-Candoiinae | 0.05 | -2.47 | 2.57 | 1.00 |
| Loxoceminae-Candoiinae | 0.71 | -2.59 | 4.01 | 1.00 |
| Natricinae-Candoiinae | -1.61 | -3.96 | 0.74 | 0.58 |
| Pseudaspidinae-Candoiinae | -1.47 | -4.77 | 1.83 | 0.98 |
| Pythoninae-Candoiinae | -0.45 | -3.75 | 2.85 | 1.00 |
| Tropidophiinae-Candoiinae | -0.66 | -3.52 | 2.20 | 1.00 |
| Viperinae-Candoiinae | -0.18 | -3.03 | 2.68 | 1.00 |
| Colubrinae-Charininae | -2.94 | -3.56 | -2.31 | <0.01 |
| Crotalinae-Charininae | -1.54 | -2.19 | -0.90 | <0.01 |
| Dipsadinae-Charininae | -2.37 | -3.05 | -1.69 | <0.01 |
| Elapinae-Charininae | -2.07 | -2.91 | -1.23 | <0.01 |
| Hydrophiinae-Charininae | -3.74 | -4.94 | -2.53 | <0.01 |
| Leptotyphlopinae-Charininae | -0.98 | -2.11 | 0.15 | 0.18 |
| Loxoceminae-Charininae | -0.32 | -2.73 | 2.09 | 1.00 |
| Natricinae-Charininae | -2.64 | -3.29 | -1.99 | <0.01 |
| Pseudaspidinae-Charininae | -2.50 | -4.91 | -0.09 | 0.03 |
| Pythoninae-Charininae | -1.48 | -3.89 | 0.93 | 0.76 |
| Tropidophiinae-Charininae | -1.69 | -3.45 | 0.06 | 0.07 |
| Viperinae-Charininae | -1.21 | -2.96 | 0.55 | 0.57 |
| Crotalinae-Colubrinae | 1.39 | 1.10 | 1.68 | <0.01 |
| Dipsadinae-Colubrinae | 0.57 | 0.21 | 0.92 | <0.01 |
| Elapinae-Colubrinae | 0.87 | 0.26 | 1.47 | <0.01 |
| Hydrophiinae-Colubrinae | -0.80 | -1.86 | 0.25 | 0.39 |

**Table 11. (continued)**

| | | | | |
|---|---|---|---|---|
| Leptotyphlopinae-Colubrinae | 1.96 | 0.99 | 2.93 | <0.01 |
| Loxoceminae-Colubrinae | 2.61 | 0.27 | 4.95 | 0.01 |
| Natricinae-Colubrinae | 0.30 | -0.01 | 0.60 | 0.06 |
| Pseudaspidinae-Colubrinae | 0.44 | -1.90 | 2.77 | 1.00 |
| Pythoninae-Colubrinae | 1.46 | -0.88 | 3.80 | 0.73 |
| Tropidophiinae-Colubrinae | 1.24 | -0.41 | 2.90 | 0.41 |
| Viperinae-Colubrinae | 1.73 | 0.07 | 3.39 | 0.03 |
| Dipsadinae-Crotalinae | -0.83 | -1.22 | -0.43 | <0.01 |
| Elapinae-Crotalinae | -0.52 | -1.15 | 0.11 | 0.23 |
| Hydrophiinae-Crotalinae | -2.19 | -3.26 | -1.12 | <0.01 |
| Leptotyphlopinae-Crotalinae | 0.57 | -0.42 | 1.55 | 0.83 |
| Loxoceminae-Crotalinae | 1.22 | -1.12 | 3.57 | 0.92 |
| Natricinae-Crotalinae | -1.10 | -1.44 | -0.75 | <0.01 |
| Pseudaspidinae-Crotalinae | -0.96 | -3.30 | 1.39 | 0.99 |
| Pythoninae-Crotalinae | 0.07 | -2.28 | 2.41 | 1.00 |
| Tropidophiinae-Crotalinae | -0.15 | -1.81 | 1.52 | 1.00 |
| Viperinae-Crotalinae | 0.34 | -1.33 | 2.00 | 1.00 |
| Elapinae-Dipsadinae | 0.30 | -0.36 | 0.96 | 0.97 |
| Hydrophiinae-Dipsadinae | -1.37 | -2.46 | -0.28 | <0.01 |
| Leptotyphlopinae-Dipsadinae | 1.39 | 0.39 | 2.40 | <0.01 |
| Loxoceminae-Dipsadinae | 2.05 | -0.31 | 4.40 | 0.18 |
| Natricinae-Dipsadinae | -0.27 | -0.67 | 0.13 | 0.61 |
| Pseudaspidinae-Dipsadinae | -0.13 | -2.49 | 2.22 | 1.00 |
| Pythoninae-Dipsadinae | 0.89 | -1.46 | 3.25 | 1.00 |
| Tropidophiinae-Dipsadinae | 0.68 | -1.00 | 2.36 | 0.99 |
| Viperinae-Dipsadinae | 1.16 | -0.52 | 2.84 | 0.56 |
| Hydrophiinae-Elapinae | -1.67 | -2.87 | -0.47 | <0.01 |
| Leptotyphlopinae-Elapinae | 1.09 | -0.03 | 2.21 | 0.06 |
| Loxoceminae-Elapinae | 1.75 | -0.66 | 4.15 | 0.47 |
| Natricinae-Elapinae | -0.57 | -1.21 | 0.06 | 0.13 |
| Pseudaspidinae-Elapinae | -0.43 | -2.84 | 1.97 | 1.00 |
| Pythoninae-Elapinae | 0.59 | -1.81 | 3.00 | 1.00 |
| Tropidophiinae-Elapinae | 0.38 | -1.37 | 2.13 | 1.00 |
| Viperinae-Elapinae | 0.86 | -0.89 | 2.61 | 0.95 |
| Leptotyphlopinae-Hydrophiinae | 2.76 | 1.35 | 4.17 | <0.01 |
| Loxoceminae-Hydrophiinae | 3.42 | 0.86 | 5.97 | <0.01 |

**Table 11. (continued)**

| | | | | |
|---|---|---|---|---|
| Natricinae-Hydrophiinae | 1.10 | 0.03 | 2.17 | 0.04 |
| Pseudaspidinae-Hydrophiinae | 1.24 | -1.32 | 3.79 | 0.95 |
| Pythoninae-Hydrophiinae | 2.26 | -0.29 | 4.82 | 0.15 |
| Tropidophiinae-Hydrophiinae | 2.05 | 0.09 | 4.00 | 0.03 |
| Viperinae-Hydrophiinae | 2.53 | 0.58 | 4.48 | <0.01 |
| Loxoceminae-Leptotyphlopinae | 0.65 | -1.87 | 3.18 | 1.00 |
| Natricinae-Leptotyphlopinae | -1.66 | -2.65 | -0.68 | <0.01 |
| Pseudaspidinae-Leptotyphlopinae | -1.52 | -4.04 | 1.00 | 0.78 |
| Pythoninae-Leptotyphlopinae | -0.50 | -3.02 | 2.02 | 1.00 |
| Tropidophiinae-Leptotyphlopinae | -0.71 | -2.62 | 1.19 | 1.00 |
| Viperinae-Leptotyphlopinae | -0.23 | -2.13 | 1.68 | 1.00 |
| Natricinae-Loxoceminae | -2.32 | -4.66 | 0.03 | 0.06 |
| Pseudaspidinae-Loxoceminae | -2.18 | -5.48 | 1.12 | 0.64 |
| Pythoninae-Loxoceminae | -1.15 | -4.45 | 2.15 | 1.00 |
| Tropidophiinae-Loxoceminae | -1.37 | -4.23 | 1.49 | 0.96 |
| Viperinae-Loxoceminae | -0.88 | -3.74 | 1.97 | 1.00 |
| Pseudaspidinae-Natricinae | 0.14 | -2.21 | 2.49 | 1.00 |
| Pythoninae-Natricinae | 1.16 | -1.18 | 3.51 | 0.94 |
| Tropidophiinae-Natricinae | 0.95 | -0.72 | 2.62 | 0.85 |
| Viperinae-Natricinae | 1.43 | -0.24 | 3.10 | 0.19 |
| Pythoninae-Pseudaspidinae | 1.02 | -2.28 | 4.32 | 1.00 |
| Tropidophiinae-Pseudaspidinae | 0.81 | -2.05 | 3.67 | 1.00 |
| Viperinae-Pseudaspidinae | 1.29 | -1.56 | 4.15 | 0.97 |
| Tropidophiinae-Pythoninae | -0.21 | -3.07 | 2.64 | 1.00 |
| Viperinae-Pythoninae | 0.27 | -2.59 | 3.13 | 1.00 |
| Viperinae-Tropidophiinae | 0.49 | -1.85 | 2.82 | 1.00 |
| PC3 All | | | | |
| Boinae-Ahaetuliinae | -0.04 | -1.71 | 1.64 | 1.00 |
| Candoiinae-Ahaetuliinae | -0.22 | -3.91 | 3.46 | 1.00 |
| Charininae-Ahaetuliinae | -0.97 | -2.71 | 0.76 | 0.86 |
| Colubrinae-Ahaetuliinae | -0.47 | -1.99 | 1.06 | 1.00 |
| Crotalinae-Ahaetuliinae | -0.09 | -1.64 | 1.45 | 1.00 |
| Dipsadinae-Ahaetuliinae | -0.68 | -2.25 | 0.89 | 0.98 |
| Elapinae-Ahaetuliinae | -0.52 | -2.24 | 1.20 | 1.00 |

**Table 11. (continued)**

| | | | | |
|---|---|---|---|---|
| Hydrophiinae-Ahaetuliinae | -0.20 | -2.33 | 1.93 | 1.00 |
| Leptotyphlopinae-Ahaetuliinae | 0.79 | -1.25 | 2.83 | 0.99 |
| Loxoceminae-Ahaetuliinae | -0.71 | -4.39 | 2.98 | 1.00 |
| Natricinae-Ahaetuliinae | -0.55 | -2.10 | 1.00 | 1.00 |
| Pseudaspidinae-Ahaetuliinae | -0.79 | -4.48 | 2.89 | 1.00 |
| Pythoninae-Ahaetuliinae | -1.46 | -5.14 | 2.23 | 0.99 |
| Tropidophiinae-Ahaetuliinae | 0.30 | -2.52 | 3.12 | 1.00 |
| Viperinae-Ahaetuliinae | -1.22 | -4.04 | 1.59 | 0.98 |
| Candoiinae-Boinae | -0.18 | -3.63 | 3.26 | 1.00 |
| Charininae-Boinae | -0.93 | -2.07 | 0.20 | 0.26 |
| Colubrinae-Boinae | -0.43 | -1.20 | 0.35 | 0.87 |
| Crotalinae-Boinae | -0.06 | -0.87 | 0.75 | 1.00 |
| Dipsadinae-Boinae | -0.64 | -1.50 | 0.22 | 0.43 |
| Elapinae-Boinae | -0.48 | -1.60 | 0.64 | 0.98 |
| Hydrophiinae-Boinae | -0.16 | -1.84 | 1.51 | 1.00 |
| Leptotyphlopinae-Boinae | 0.83 | -0.73 | 2.39 | 0.90 |
| Loxoceminae-Boinae | -0.67 | -4.11 | 2.78 | 1.00 |
| Natricinae-Boinae | -0.51 | -1.33 | 0.31 | 0.74 |
| Pseudaspidinae-Boinae | -0.75 | -4.20 | 2.69 | 1.00 |
| Pythoninae-Boinae | -1.42 | -4.86 | 2.03 | 0.99 |
| Tropidophiinae-Boinae | 0.34 | -2.15 | 2.83 | 1.00 |
| Viperinae-Boinae | -1.18 | -3.67 | 1.31 | 0.96 |
| Charininae-Candoiinae | -0.75 | -4.23 | 2.73 | 1.00 |
| Colubrinae-Candoiinae | -0.24 | -3.62 | 3.13 | 1.00 |
| Crotalinae-Candoiinae | 0.13 | -3.25 | 3.51 | 1.00 |
| Dipsadinae-Candoiinae | -0.46 | -3.85 | 2.94 | 1.00 |
| Elapinae-Candoiinae | -0.30 | -3.77 | 3.17 | 1.00 |
| Hydrophiinae-Candoiinae | 0.02 | -3.67 | 3.71 | 1.00 |
| Leptotyphlopinae-Candoiinae | 1.01 | -2.62 | 4.65 | 1.00 |
| Loxoceminae-Candoiinae | -0.48 | -5.24 | 4.28 | 1.00 |
| Natricinae-Candoiinae | -0.32 | -3.71 | 3.06 | 1.00 |
| Pseudaspidinae-Candoiinae | -0.57 | -5.33 | 4.19 | 1.00 |
| Pythoninae-Candoiinae | -1.23 | -5.99 | 3.52 | 1.00 |
| Tropidophiinae-Candoiinae | 0.52 | -3.60 | 4.64 | 1.00 |
| Viperinae-Candoiinae | -1.00 | -5.12 | 3.12 | 1.00 |
| Colubrinae-Charininae | 0.51 | -0.40 | 1.41 | 0.86 |

**Table 11. (continued)**

| | | | | |
|---|---|---|---|---|
| Crotalinae-Charininae | 0.88 | -0.06 | 1.81 | 0.09 |
| Dipsadinae-Charininae | 0.29 | -0.69 | 1.27 | 1.00 |
| Elapinae-Charininae | 0.45 | -0.76 | 1.66 | 1.00 |
| Hydrophiinae-Charininae | 0.77 | -0.97 | 2.51 | 0.98 |
| Leptotyphlopinae-Charininae | 1.76 | 0.14 | 3.39 | 0.02 |
| Loxoceminae-Charininae | 0.27 | -3.21 | 3.74 | 1.00 |
| Natricinae-Charininae | 0.43 | -0.52 | 1.37 | 0.97 |
| Pseudaspidinae-Charininae | 0.18 | -3.29 | 3.66 | 1.00 |
| Pythoninae-Charininae | -0.48 | -3.96 | 2.99 | 1.00 |
| Tropidophiinae-Charininae | 1.27 | -1.26 | 3.81 | 0.94 |
| Viperinae-Charininae | -0.25 | -2.78 | 2.28 | 1.00 |
| Crotalinae-Colubrinae | 0.37 | -0.05 | 0.79 | 0.15 |
| Dipsadinae-Colubrinae | -0.21 | -0.73 | 0.30 | 0.99 |
| Elapinae-Colubrinae | -0.05 | -0.93 | 0.82 | 1.00 |
| Hydrophiinae-Colubrinae | 0.26 | -1.26 | 1.79 | 1.00 |
| Leptotyphlopinae-Colubrinae | 1.26 | -0.14 | 2.65 | 0.13 |
| Loxoceminae-Colubrinae | -0.24 | -3.61 | 3.13 | 1.00 |
| Natricinae-Colubrinae | -0.08 | -0.52 | 0.35 | 1.00 |
| Pseudaspidinae-Colubrinae | -0.33 | -3.70 | 3.05 | 1.00 |
| Pythoninae-Colubrinae | -0.99 | -4.37 | 2.38 | 1.00 |
| Tropidophiinae-Colubrinae | 0.77 | -1.63 | 3.16 | 1.00 |
| Viperinae-Colubrinae | -0.76 | -3.15 | 1.64 | 1.00 |
| Dipsadinae-Crotalinae | -0.58 | -1.15 | -0.02 | 0.04 |
| Elapinae-Crotalinae | -0.42 | -1.33 | 0.48 | 0.97 |
| Hydrophiinae-Crotalinae | -0.11 | -1.65 | 1.44 | 1.00 |
| Leptotyphlopinae-Crotalinae | 0.89 | -0.53 | 2.30 | 0.73 |
| Loxoceminae-Crotalinae | -0.61 | -3.99 | 2.77 | 1.00 |
| Natricinae-Crotalinae | -0.45 | -0.95 | 0.04 | 0.12 |
| Pseudaspidinae-Crotalinae | -0.70 | -4.08 | 2.69 | 1.00 |
| Pythoninae-Crotalinae | -1.36 | -4.75 | 2.02 | 0.99 |
| Tropidophiinae-Crotalinae | 0.40 | -2.01 | 2.80 | 1.00 |
| Viperinae-Crotalinae | -1.13 | -3.53 | 1.28 | 0.96 |
| Elapinae-Dipsadinae | 0.16 | -0.80 | 1.12 | 1.00 |
| Hydrophiinae-Dipsadinae | 0.48 | -1.10 | 2.05 | 1.00 |
| Leptotyphlopinae-Dipsadinae | 1.47 | 0.02 | 2.92 | 0.04 |
| Loxoceminae-Dipsadinae | -0.03 | -3.42 | 3.37 | 1.00 |
| Natricinae-Dipsadinae | 0.13 | -0.45 | 0.71 | 1.00 |

**Table 11. (continued)**

| | | | | |
|---|---|---|---|---|
| Pseudaspidinae-Dipsadinae | -0.11 | -3.51 | 3.28 | 1.00 |
| Pythoninae-Dipsadinae | -0.78 | -4.17 | 2.62 | 1.00 |
| Tropidophiinae-Dipsadinae | 0.98 | -1.44 | 3.40 | 0.99 |
| Viperinae-Dipsadinae | -0.54 | -2.97 | 1.88 | 1.00 |
| Hydrophiinae-Elapinae | 0.32 | -1.41 | 2.04 | 1.00 |
| Leptotyphlopinae-Elapinae | 1.31 | -0.30 | 2.92 | 0.27 |
| Loxoceminae-Elapinae | -0.19 | -3.66 | 3.28 | 1.00 |
| Natricinae-Elapinae | -0.03 | -0.94 | 0.89 | 1.00 |
| Pseudaspidinae-Elapinae | -0.27 | -3.74 | 3.20 | 1.00 |
| Pythoninae-Elapinae | -0.94 | -4.41 | 2.53 | 1.00 |
| Tropidophiinae-Elapinae | 0.82 | -1.70 | 3.34 | 1.00 |
| Viperinae-Elapinae | -0.70 | -3.23 | 1.82 | 1.00 |
| Leptotyphlopinae-Hydrophiinae | 0.99 | -1.04 | 3.03 | 0.95 |
| Loxoceminae-Hydrophiinae | -0.50 | -4.19 | 3.18 | 1.00 |
| Natricinae-Hydrophiinae | -0.35 | -1.89 | 1.20 | 1.00 |
| Pseudaspidinae-Hydrophiinae | -0.59 | -4.28 | 3.10 | 1.00 |
| Pythoninae-Hydrophiinae | -1.26 | -4.94 | 2.43 | 1.00 |
| Tropidophiinae-Hydrophiinae | 0.50 | -2.31 | 3.32 | 1.00 |
| Viperinae-Hydrophiinae | -1.02 | -3.84 | 1.79 | 1.00 |
| Loxoceminae-Leptotyphlopinae | -1.50 | -5.13 | 2.14 | 0.99 |
| Natricinae-Leptotyphlopinae | -1.34 | -2.76 | 0.08 | 0.09 |
| Pseudaspidinae-Leptotyphlopinae | -1.58 | -5.22 | 2.05 | 0.98 |
| Pythoninae-Leptotyphlopinae | -2.25 | -5.88 | 1.39 | 0.74 |
| Tropidophiinae-Leptotyphlopinae | -0.49 | -3.24 | 2.26 | 1.00 |
| Viperinae-Leptotyphlopinae | -2.01 | -4.76 | 0.73 | 0.46 |
| Natricinae-Loxoceminae | 0.16 | -3.23 | 3.54 | 1.00 |
| Pseudaspidinae-Loxoceminae | -0.09 | -4.85 | 4.67 | 1.00 |
| Pythoninae-Loxoceminae | -0.75 | -5.51 | 4.01 | 1.00 |
| Tropidophiinae-Loxoceminae | 1.01 | -3.12 | 5.13 | 1.00 |
| Viperinae-Loxoceminae | -0.52 | -4.64 | 3.60 | 1.00 |
| Pseudaspidinae-Natricinae | -0.24 | -3.63 | 3.14 | 1.00 |
| Pythoninae-Natricinae | -0.91 | -4.29 | 2.47 | 1.00 |
| Tropidophiinae-Natricinae | 0.85 | -1.56 | 3.25 | 1.00 |
| Viperinae-Natricinae | -0.68 | -3.08 | 1.73 | 1.00 |

**Table 11. (continued)**

| | | | | |
|---|---|---|---|---|
| Pythoninae-Pseudaspidinae | -0.67 | -5.42 | 4.09 | 1.00 |
| Tropidophiinae-Pseudaspidinae | 1.09 | -3.03 | 5.21 | 1.00 |
| Viperinae-Pseudaspidinae | -0.43 | -4.55 | 3.69 | 1.00 |
| Tropidophiinae-Pythoninae | 1.76 | -2.36 | 5.88 | 0.99 |
| Viperinae-Pythoninae | 0.23 | -3.89 | 4.36 | 1.00 |
| Viperinae-Tropidophiinae | -1.52 | -4.89 | 1.84 | 0.97 |
| PC4 All | | | | |
| Boinae-Ahaetuliinae | 1.83 | 0.36 | 3.31 | <0.01 |
| Candoiinae-Ahaetuliinae | 2.42 | -0.83 | 5.68 | 0.42 |
| Charininae-Ahaetuliinae | 1.21 | -0.32 | 2.75 | 0.32 |
| Colubrinae-Ahaetuliinae | 1.30 | -0.05 | 2.64 | 0.07 |
| Crotalinae-Ahaetuliinae | 0.80 | -0.56 | 2.16 | 0.81 |
| Dipsadinae-Ahaetuliinae | 1.70 | 0.31 | 3.09 | <0.01 |
| Elapinae-Ahaetuliinae | -0.39 | -1.91 | 1.13 | 1.00 |
| Hydrophiinae-Ahaetuliinae | 1.15 | -0.73 | 3.03 | 0.76 |
| Leptotyphlopinae-Ahaetuliinae | 3.85 | 2.05 | 5.65 | <0.01 |
| Loxoceminae-Ahaetuliinae | 0.35 | -2.90 | 3.60 | 1.00 |
| Natricinae-Ahaetuliinae | 1.03 | -0.34 | 2.39 | 0.41 |
| Pseudaspidinae-Ahaetuliinae | 1.39 | -1.87 | 4.64 | 0.99 |
| Pythoninae-Ahaetuliinae | 2.38 | -0.87 | 5.64 | 0.46 |
| Tropidophiinae-Ahaetuliinae | 0.08 | -2.40 | 2.56 | 1.00 |
| Viperinae-Ahaetuliinae | 2.29 | -0.19 | 4.78 | 0.11 |
| Candoiinae-Boinae | 0.59 | -2.45 | 3.63 | 1.00 |
| Charininae-Boinae | -0.62 | -1.62 | 0.38 | 0.75 |
| Colubrinae-Boinae | -0.54 | -1.22 | 0.15 | 0.33 |
| Crotalinae-Boinae | -1.03 | -1.75 | -0.32 | <0.01 |
| Dipsadinae-Boinae | -0.13 | -0.89 | 0.63 | 1.00 |
| Elapinae-Boinae | -2.22 | -3.20 | -1.23 | <0.01 |
| Hydrophiinae-Boinae | -0.68 | -2.16 | 0.79 | 0.97 |
| Leptotyphlopinae-Boinae | 2.02 | 0.65 | 3.39 | <0.01 |
| Loxoceminae-Boinae | -1.49 | -4.52 | 1.55 | 0.95 |
| Natricinae-Boinae | -0.81 | -1.53 | -0.08 | 0.01 |
| Pseudaspidinae-Boinae | -0.45 | -3.48 | 2.59 | 1.00 |
| Pythoninae-Boinae | 0.55 | -2.49 | 3.59 | 1.00 |
| Tropidophiinae-Boinae | -1.75 | -3.95 | 0.44 | 0.30 |
| Viperinae-Boinae | 0.46 | -1.74 | 2.66 | 1.00 |

**Table 11. (continued)**

| | | | | |
|---|---|---|---|---|
| Charininae-Candoiinae | -1.21 | -4.28 | 1.86 | 0.99 |
| Colubrinae-Candoiinae | -1.13 | -4.10 | 1.85 | 1.00 |
| Crotalinae-Candoiinae | -1.62 | -4.61 | 1.36 | 0.89 |
| Dipsadinae-Candoiinae | -0.72 | -3.72 | 2.27 | 1.00 |
| Elapinae-Candoiinae | -2.81 | -5.87 | 0.25 | 0.11 |
| Hydrophiinae-Candoiinae | -1.27 | -4.53 | 1.98 | 0.99 |
| Leptotyphlopinae-Candoiinae | 1.43 | -1.78 | 4.64 | 0.98 |
| Loxoceminae-Candoiinae | -2.08 | -6.28 | 2.12 | 0.95 |
| Natricinae-Candoiinae | -1.40 | -4.38 | 1.59 | 0.97 |
| Pseudaspidinae-Candoiinae | -1.04 | -5.24 | 3.16 | 1.00 |
| Pythoninae-Candoiinae | -0.04 | -4.24 | 4.16 | 1.00 |
| Tropidophiinae-Candoiinae | -2.34 | -5.98 | 1.29 | 0.68 |
| Viperinae-Candoiinae | -0.13 | -3.77 | 3.50 | 1.00 |
| Colubrinae-Charininae | 0.08 | -0.71 | 0.88 | 1.00 |
| Crotalinae-Charininae | -0.41 | -1.24 | 0.41 | 0.94 |
| Dipsadinae-Charininae | 0.49 | -0.38 | 1.35 | 0.86 |
| Elapinae-Charininae | -1.60 | -2.67 | -0.53 | <0.01 |
| Hydrophiinae-Charininae | -0.06 | -1.60 | 1.47 | 1.00 |
| Leptotyphlopinae-Charininae | 2.64 | 1.20 | 4.07 | <0.01 |
| Loxoceminae-Charininae | -0.87 | -3.93 | 2.20 | 1.00 |
| Natricinae-Charininae | -0.19 | -1.02 | 0.64 | 1.00 |
| Pseudaspidinae-Charininae | 0.17 | -2.89 | 3.24 | 1.00 |
| Pythoninae-Charininae | 1.17 | -1.90 | 4.24 | 1.00 |
| Tropidophiinae-Charininae | -1.13 | -3.37 | 1.10 | 0.93 |
| Viperinae-Charininae | 1.08 | -1.16 | 3.31 | 0.96 |
| Crotalinae-Colubrinae | -0.50 | -0.87 | -0.13 | <0.01 |
| Dipsadinae-Colubrinae | 0.40 | -0.05 | 0.86 | 0.15 |
| Elapinae-Colubrinae | -1.68 | -2.46 | -0.91 | <0.01 |
| Hydrophiinae-Colubrinae | -0.15 | -1.49 | 1.20 | 1.00 |
| Leptotyphlopinae-Colubrinae | 2.56 | 1.32 | 3.79 | <0.01 |
| Loxoceminae-Colubrinae | -0.95 | -3.93 | 2.03 | 1.00 |
| Natricinae-Colubrinae | -0.27 | -0.66 | 0.11 | 0.53 |
| Pseudaspidinae-Colubrinae | 0.09 | -2.89 | 3.07 | 1.00 |
| Pythoninae-Colubrinae | 1.09 | -1.89 | 4.06 | 1.00 |
| Tropidophiinae-Colubrinae | -1.22 | -3.33 | 0.89 | 0.83 |
| Viperinae-Colubrinae | 1.00 | -1.12 | 3.11 | 0.96 |
| Dipsadinae-Crotalinae | 0.90 | 0.40 | 1.40 | <0.01 |

**Table 11. (continued)**

| | | | | |
|---|---|---|---|---|
| Elapinae-Crotalinae | -1.19 | -1.99 | -0.39 | <0.01 |
| Hydrophiinae-Crotalinae | 0.35 | -1.01 | 1.71 | 1.00 |
| Leptotyphlopinae-Crotalinae | 3.05 | 1.80 | 4.30 | <0.01 |
| Loxoceminae-Crotalinae | -0.45 | -3.44 | 2.53 | 1.00 |
| Natricinae-Crotalinae | 0.23 | -0.21 | 0.67 | 0.92 |
| Pseudaspidinae-Crotalinae | 0.59 | -2.40 | 3.57 | 1.00 |
| Pythoninae-Crotalinae | 1.58 | -1.40 | 4.57 | 0.91 |
| Tropidophiinae-Crotalinae | -0.72 | -2.84 | 1.40 | 1.00 |
| Viperinae-Crotalinae | 1.49 | -0.63 | 3.61 | 0.53 |
| Elapinae-Dipsadinae | -2.09 | -2.93 | -1.24 | <0.01 |
| Hydrophiinae-Dipsadinae | -0.55 | -1.94 | 0.84 | 0.99 |
| Leptotyphlopinae-Dipsadinae | 2.15 | 0.87 | 3.43 | <0.01 |
| Loxoceminae-Dipsadinae | -1.35 | -4.35 | 1.64 | 0.97 |
| Natricinae-Dipsadinae | -0.67 | -1.19 | -0.16 | <0.01 |
| Pseudaspidinae-Dipsadinae | -0.31 | -3.31 | 2.68 | 1.00 |
| Pythoninae-Dipsadinae | 0.68 | -2.31 | 3.68 | 1.00 |
| Tropidophiinae-Dipsadinae | -1.62 | -3.76 | 0.52 | 0.39 |
| Viperinae-Dipsadinae | 0.59 | -1.55 | 2.73 | 1.00 |
| Hydrophiinae-Elapinae | 1.54 | 0.02 | 3.06 | 0.04 |
| Leptotyphlopinae-Elapinae | 4.24 | 2.82 | 5.66 | <0.01 |
| Loxoceminae-Elapinae | 0.73 | -2.33 | 3.79 | 1.00 |
| Natricinae-Elapinae | 1.41 | 0.61 | 2.22 | <0.01 |
| Pseudaspidinae-Elapinae | 1.77 | -1.29 | 4.83 | 0.83 |
| Pythoninae-Elapinae | 2.77 | -0.29 | 5.83 | 0.13 |
| Tropidophiinae-Elapinae | 0.47 | -1.76 | 2.69 | 1.00 |
| Viperinae-Elapinae | 2.68 | 0.45 | 4.91 | <0.01 |
| Leptotyphlopinae-Hydrophiinae | 2.70 | 0.90 | 4.50 | <0.01 |
| Loxoceminae-Hydrophiinae | -0.80 | -4.06 | 2.45 | 1.00 |
| Natricinae-Hydrophiinae | -0.12 | -1.49 | 1.24 | 1.00 |
| Pseudaspidinae-Hydrophiinae | 0.24 | -3.02 | 3.49 | 1.00 |
| Pythoninae-Hydrophiinae | 1.23 | -2.02 | 4.48 | 1.00 |
| Tropidophiinae-Hydrophiinae | -1.07 | -3.55 | 1.41 | 0.98 |
| Viperinae-Hydrophiinae | 1.14 | -1.34 | 3.63 | 0.97 |
| Loxoceminae-Leptotyphlopinae | -3.50 | -6.71 | -0.30 | 0.02 |
| Natricinae-Leptotyphlopinae | -2.83 | -4.08 | -1.57 | <0.01 |

**Table 11. (continued)**

| | | | | |
|---|---|---|---|---|
| Pseudaspidinae-Leptotyphlopinae | -2.47 | -5.67 | 0.74 | 0.37 |
| Pythoninae-Leptotyphlopinae | -1.47 | -4.68 | 1.74 | 0.97 |
| Tropidophiinae-Leptotyphlopinae | -3.77 | -6.20 | -1.35 | <0.01 |
| Viperinae-Leptotyphlopinae | -1.56 | -3.98 | 0.86 | 0.69 |
| Natricinae-Loxoceminae | 0.68 | -2.31 | 3.67 | 1.00 |
| Pseudaspidinae-Loxoceminae | 1.04 | -3.16 | 5.24 | 1.00 |
| Pythoninae-Loxoceminae | 2.04 | -2.16 | 6.23 | 0.95 |
| Tropidophiinae-Loxoceminae | -0.27 | -3.90 | 3.37 | 1.00 |
| Viperinae-Loxoceminae | 1.95 | -1.69 | 5.58 | 0.90 |
| Pseudaspidinae-Natricinae | 0.36 | -2.63 | 3.35 | 1.00 |
| Pythoninae-Natricinae | 1.36 | -1.63 | 4.34 | 0.97 |
| Tropidophiinae-Natricinae | -0.95 | -3.07 | 1.18 | 0.98 |
| Viperinae-Natricinae | 1.27 | -0.86 | 3.39 | 0.79 |
| Pythoninae-Pseudaspidinae | 1.00 | -3.20 | 5.19 | 1.00 |
| Tropidophiinae-Pseudaspidinae | -1.31 | -4.94 | 2.33 | 1.00 |
| Viperinae-Pseudaspidinae | 0.91 | -2.73 | 4.54 | 1.00 |
| Tropidophiinae-Pythoninae | -2.30 | -5.94 | 1.33 | 0.71 |
| Viperinae-Pythoninae | -0.09 | -3.73 | 3.55 | 1.00 |
| Viperinae-Tropidophiinae | 2.21 | -0.76 | 5.18 | 0.42 |
| PC5 All | | | | |
| Boinae-Ahaetuliinae | -0.82 | -2.12 | 0.47 | 0.71 |
| Candoiinae-Ahaetuliinae | -0.42 | -3.28 | 2.43 | 1.00 |
| Charininae-Ahaetuliinae | -0.03 | -1.38 | 1.31 | 1.00 |
| Colubrinae-Ahaetuliinae | 0.96 | -0.22 | 2.14 | 0.27 |
| Crotalinae-Ahaetuliinae | 1.48 | 0.28 | 2.67 | <0.01 |
| Dipsadinae-Ahaetuliinae | 0.94 | -0.27 | 2.16 | 0.35 |
| Elapinae-Ahaetuliinae | 1.12 | -0.22 | 2.45 | 0.23 |
| Hydrophiinae-Ahaetuliinae | 1.03 | -0.62 | 2.68 | 0.73 |
| Leptotyphlopinae-Ahaetuliinae | 1.50 | -0.08 | 3.08 | 0.08 |
| Loxoceminae-Ahaetuliinae | -0.58 | -3.43 | 2.28 | 1.00 |
| Natricinae-Ahaetuliinae | 2.11 | 0.92 | 3.31 | <0.01 |
| Pseudaspidinae-Ahaetuliinae | 1.11 | -1.74 | 3.97 | 0.99 |
| Pythoninae-Ahaetuliinae | -1.36 | -4.22 | 1.50 | 0.96 |
| Tropidophiinae-Ahaetuliinae | 0.75 | -1.43 | 2.93 | 1.00 |

**Table 11. (continued)**

| | | | | |
|---|---|---|---|---|
| Viperinae-Ahaetuliinae | 0.71 | -1.48 | 2.89 | 1.00 |
| Candoiinae-Boinae | 0.40 | -2.27 | 3.07 | 1.00 |
| Charininae-Boinae | 0.79 | -0.09 | 1.67 | 0.14 |
| Colubrinae-Boinae | 1.79 | 1.19 | 2.39 | <0.01 |
| Crotalinae-Boinae | 2.30 | 1.67 | 2.93 | <0.01 |
| Dipsadinae-Boinae | 1.77 | 1.10 | 2.44 | <0.01 |
| Elapinae-Boinae | 1.94 | 1.07 | 2.80 | <0.01 |
| Hydrophiinae-Boinae | 1.86 | 0.56 | 3.16 | <0.01 |
| Leptotyphlopinae-Boinae | 2.33 | 1.12 | 3.53 | <0.01 |
| Loxoceminae-Boinae | 0.25 | -2.42 | 2.92 | 1.00 |
| Natricinae-Boinae | 2.94 | 2.30 | 3.57 | <0.01 |
| Pseudaspidinae-Boinae | 1.94 | -0.73 | 4.61 | 0.47 |
| Pythoninae-Boinae | -0.53 | -3.20 | 2.14 | 1.00 |
| Tropidophiinae-Boinae | 1.57 | -0.36 | 3.50 | 0.27 |
| Viperinae-Boinae | 1.53 | -0.40 | 3.46 | 0.31 |
| Charininae-Candoiinae | 0.39 | -2.30 | 3.08 | 1.00 |
| Colubrinae-Candoiinae | 1.38 | -1.23 | 4.00 | 0.91 |
| Crotalinae-Candoiinae | 1.90 | -0.72 | 4.52 | 0.48 |
| Dipsadinae-Candoiinae | 1.37 | -1.26 | 4.00 | 0.92 |
| Elapinae-Candoiinae | 1.54 | -1.15 | 4.23 | 0.84 |
| Hydrophiinae-Candoiinae | 1.46 | -1.40 | 4.31 | 0.93 |
| Leptotyphlopinae-Candoiinae | 1.93 | -0.89 | 4.74 | 0.58 |
| Loxoceminae-Candoiinae | -0.15 | -3.84 | 3.54 | 1.00 |
| Natricinae-Candoiinae | 2.54 | -0.08 | 5.16 | 0.07 |
| Pseudaspidinae-Candoiinae | 1.54 | -2.15 | 5.23 | 0.99 |
| Pythoninae-Candoiinae | -0.93 | -4.62 | 2.75 | 1.00 |
| Tropidophiinae-Candoiinae | 1.17 | -2.02 | 4.37 | 1.00 |
| Viperinae-Candoiinae | 1.13 | -2.06 | 4.32 | 1.00 |
| Colubrinae-Charininae | 0.99 | 0.30 | 1.69 | <0.01 |
| Crotalinae-Charininae | 1.51 | 0.79 | 2.23 | <0.01 |
| Dipsadinae-Charininae | 0.98 | 0.22 | 1.74 | <0.01 |
| Elapinae-Charininae | 1.15 | 0.21 | 2.09 | <0.01 |
| Hydrophiinae-Charininae | 1.07 | -0.28 | 2.41 | 0.31 |
| Leptotyphlopinae-Charininae | 1.54 | 0.28 | 2.80 | <0.01 |
| Loxoceminae-Charininae | -0.54 | -3.24 | 2.15 | 1.00 |
| Natricinae-Charininae | 2.15 | 1.42 | 2.88 | <0.01 |
| Pseudaspidinae-Charininae | 1.15 | -1.55 | 3.84 | 0.99 |

**Table 11. (continued)**

| | | | | |
|---|---|---|---|---|
| Pythoninae-Charininae | -1.32 | -4.02 | 1.37 | 0.95 |
| Tropidophiinae-Charininae | 0.78 | -1.18 | 2.75 | 0.99 |
| Viperinae-Charininae | 0.74 | -1.22 | 2.70 | 1.00 |
| Crotalinae-Colubrinae | 0.52 | 0.19 | 0.84 | <0.01 |
| Dipsadinae-Colubrinae | -0.02 | -0.42 | 0.38 | 1.00 |
| Elapinae-Colubrinae | 0.15 | -0.52 | 0.83 | 1.00 |
| Hydrophiinae-Colubrinae | 0.07 | -1.11 | 1.25 | 1.00 |
| Leptotyphlopinae-Colubrinae | 0.54 | -0.54 | 1.62 | 0.94 |
| Loxoceminae-Colubrinae | -1.54 | -4.15 | 1.08 | 0.81 |
| Natricinae-Colubrinae | 1.15 | 0.82 | 1.49 | <0.01 |
| Pseudaspidinae-Colubrinae | 0.15 | -2.46 | 2.77 | 1.00 |
| Pythoninae-Colubrinae | -2.32 | -4.93 | 0.30 | 0.15 |
| Tropidophiinae-Colubrinae | -0.21 | -2.07 | 1.64 | 1.00 |
| Viperinae-Colubrinae | -0.26 | -2.11 | 1.60 | 1.00 |
| Dipsadinae-Crotalinae | -0.53 | -0.97 | -0.09 | <0.01 |
| Elapinae-Crotalinae | -0.36 | -1.07 | 0.34 | 0.93 |
| Hydrophiinae-Crotalinae | -0.44 | -1.64 | 0.75 | 1.00 |
| Leptotyphlopinae-Crotalinae | 0.03 | -1.07 | 1.12 | 1.00 |
| Loxoceminae-Crotalinae | -2.05 | -4.68 | 0.57 | 0.33 |
| Natricinae-Crotalinae | 0.64 | 0.25 | 1.02 | <0.01 |
| Pseudaspidinae-Crotalinae | -0.36 | -2.99 | 2.26 | 1.00 |
| Pythoninae-Crotalinae | -2.83 | -5.46 | -0.21 | 0.02 |
| Tropidophiinae-Crotalinae | -0.73 | -2.59 | 1.13 | 0.99 |
| Viperinae-Crotalinae | -0.77 | -2.63 | 1.09 | 0.99 |
| Elapinae-Dipsadinae | 0.17 | -0.57 | 0.91 | 1.00 |
| Hydrophiinae-Dipsadinae | 0.09 | -1.13 | 1.31 | 1.00 |
| Leptotyphlopinae-Dipsadinae | 0.56 | -0.56 | 1.68 | 0.94 |
| Loxoceminae-Dipsadinae | -1.52 | -4.15 | 1.11 | 0.83 |
| Natricinae-Dipsadinae | 1.17 | 0.72 | 1.62 | <0.01 |
| Pseudaspidinae-Dipsadinae | 0.17 | -2.46 | 2.80 | 1.00 |
| Pythoninae-Dipsadinae | -2.30 | -4.93 | 0.33 | 0.17 |
| Tropidophiinae-Dipsadinae | -0.20 | -2.07 | 1.68 | 1.00 |
| Viperinae-Dipsadinae | -0.24 | -2.12 | 1.64 | 1.00 |
| Hydrophiinae-Elapinae | -0.08 | -1.42 | 1.26 | 1.00 |
| Leptotyphlopinae-Elapinae | 0.39 | -0.86 | 1.64 | 1.00 |
| Loxoceminae-Elapinae | -1.69 | -4.38 | 1.00 | 0.72 |
| Natricinae-Elapinae | 1.00 | 0.29 | 1.71 | <0.01 |

**Table 11. (continued)**

| | | | | |
|---|---|---|---|---|
| Pseudaspidinae-Elapinae | <0.01 | -2.69 | 2.69 | 1.00 |
| Pythoninae-Elapinae | -2.47 | -5.16 | 0.21 | 0.11 |
| Tropidophiinae-Elapinae | -0.37 | -2.32 | 1.59 | 1.00 |
| Viperinae-Elapinae | -0.41 | -2.37 | 1.55 | 1.00 |
| Leptotyphlopinae-Hydrophiinae | 0.47 | -1.11 | 2.05 | 1.00 |
| Loxoceminae-Hydrophiinae | -1.61 | -4.47 | 1.25 | 0.85 |
| Natricinae-Hydrophiinae | 1.08 | -0.12 | 2.28 | 0.13 |
| Pseudaspidinae-Hydrophiinae | 0.08 | -2.78 | 2.94 | 1.00 |
| Pythoninae-Hydrophiinae | -2.39 | -5.25 | 0.46 | 0.23 |
| Tropidophiinae-Hydrophiinae | -0.29 | -2.47 | 1.90 | 1.00 |
| Viperinae-Hydrophiinae | -0.33 | -2.51 | 1.85 | 1.00 |
| Loxoceminae-Leptotyphlopinae | -2.08 | -4.90 | 0.74 | 0.44 |
| Natricinae-Leptotyphlopinae | 0.61 | -0.49 | 1.71 | 0.87 |
| Pseudaspidinae-Leptotyphlopinae | -0.39 | -3.21 | 2.43 | 1.00 |
| Pythoninae-Leptotyphlopinae | -2.86 | -5.68 | -0.04 | 0.04 |
| Tropidophiinae-Leptotyphlopinae | -0.76 | -2.88 | 1.37 | 1.00 |
| Viperinae-Leptotyphlopinae | -0.80 | -2.93 | 1.33 | 1.00 |
| Natricinae-Loxoceminae | 2.69 | 0.07 | 5.32 | 0.04 |
| Pseudaspidinae-Loxoceminae | 1.69 | -2.00 | 5.38 | 0.97 |
| Pythoninae-Loxoceminae | -0.78 | -4.47 | 2.91 | 1.00 |
| Tropidophiinae-Loxoceminae | 1.33 | -1.87 | 4.52 | 0.99 |
| Viperinae-Loxoceminae | 1.28 | -1.91 | 4.48 | 0.99 |
| Pseudaspidinae-Natricinae | -1.00 | -3.63 | 1.62 | 1.00 |
| Pythoninae-Natricinae | -3.47 | -6.10 | -0.85 | <0.01 |
| Tropidophiinae-Natricinae | -1.37 | -3.23 | 0.50 | 0.46 |
| Viperinae-Natricinae | -1.41 | -3.27 | 0.46 | 0.40 |
| Pythoninae-Pseudaspidinae | -2.47 | -6.16 | 1.22 | 0.62 |
| Tropidophiinae-Pseudaspidinae | -0.36 | -3.56 | 2.83 | 1.00 |
| Viperinae-Pseudaspidinae | -0.41 | -3.60 | 2.79 | 1.00 |
| Tropidophiinae-Pythoninae | 2.11 | -1.09 | 5.30 | 0.65 |
| Viperinae-Pythoninae | 2.06 | -1.13 | 5.26 | 0.68 |
| Viperinae-Tropidophiinae | -0.04 | -2.65 | 2.57 | 1.00 |

**Table 11. (continued)**

| | | | | |
|---|---|---|---|---|
| PC6 All | | | | |
| Boinae-Ahaetuliinae | 1.63 | 0.16 | 3.11 | 0.01 |
| Candoiinae-Ahaetuliinae | 1.01 | -2.24 | 4.26 | 1.00 |
| Charininae-Ahaetuliinae | 1.60 | 0.07 | 3.13 | 0.03 |
| Colubrinae-Ahaetuliinae | 1.00 | -0.34 | 2.35 | 0.42 |
| Crotalinae-Ahaetuliinae | -0.04 | -1.40 | 1.32 | 1.00 |
| Dipsadinae-Ahaetuliinae | 0.33 | -1.06 | 1.71 | 1.00 |
| Elapinae-Ahaetuliinae | -0.21 | -1.73 | 1.31 | 1.00 |
| Hydrophiinae-Ahaetuliinae | 0.09 | -1.79 | 1.97 | 1.00 |
| Leptotyphlopinae-Ahaetuliinae | -0.79 | -2.59 | 1.01 | 0.98 |
| Loxoceminae-Ahaetuliinae | 1.72 | -1.53 | 4.97 | 0.91 |
| Natricinae-Ahaetuliinae | 0.56 | -0.80 | 1.93 | 0.99 |
| Pseudaspidinae-Ahaetuliinae | 1.35 | -1.90 | 4.60 | 0.99 |
| Pythoninae-Ahaetuliinae | 1.52 | -1.73 | 4.77 | 0.97 |
| Tropidophiinae-Ahaetuliinae | 0.52 | -1.96 | 3.01 | 1.00 |
| Viperinae-Ahaetuliinae | -0.01 | -2.50 | 2.47 | 1.00 |
| Candoiinae-Boinae | -0.62 | -3.66 | 2.42 | 1.00 |
| Charininae-Boinae | -0.03 | -1.04 | 0.97 | 1.00 |
| Colubrinae-Boinae | -0.63 | -1.31 | 0.05 | 0.11 |
| Crotalinae-Boinae | -1.67 | -2.39 | -0.96 | <0.01 |
| Dipsadinae-Boinae | -1.31 | -2.07 | -0.55 | <0.01 |
| Elapinae-Boinae | -1.84 | -2.83 | -0.86 | <0.01 |
| Hydrophiinae-Boinae | -1.54 | -3.02 | -0.07 | 0.03 |
| Leptotyphlopinae-Boinae | -2.42 | -3.80 | -1.05 | <0.01 |
| Loxoceminae-Boinae | 0.09 | -2.95 | 3.12 | 1.00 |
| Natricinae-Boinae | -1.07 | -1.79 | -0.35 | <0.01 |
| Pseudaspidinae-Boinae | -0.29 | -3.32 | 2.75 | 1.00 |
| Pythoninae-Boinae | -0.11 | -3.15 | 2.92 | 1.00 |
| Tropidophiinae-Boinae | -1.11 | -3.31 | 1.09 | 0.93 |
| Viperinae-Boinae | -1.65 | -3.84 | 0.55 | 0.41 |
| Charininae-Candoiinae | 0.59 | -2.48 | 3.65 | 1.00 |
| Colubrinae-Candoiinae | -0.01 | -2.99 | 2.96 | 1.00 |
| Crotalinae-Candoiinae | -1.05 | -4.04 | 1.93 | 1.00 |
| Dipsadinae-Candoiinae | -0.69 | -3.68 | 2.31 | 1.00 |
| Elapinae-Candoiinae | -1.22 | -4.28 | 1.84 | 0.99 |
| Hydrophiinae-Candoiinae | -0.92 | -4.17 | 2.33 | 1.00 |

**Table 11. (continued)**

| | | | | |
|---|---|---|---|---|
| Leptotyphlopinae-Candoiinae | -1.80 | -5.01 | 1.40 | 0.86 |
| Loxoceminae-Candoiinae | 0.71 | -3.49 | 4.90 | 1.00 |
| Natricinae-Candoiinae | -0.45 | -3.44 | 2.53 | 1.00 |
| Pseudaspidinae-Candoiinae | 0.33 | -3.86 | 4.53 | 1.00 |
| Pythoninae-Candoiinae | 0.51 | -3.69 | 4.70 | 1.00 |
| Tropidophiinae-Candoiinae | -0.49 | -4.12 | 3.14 | 1.00 |
| Viperinae-Candoiinae | -1.03 | -4.66 | 2.61 | 1.00 |
| Colubrinae-Charininae | -0.60 | -1.39 | 0.20 | 0.41 |
| Crotalinae-Charininae | -1.64 | -2.46 | -0.82 | <0.01 |
| Dipsadinae-Charininae | -1.28 | -2.14 | -0.41 | <0.01 |
| Elapinae-Charininae | -1.81 | -2.87 | -0.74 | <0.01 |
| Hydrophiinae-Charininae | -1.51 | -3.04 | 0.02 | 0.06 |
| Leptotyphlopinae-Charininae | -2.39 | -3.83 | -0.96 | <0.01 |
| Loxoceminae-Charininae | 0.12 | -2.95 | 3.18 | 1.00 |
| Natricinae-Charininae | -1.04 | -1.87 | -0.21 | <0.01 |
| Pseudaspidinae-Charininae | -0.25 | -3.32 | 2.81 | 1.00 |
| Pythoninae-Charininae | -0.08 | -3.14 | 2.98 | 1.00 |
| Tropidophiinae-Charininae | -1.08 | -3.31 | 1.16 | 0.96 |
| Viperinae-Charininae | -1.62 | -3.85 | 0.62 | 0.48 |
| Crotalinae-Colubrinae | -1.04 | -1.41 | -0.67 | <0.01 |
| Dipsadinae-Colubrinae | -0.68 | -1.13 | -0.22 | <0.01 |
| Elapinae-Colubrinae | -1.21 | -1.98 | -0.44 | <0.01 |
| Hydrophiinae-Colubrinae | -0.91 | -2.26 | 0.43 | 0.59 |
| Leptotyphlopinae-Colubrinae | -1.79 | -3.02 | -0.56 | <0.01 |
| Loxoceminae-Colubrinae | 0.72 | -2.26 | 3.69 | 1.00 |
| Natricinae-Colubrinae | -0.44 | -0.83 | -0.06 | 0.01 |
| Pseudaspidinae-Colubrinae | 0.34 | -2.63 | 3.32 | 1.00 |
| Pythoninae-Colubrinae | 0.52 | -2.46 | 3.49 | 1.00 |
| Tropidophiinae-Colubrinae | -0.48 | -2.59 | 1.63 | 1.00 |
| Viperinae-Colubrinae | -1.02 | -3.13 | 1.09 | 0.96 |
| Dipsadinae-Crotalinae | 0.36 | -0.14 | 0.86 | 0.47 |
| Elapinae-Crotalinae | -0.17 | -0.97 | 0.63 | 1.00 |
| Hydrophiinae-Crotalinae | 0.13 | -1.23 | 1.49 | 1.00 |
| Leptotyphlopinae-Crotalinae | -0.75 | -2.00 | 0.50 | 0.78 |
| Loxoceminae-Crotalinae | 1.76 | -1.22 | 4.74 | 0.81 |
| Natricinae-Crotalinae | 0.60 | 0.16 | 1.04 | <0.01 |
| Pseudaspidinae-Crotalinae | 1.39 | -1.60 | 4.37 | 0.97 |

**Table 11. (continued)**

| | | | | |
|---|---|---|---|---|
| Pythoninae-Crotalinae | 1.56 | -1.42 | 4.54 | 0.91 |
| Tropidophiinae-Crotalinae | 0.56 | -1.56 | 2.68 | 1.00 |
| Viperinae-Crotalinae | 0.02 | -2.10 | 2.14 | 1.00 |
| Elapinae-Dipsadinae | -0.53 | -1.37 | 0.31 | 0.72 |
| Hydrophiinae-Dipsadinae | -0.24 | -1.62 | 1.15 | 1.00 |
| Leptotyphlopinae-Dipsadinae | -1.12 | -2.39 | 0.16 | 0.17 |
| Loxoceminae-Dipsadinae | 1.40 | -1.60 | 4.39 | 0.97 |
| Natricinae-Dipsadinae | 0.24 | -0.27 | 0.75 | 0.97 |
| Pseudaspidinae-Dipsadinae | 1.02 | -1.97 | 4.02 | 1.00 |
| Pythoninae-Dipsadinae | 1.20 | -1.80 | 4.19 | 0.99 |
| Tropidophiinae-Dipsadinae | 0.20 | -1.94 | 2.34 | 1.00 |
| Viperinae-Dipsadinae | -0.34 | -2.48 | 1.80 | 1.00 |
| Hydrophiinae-Elapinae | 0.30 | -1.22 | 1.82 | 1.00 |
| Leptotyphlopinae-Elapinae | -0.58 | -2.00 | 0.84 | 0.99 |
| Loxoceminae-Elapinae | 1.93 | -1.13 | 4.99 | 0.72 |
| Natricinae-Elapinae | 0.77 | -0.04 | 1.58 | 0.08 |
| Pseudaspidinae-Elapinae | 1.55 | -1.50 | 4.61 | 0.93 |
| Pythoninae-Elapinae | 1.73 | -1.33 | 4.79 | 0.85 |
| Tropidophiinae-Elapinae | 0.73 | -1.49 | 2.96 | 1.00 |
| Viperinae-Elapinae | 0.19 | -2.03 | 2.42 | 1.00 |
| Leptotyphlopinae-Hydrophiinae | -0.88 | -2.68 | 0.92 | 0.95 |
| Loxoceminae-Hydrophiinae | 1.63 | -1.62 | 4.88 | 0.94 |
| Natricinae-Hydrophiinae | 0.47 | -0.89 | 1.84 | 1.00 |
| Pseudaspidinae-Hydrophiinae | 1.26 | -1.99 | 4.51 | 0.99 |
| Pythoninae-Hydrophiinae | 1.43 | -1.82 | 4.68 | 0.98 |
| Tropidophiinae-Hydrophiinae | 0.43 | -2.05 | 2.92 | 1.00 |
| Viperinae-Hydrophiinae | -0.10 | -2.59 | 2.38 | 1.00 |
| Loxoceminae-Leptotyphlopinae | 2.51 | -0.69 | 5.72 | 0.34 |
| Natricinae-Leptotyphlopinae | 1.35 | 0.10 | 2.60 | 0.02 |
| Pseudaspidinae-Leptotyphlopinae | 2.14 | -1.07 | 5.34 | 0.63 |
| Pythoninae-Leptotyphlopinae | 2.31 | -0.89 | 5.52 | 0.48 |
| Tropidophiinae-Leptotyphlopinae | 1.31 | -1.11 | 3.74 | 0.89 |
| Viperinae-Leptotyphlopinae | 0.78 | -1.65 | 3.20 | 1.00 |
| Natricinae-Loxoceminae | -1.16 | -4.14 | 1.83 | 0.99 |

**Table 11. (continued)**

| | | | | |
|---|---|---|---|---|
| Pseudaspidinae-Loxoceminae | -0.37 | -4.57 | 3.82 | 1.00 |
| Pythoninae-Loxoceminae | -0.20 | -4.39 | 4.00 | 1.00 |
| Tropidophiinae-Loxoceminae | -1.20 | -4.83 | 2.44 | 1.00 |
| Viperinae-Loxoceminae | -1.73 | -5.37 | 1.90 | 0.96 |
| Pseudaspidinae-Natricinae | 0.79 | -2.20 | 3.77 | 1.00 |
| Pythoninae-Natricinae | 0.96 | -2.02 | 3.95 | 1.00 |
| Tropidophiinae-Natricinae | -0.04 | -2.16 | 2.09 | 1.00 |
| Viperinae-Natricinae | -0.58 | -2.70 | 1.55 | 1.00 |
| Pythoninae-Pseudaspidinae | 0.18 | -4.02 | 4.37 | 1.00 |
| Tropidophiinae-Pseudaspidinae | -0.82 | -4.46 | 2.81 | 1.00 |
| Viperinae-Pseudaspidinae | -1.36 | -5.00 | 2.27 | 1.00 |
| Tropidophiinae-Pythoninae | -1.00 | -4.63 | 2.64 | 1.00 |
| Viperinae-Pythoninae | -1.54 | -5.17 | 2.10 | 0.99 |
| Viperinae-Tropidophiinae | -0.54 | -3.51 | 2.43 | 1.00 |

**Table 12. Tukey's test results of primary foraging habitat for PCs 1-6 of the all-groups data.**

| PC1 All | diff | lwr | upr | p adj |
|---|---|---|---|---|
| Arboreal-Aquatic | 0.71 | 0.15 | 1.27 | <0.01 |
| Fossorial-Aquatic | 1.58 | 1.03 | 2.14 | <0.01 |
| Semiaquatic-Aquatic | 0.03 | -0.55 | 0.62 | 1.00 |
| Semiarboreal-Aquatic | 0.63 | 0.06 | 1.21 | 0.02 |
| Semifossorial-Aquatic | 1.04 | 0.52 | 1.56 | <0.01 |
| Terrestrial-Aquatic | 0.13 | -0.36 | 0.61 | 0.99 |
| Fossorial-Arboreal | 0.87 | 0.39 | 1.35 | <0.01 |
| Semiaquatic-Arboreal | -0.68 | -1.20 | -0.16 | <0.01 |
| Semiarboreal-Arboreal | -0.08 | -0.58 | 0.42 | 1.00 |
| Semifossorial-Arboreal | 0.33 | -0.11 | 0.76 | 0.28 |
| Terrestrial-Arboreal | -0.59 | -0.98 | -0.20 | <0.01 |
| Semiaquatic-Fossorial | -1.55 | -2.06 | -1.03 | <0.01 |
| Semiarboreal-Fossorial | -0.95 | -1.45 | -0.45 | <0.01 |
| Semifossorial-Fossorial | -0.54 | -0.97 | -0.11 | <0.01 |
| Terrestrial-Fossorial | -1.46 | -1.84 | -1.07 | <0.01 |
| Semiarboreal-Semiaquatic | 0.60 | 0.07 | 1.13 | 0.02 |
| Semifossorial-Semiaquatic | 1.01 | 0.53 | 1.48 | <0.01 |
| Terrestrial-Semiaquatic | 0.09 | -0.34 | 0.52 | 1.00 |
| Semifossorial-Semiarboreal | 0.41 | -0.05 | 0.86 | 0.11 |
| Terrestrial-Semiarboreal | -0.51 | -0.92 | -0.09 | 0.01 |
| Terrestrial-Semifossorial | -0.91 | -1.24 | -0.58 | <0.01 |
| PC2 All | | | | |
| Arboreal-Aquatic | 0.23 | -0.36 | 0.82 | 0.91 |
| Fossorial-Aquatic | 0.92 | 0.34 | 1.51 | <0.01 |
| Semiaquatic-Aquatic | 0.61 | -0.01 | 1.22 | 0.06 |
| Semiarboreal-Aquatic | -0.03 | -0.63 | 0.58 | 1.00 |
| Semifossorial-Aquatic | 1.27 | 0.72 | 1.81 | <0.01 |
| Terrestrial-Aquatic | 0.98 | 0.47 | 1.49 | <0.01 |
| Fossorial-Arboreal | 0.69 | 0.19 | 1.20 | <0.01 |
| Semiaquatic-Arboreal | 0.38 | -0.17 | 0.92 | 0.39 |
| Semiarboreal-Arboreal | -0.26 | -0.78 | 0.27 | 0.78 |
| Semifossorial-Arboreal | 1.04 | 0.58 | 1.49 | <0.01 |
| Terrestrial-Arboreal | 0.75 | 0.34 | 1.16 | <0.01 |
| Semiaquatic-Fossorial | -0.32 | -0.86 | 0.22 | 0.59 |
| Semiarboreal-Fossorial | -0.95 | -1.47 | -0.42 | <0.01 |
| Semifossorial-Fossorial | 0.34 | -0.11 | 0.80 | 0.27 |
| Terrestrial-Fossorial | 0.06 | -0.35 | 0.46 | 1.00 |

**Table 12. (continued)**

| | | | | |
|---|---|---|---|---|
| Semiarboreal-Semiaquatic | -0.63 | -1.19 | -0.07 | 0.02 |
| Semifossorial-Semiaquatic | 0.66 | 0.17 | 1.16 | <0.01 |
| Terrestrial-Semiaquatic | 0.37 | -0.08 | 0.83 | 0.19 |
| Semifossorial-Semiarboreal | 1.29 | 0.82 | 1.77 | <0.01 |
| Terrestrial-Semiarboreal | 1.01 | 0.57 | 1.44 | <0.01 |
| Terrestrial-Semifossorial | -0.29 | -0.63 | 0.06 | 0.18 |
| PC3 All | | | | |
| Arboreal-Aquatic | 0.84 | 0.22 | 1.47 | <0.01 |
| Fossorial-Aquatic | 0.54 | -0.08 | 1.16 | 0.14 |
| Semiaquatic-Aquatic | 0.02 | -0.64 | 0.68 | 1.00 |
| Semiarboreal-Aquatic | 0.07 | -0.57 | 0.71 | 1.00 |
| Semifossorial-Aquatic | -0.17 | -0.75 | 0.41 | 0.98 |
| Terrestrial-Aquatic | 0.20 | -0.34 | 0.74 | 0.93 |
| Fossorial-Arboreal | -0.30 | -0.84 | 0.23 | 0.63 |
| Semiaquatic-Arboreal | -0.83 | -1.41 | -0.25 | <0.01 |
| Semiarboreal-Arboreal | -0.78 | -1.34 | -0.22 | <0.01 |
| Semifossorial-Arboreal | -1.01 | -1.50 | -0.53 | <0.01 |
| Terrestrial-Arboreal | -0.64 | -1.08 | -0.21 | <0.01 |
| Semiaquatic-Fossorial | -0.52 | -1.10 | 0.05 | 0.10 |
| Semiarboreal-Fossorial | -0.47 | -1.03 | 0.08 | 0.16 |
| Semifossorial-Fossorial | -0.71 | -1.19 | -0.23 | <0.01 |
| Terrestrial-Fossorial | -0.34 | -0.77 | 0.09 | 0.23 |
| Semiarboreal-Semiaquatic | 0.05 | -0.55 | 0.65 | 1.00 |
| Semifossorial-Semiaquatic | -0.19 | -0.71 | 0.34 | 0.94 |
| Terrestrial-Semiaquatic | 0.18 | -0.30 | 0.67 | 0.92 |
| Semifossorial-Semiarboreal | -0.24 | -0.74 | 0.27 | 0.81 |
| Terrestrial-Semiarboreal | 0.13 | -0.33 | 0.60 | 0.98 |
| Terrestrial-Semifossorial | 0.37 | <0.01 | 0.74 | 0.05 |
| PC4 All | | | | |
| Arboreal-Aquatic | -0.23 | -0.87 | 0.42 | 0.94 |
| Fossorial-Aquatic | 0.03 | -0.61 | 0.67 | 1.00 |
| Semiaquatic-Aquatic | 0.05 | -0.63 | 0.73 | 1.00 |
| Semiarboreal-Aquatic | 0.21 | -0.45 | 0.88 | 0.96 |
| Semifossorial-Aquatic | 0.09 | -0.51 | 0.68 | 1.00 |
| Terrestrial-Aquatic | -0.27 | -0.82 | 0.29 | 0.79 |
| Fossorial-Arboreal | 0.26 | -0.29 | 0.81 | 0.81 |
| Semiaquatic-Arboreal | 0.28 | -0.32 | 0.87 | 0.82 |

**Table 12. (continued)**

| | | | | |
|---|---|---|---|---|
| Semiarboreal-Arboreal | 0.44 | -0.14 | 1.02 | 0.26 |
| Semifossorial-Arboreal | 0.32 | -0.18 | 0.82 | 0.50 |
| Terrestrial-Arboreal | -0.04 | -0.49 | 0.41 | 1.00 |
| Semiaquatic-Fossorial | 0.02 | -0.58 | 0.61 | 1.00 |
| Semiarboreal-Fossorial | 0.18 | -0.39 | 0.76 | 0.96 |
| Semifossorial-Fossorial | 0.06 | -0.44 | 0.55 | 1.00 |
| Terrestrial-Fossorial | -0.30 | -0.75 | 0.15 | 0.43 |
| Semiarboreal-Semiaquatic | 0.17 | -0.45 | 0.78 | 0.98 |
| Semifossorial-Semiaquatic | 0.04 | -0.50 | 0.58 | 1.00 |
| Terrestrial-Semiaquatic | -0.31 | -0.81 | 0.18 | 0.50 |
| Semifossorial-Semiarboreal | -0.13 | -0.65 | 0.40 | 0.99 |
| Terrestrial-Semiarboreal | -0.48 | -0.96 | -0.01 | 0.04 |
| Terrestrial-Semifossorial | -0.36 | -0.74 | 0.02 | 0.08 |
| PC5 All | | | | |
| Arboreal-Aquatic | -1.95 | -2.52 | -1.38 | <0.01 |
| Fossorial-Aquatic | -0.55 | -1.12 | 0.02 | 0.07 |
| Semiaquatic-Aquatic | -0.24 | -0.84 | 0.36 | 0.90 |
| Semiarboreal-Aquatic | -1.17 | -1.76 | -0.58 | <0.01 |
| Semifossorial-Aquatic | -0.82 | -1.35 | -0.29 | <0.01 |
| Terrestrial-Aquatic | -0.68 | -1.17 | -0.19 | <0.01 |
| Fossorial-Arboreal | 1.40 | 0.91 | 1.89 | <0.01 |
| Semiaquatic-Arboreal | 1.71 | 1.18 | 2.24 | <0.01 |
| Semiarboreal-Arboreal | 0.78 | 0.26 | 1.29 | <0.01 |
| Semifossorial-Arboreal | 1.13 | 0.68 | 1.57 | <0.01 |
| Terrestrial-Arboreal | 1.27 | 0.87 | 1.67 | <0.01 |
| Semiaquatic-Fossorial | 0.31 | -0.21 | 0.84 | 0.58 |
| Semiarboreal-Fossorial | -0.62 | -1.13 | -0.11 | 0.01 |
| Semifossorial-Fossorial | -0.27 | -0.71 | 0.17 | 0.54 |
| Terrestrial-Fossorial | -0.13 | -0.52 | 0.27 | 0.96 |
| Semiarboreal-Semiaquatic | -0.93 | -1.48 | -0.39 | <0.01 |
| Semifossorial-Semiaquatic | -0.58 | -1.06 | -0.10 | 0.01 |
| Terrestrial-Semiaquatic | -0.44 | -0.88 | <0.01 | 0.05 |
| Semifossorial-Semiarboreal | 0.35 | -0.11 | 0.82 | 0.28 |
| Terrestrial-Semiarboreal | 0.49 | 0.07 | 0.91 | 0.01 |
| Terrestrial-Semifossorial | 0.14 | -0.20 | 0.48 | 0.88 |

**Table 12. (continued)**

| | | | | |
|---|---|---|---|---|
| PC6 All | | | | |
| Arboreal-Aquatic | -0.38 | -0.96 | 0.20 | 0.47 |
| Fossorial-Aquatic | -0.03 | -0.61 | 0.55 | 1.00 |
| Semiaquatic-Aquatic | -0.49 | -1.09 | 0.12 | 0.22 |
| Semiarboreal-Aquatic | 0.66 | 0.06 | 1.25 | 0.02 |
| Semifossorial-Aquatic | 0.77 | 0.24 | 1.31 | <0.01 |
| Terrestrial-Aquatic | -0.32 | -0.82 | 0.18 | 0.48 |
| Fossorial-Arboreal | 0.35 | -0.15 | 0.84 | 0.38 |
| Semiaquatic-Arboreal | -0.11 | -0.64 | 0.43 | 1.00 |
| Semiarboreal-Arboreal | 1.03 | 0.51 | 1.55 | <0.01 |
| Semifossorial-Arboreal | 1.15 | 0.70 | 1.60 | <0.01 |
| Terrestrial-Arboreal | 0.06 | -0.35 | 0.46 | 1.00 |
| Semiaquatic-Fossorial | -0.45 | -0.99 | 0.08 | 0.15 |
| Semiarboreal-Fossorial | 0.69 | 0.17 | 1.20 | <0.01 |
| Semifossorial-Fossorial | 0.80 | 0.36 | 1.25 | <0.01 |
| Terrestrial-Fossorial | -0.29 | -0.69 | 0.11 | 0.33 |
| Semiarboreal-Semiaquatic | 1.14 | 0.59 | 1.69 | <0.01 |
| Semifossorial-Semiaquatic | 1.26 | 0.77 | 1.74 | <0.01 |
| Terrestrial-Semiaquatic | 0.17 | -0.28 | 0.61 | 0.93 |
| Semifossorial-Semiarboreal | 0.11 | -0.36 | 0.58 | 0.99 |
| Terrestrial-Semiarboreal | -0.98 | -1.40 | -0.55 | <0.01 |
| Terrestrial-Semifossorial | -1.09 | -1.43 | -0.75 | <0.01 |

**Table 13. Tukey's test results of genus-level taxonomy for PCs 1-6 of the Crotalinae-only data.**

| PC1 Crotalinae | diff | lwr | upr | p adj |
|---|---|---|---|---|
| Bothriechis-Agkistrodon | 1.35 | 0.35 | 2.35 | <0.01 |
| Bothrops-Agkistrodon | 0.30 | -2.12 | 2.72 | 1.00 |
| Crotalus-Agkistrodon | 0.24 | -0.40 | 0.87 | 0.84 |
| Sistrurus-Agkistrodon | -1.12 | -2.02 | -0.22 | 0.01 |
| Bothrops-Bothriechis | -1.05 | -3.55 | 1.45 | 0.77 |
| Crotalus-Bothriechis | -1.11 | -2.00 | -0.23 | 0.01 |
| Sistrurus-Bothriechis | -2.47 | -3.56 | -1.37 | <0.01 |
| Crotalus-Bothrops | -0.07 | -2.44 | 2.31 | 1.00 |
| Sistrurus-Bothrops | -1.42 | -3.88 | 1.04 | 0.50 |
| Sistrurus-Crotalus | -1.35 | -2.13 | -0.58 | <0.01 |
| PC2 Crotalinae | | | | |
| Bothriechis-Agkistrodon | 0.43 | -0.65 | 1.50 | 0.80 |
| Bothrops-Agkistrodon | -1.73 | -4.33 | 0.88 | 0.35 |
| Crotalus-Agkistrodon | 0.64 | -0.04 | 1.32 | 0.08 |
| Sistrurus-Agkistrodon | -0.51 | -1.48 | 0.46 | 0.60 |
| Bothrops-Bothriechis | -2.15 | -4.84 | 0.54 | 0.18 |
| Crotalus-Bothriechis | 0.22 | -0.74 | 1.17 | 0.97 |
| Sistrurus-Bothriechis | -0.93 | -2.11 | 0.24 | 0.19 |
| Crotalus-Bothrops | 2.37 | -0.19 | 4.92 | 0.08 |
| Sistrurus-Bothrops | 1.22 | -1.43 | 3.87 | 0.70 |
| Sistrurus-Crotalus | -1.15 | -1.98 | -0.32 | <0.01 |
| PC3 Crotalinae | | | | |
| Bothriechis-Agkistrodon | 1.42 | 0.36 | 2.49 | <0.01 |
| Bothrops-Agkistrodon | -0.62 | -3.20 | 1.96 | 0.96 |
| Crotalus-Agkistrodon | 0.84 | 0.16 | 1.52 | 0.01 |
| Sistrurus-Agkistrodon | 1.43 | 0.46 | 2.39 | <0.01 |
| Bothrops-Bothriechis | -2.04 | -4.70 | 0.63 | 0.22 |
| Crotalus-Bothriechis | -0.58 | -1.53 | 0.37 | 0.43 |
| Sistrurus-Bothriechis | <0.01 | -1.16 | 1.17 | 1.00 |
| Crotalus-Bothrops | 1.46 | -1.08 | 3.99 | 0.50 |
| Sistrurus-Bothrops | 2.04 | -0.58 | 4.67 | 0.20 |
| Sistrurus-Crotalus | 0.58 | -0.24 | 1.41 | 0.29 |

**Table 13. (continued)**

| | | | | |
|---|---|---|---|---|
| PC4 Crotalinae | | | | |
| Bothriechis-Agkistrodon | -0.96 | -2.02 | 0.10 | 0.10 |
| Bothrops-Agkistrodon | -0.54 | -3.10 | 2.03 | 0.98 |
| Crotalus-Agkistrodon | -0.09 | -0.76 | 0.59 | 1.00 |
| Sistrurus-Agkistrodon | 1.13 | 0.18 | 2.09 | 0.01 |
| Bothrops-Bothriechis | 0.42 | -2.22 | 3.07 | 0.99 |
| Crotalus-Bothriechis | 0.87 | -0.07 | 1.81 | 0.08 |
| Sistrurus-Bothriechis | 2.09 | 0.93 | 3.25 | <0.01 |
| Crotalus-Bothrops | 0.45 | -2.06 | 2.96 | 0.99 |
| Sistrurus-Bothrops | 1.67 | -0.94 | 4.27 | 0.39 |
| Sistrurus-Crotalus | 1.22 | 0.40 | 2.04 | <0.01 |
| PC5 Crotalinae | | | | |
| Bothriechis-Agkistrodon | -0.33 | -1.53 | 0.87 | 0.94 |
| Bothrops-Agkistrodon | -0.38 | -3.29 | 2.52 | 1.00 |
| Crotalus-Agkistrodon | -0.10 | -0.86 | 0.66 | 1.00 |
| Sistrurus-Agkistrodon | 0.08 | -1.00 | 1.17 | 1.00 |
| Bothrops-Bothriechis | -0.06 | -3.05 | 2.94 | 1.00 |
| Crotalus-Bothriechis | 0.23 | -0.83 | 1.30 | 0.97 |
| Sistrurus-Bothriechis | 0.41 | -0.90 | 1.73 | 0.91 |
| Crotalus-Bothrops | 0.29 | -2.56 | 3.14 | 1.00 |
| Sistrurus-Bothrops | 0.47 | -2.48 | 3.42 | 0.99 |
| Sistrurus-Crotalus | 0.18 | -0.75 | 1.11 | 0.98 |
| PC6 Crotalinae | | | | |
| Bothriechis-Agkistrodon | -1.08 | -2.10 | -0.05 | 0.03 |
| Bothrops-Agkistrodon | -1.01 | -3.48 | 1.47 | 0.79 |
| Crotalus-Agkistrodon | 0.69 | 0.04 | 1.33 | 0.03 |
| Sistrurus-Agkistrodon | 0.30 | -0.62 | 1.22 | 0.89 |
| Bothrops-Bothriechis | 0.07 | -2.49 | 2.63 | 1.00 |
| Crotalus-Bothriechis | 1.76 | 0.85 | 2.67 | <0.01 |
| Sistrurus-Bothriechis | 1.38 | 0.26 | 2.50 | 0.01 |
| Crotalus-Bothrops | 1.69 | -0.74 | 4.12 | 0.31 |
| Sistrurus-Bothrops | 1.31 | -1.21 | 3.82 | 0.60 |
| Sistrurus-Crotalus | -0.39 | -1.18 | 0.41 | 0.66 |

**Table 14. Tukey's test results of primary foraging habitat for PCs 1-6 of the Crotalinae-only data.**

| | | | | |
|---|---|---|---|---|
| PC1 Crotalinae | | | | |
| Semiaquatic-Arboreal | -2.13 | -3.17 | -1.08 | <0.01 |
| Terrestrial-Arboreal | -1.24 | -2.04 | -0.44 | <0.01 |
| Terrestrial-Semiaquatic | 0.89 | 0.13 | 1.64 | 0.02 |
| PC2 Crotalinae | | | | |
| Semiaquatic-Arboreal | 0.07 | -1.09 | 1.24 | 0.99 |
| Terrestrial-Arboreal | -0.10 | -0.99 | 0.79 | 0.96 |
| Terrestrial-Semiaquatic | -0.18 | -1.02 | 0.67 | 0.87 |
| PC3 Crotalinae | | | | |
| Semiaquatic-Arboreal | -1.20 | -2.34 | -0.07 | 0.03 |
| Terrestrial-Arboreal | -0.64 | -1.50 | 0.22 | 0.19 |
| Terrestrial-Semiaquatic | 0.57 | -0.25 | 1.38 | 0.23 |
| PC4 Crotalinae | | | | |
| Semiaquatic-Arboreal | 0.13 | -0.94 | 1.19 | 0.96 |
| Terrestrial-Arboreal | 1.14 | 0.32 | 1.95 | <0.01 |
| Terrestrial-Semiaquatic | 1.01 | 0.24 | 1.78 | 0.01 |
| PC5 Crotalinae | | | | |
| Semiaquatic-Arboreal | 1.03 | -0.10 | 2.16 | 0.08 |
| Terrestrial-Arboreal | 0.18 | -0.68 | 1.05 | 0.87 |
| Terrestrial-Semiaquatic | -0.85 | -1.67 | -0.03 | 0.04 |
| PC6 Crotalinae | | | | |
| Semiaquatic-Arboreal | 1.12 | 0.08 | 2.17 | 0.03 |
| Terrestrial-Arboreal | 1.61 | 0.81 | 2.40 | <0.01 |
| Terrestrial-Semiaquatic | 0.48 | -0.27 | 1.24 | 0.28 |
